# Single-nucleus transcriptomes reveal functional and evolutionary properties of cell types in the Drosophila accessory gland

**DOI:** 10.1101/2021.06.13.448152

**Authors:** Alex C. Majane, Julie M. Cridland, David J. Begun

## Abstract

Many traits responsible for male reproduction evolve quickly, including gene expression phenotypes in germline and somatic male reproductive tissues. Rapid male evolution in polyandrous species is thought to be driven by competition among males for fertilizations and conflicts between male and female fitness interests that manifest in post-copulatory phenotypes. In Drosophila, seminal fluid proteins secreted by three major cell types of the male accessory gland and ejaculatory duct are required for female sperm storage and use, and influence female post-copulatory traits. Recent work has shown that these cell types have overlapping but distinct effects on female post-copulatory biology, yet relatively little is known about their evolutionary properties. Here we use single-nucleus RNA-Seq of the accessory gland and ejaculatory duct from *Drosophila melanogaster* and two closely related species to comprehensively describe the cell diversity of these tissues and their transcriptome evolution for the first time. We find that seminal fluid transcripts are strongly partitioned across the major cell types, and expression of many other genes additionally define each cell type. We also report previously undocumented diversity in main cells. Transcriptome divergence was found to be heterogeneous across cell types and lineages, revealing a complex evolutionary process. Furthermore, protein adaptation varied across cell types, with potential consequences for our understanding of selection on male post-copulatory traits.

**SIGNIFICANCE STATEMENT:** Rapid evolution of male traits may result from competition among males or antagonistic interactions between the sexes over control of reproduction. In animals with internal fertilization, interactions may occur in the female reproductive tract. Drosophila seminal fluid proteins, which are secreted by three major cell types of the male accessory gland and ejaculatory duct, are required for female sperm storage and use, and influence female behavior and physiology. These cell types have distinct effects on females, yet relatively little is known about their evolutionary properties. Here we characterize diversity and transcriptome evolution of seminal fluid-producing tissues at the cell level. These data reveal new functional properties of these cells and complex evolutionary patterns that vary across cell types and lineages.

## INTRODUCTION

Identifying and explaining variance in rates of evolution, which is commonly observed at all levels of biological organization, has been one of the great preoccupations of evolutionary biology. For example, some genes, proteins, and chromosomes evolve more quickly than others (White 1977; Kimura 1983), some traits evolve quickly in some lineages and slowly in others (Simpson 1944), and some traits evolve much more quickly in males than in females (Darwin 1871). This truism of evolutionary biology, that evolutionary rate variance is common and demands an explanation, extends to gene expression phenotypes, which tend to evolve relatively quickly in male reproductive tissues compared to most other tissues (reviewed in Ellegren and Parsch 2007). While the explanations proffered for faster expression evolution in male reproductive tissues often invoke rapidly changing selection pressures due to sexual selection or genomic conflicts, the biological processes driving rapid divergence of male reproductive tissues remain mostly unknown. Because the level of biological organization at which an evolutionary phenomenon is measured fundamentally shapes our understanding of evolutionary patterns, the level of analysis necessarily constrains the universe of testable hypotheses and the generation of new hypotheses. In the context of Drosophila gene expression, the phenomenology of rapid male-biased expression divergence has often been observed at the whole animal level or the organ level (focusing primarily on gonads) (Ranz et al. 2003; Meiklejohn et al. 2003; Assis, Zhou, and Bachtrog 2012; Whittle and Extavour 2019). In reality, most organs are a complex mixture of many cell types, which suggests that while organ analysis is preferable to whole-animal analyses, layers of biological causation and evolutionary inferences are still missed. Indeed, since gene products are produced in individual cells, one could reasonably argue that the cell is the natural level of organization for understanding expression variation and generating hypotheses relating expression variation to downstream phenotypes.

Theoretical concepts underlying the evolution of cell type diversity and the process of evolution in different cell types within a tissue are well-developed (reviewed in Arendt et al. 2016; Musser and Wagner 2015). Single-cell data in evolutionary contexts have generally been applied to distantly related taxa (Tosches et al. 2018; Hodge et al. 2019; Liang et al. 2018), typically focusing on cell type diversity (Sebé-Pedrós et al. 2018; La Manno et al. 2016; Colquitt et al. 2021; Feregrino and Tschopp 2021; J. Wang et al. 2021). Evolutionary analysis of different cell types across species, particularly on short time scales, has received less attention (Liang et al. 2018). In this study we use the polyandrous genus *Drosophila* as a model for evolution at the cellular level, with a focus on the tissues producing seminal fluid proteins (Sfps), which are transferred to females along with the sperm during mating. Many of these secreted proteins, which are produced in the accessory glands (AG) and the ejaculatory duct, induce a set of physiological and behavioral changes in females collectively referred to as the post-mating response (PMR; reviewed in Ravi Ram and Wolfner 2007). In *D. melanogaster*, the PMR includes increased rates of egg laying (Soller, Bownes, and Kubli 1999; Heifetz et al. 2000), decreased receptivity to re- mating (Liu and Kubli 2003), storage of sperm in specialized reproductive tract tissues (Neubaum and Wolfner 1999), elevated immune response (Peng, Zipperlen, and Kubli 2005), elevated feeding rates (Carvalho et al. 2006), increased activity rate, and decreased sleep (Isaac R. Elwyn et al. 2010). Genetic variation in Sfps may also play a role in the outcome of sperm competition (Clark et al. 1995; Fiumera, Dumont, and Clark 2005). Population genetic and comparative analyses of these proteins suggest they evolve unusually rapidly, often under the influence of directional selection (Begun et al. 2000; Tsaur, Ting, and Wu 1998; Aguadé 1999). These genes are frequently gained or lost during evolution (Wagstaff and Begun 2005, Muller *et al*. 2005), even on short timescales, (Begun and Lindfors 2005) and experimental evolution has shown that sexual conflict linked to PMR phenotypes may contribute to the rapid evolution of seminal fluid proteins (Hollis et al. 2019).

The *D. melanogaster* AG consists of two specialized, morphologically distinct, secretory epithelial cell types (Bairati 1968). Main cells are smaller, hexagonal and squamous, while secondary cells are much larger, spherical, project into the lumen of the gland, and contain extensive vacuole-like compartments (Bairati 1968; Prince et al. 2018). Main cells, which constitute the vast majority of AG cells, are necessary and sufficient to initiate the PMR (Kalb, DiBenedetto, and Wolfner 1993; Sitnik et al. 2016; Hopkins et al. 2019). Secondary cells, which are located at the distal tip of the gland, appear to contribute in part to the long term maintenance of the PMR, particularly with respect to remating phenotypes; females mated to males with deficient secondary cell secretions exhibit greater receptivity to remating (Leiblich et al. 2012; Hopkins et al. 2019). The ejaculatory duct consists of a single secretory epithelial cell type (Bairati 1968), contributing additional Sfps to the ejaculate (Sepil et al. 2018; Rexhepaj et al. 2003; Takemori and Yamamoto 2009). While the duct and its products contribute to the PMR (Rexhepaj et al. 2003; Saudan et al. 2002; Xue and Noll 2000), relatively little experimental work has been performed on this tissue. It is difficult to dissect individual phenotypic contributions of each cell type, however, given their apparent interdependence in production of the seminal fluid (Hopkins et al. 2019).

While genetic and gene expression studies of the AG have revealed evidence of both shared and distinct properties of these three major cell types, and much has been learned from genetic mutants knocking out (Kalb, DiBenedetto, and Wolfner 1993; Xue and Noll 2000; Minami et al. 2012; Gligorov et al. 2013; Sitnik et al. 2016) or suppressing secretions of (Leiblich et al. 2012; Corrigan et al. 2014; Hopkins et al. 2019) specific cell types in the AG, no study has directly investigated patterns of cell-type expression bias from transcriptome data. Here we carry out single-cell transcriptome analysis of the accessory gland and ejaculatory duct in three closely related Drosophila species, *D. melanogaster*, *D. simulans*, and *D. yakuba*. We use these data to:

1. reveal new biological attributes of the various cell types in the male somatic reproductive tract,
2. investigate rates of transcriptome divergence at the cellular level in multiple lineages, (3) determine the degree to which expression evolution is concerted or independent across cell types, and (4) investigate the connection between cell type-biased gene expression and adaptive protein divergence.

## METHODS

### Fly stocks and single-nucleus RNA sequencing

Additional details of all methods in this study can be found in <u>Supplementary Information.</u> We used the following sequenced stocks to generate AG and ejaculatory duct transcriptomes from 2- 3 day old virgin males for three *melanogaster* subgroup species: *D. melanogaster* RAL 517 (Mackay et al. 2012), *D. simulans w^501^*, and *D. yakuba* Tai18E2 (hereafter referred to as *mel*, *sim*, and *yak*) (Begun et al. 2007). Nuclei were isolated into a suspension using a modified version of Luciano Martelotto’s protocol (2019). FACS was used to purify single nuclei, and single-nucleus

RNA-Seq libraries were created using the 10X Genomics Chromium platform and Illumina sequencing.

### Bioinformatic assignment of species origin, RNA-Seq alignment, QC, and ortholog formatting

We parsed the 10X barcodes of raw reads and counted the number of unique molecular identifiers (UMIs) corresponding to each. We examined the distribution of UMI counts in descending rank order, using the ‘knee’ inflection point method (Macosko et al. 2015) to identify putative nuclei and empty barcodes. We used a custom alignment-based bioinformatic pipeline to assign species-of-origin to each nucleus. We aligned reads to the appropriate species genome (Flybase; *D. melanogaster* v6.33, *D. simulans* v2.02, *D. yakuba* v1.05) using STAR v2.7.5a (Dobin et al. 2013) with default parameters. We then filtered the set of nuclei according to alignment statistics to remove probable multiplets and nuclei with low sequencing depth. Next, we counted features from BAM files using HTSeq-count v0.12.3 (Anders, Pyl, and Huber 2015) with default parameters. For comparative analyses we created a set of 1-to-1-to-1 orthologs (11,481 genes) using the *D. melanogaster* ortholog table from Flybase (2020 version 2).

### Marker gene identification and differential expression among species

Single-nucleus gene expression analyses were performed in R v3.6.1 using Seurat v3.2.2 (Satija et al. 2015; Butler et al. 2018; Stuart et al. 2019) using two parallel approaches. We did an integrated analysis (Stuart et al. 2019) of the data across species using our *mel/sim/yak* 1-to-1- to-1 orthologues. We also performed an independent analysis of *mel* using all annotated genes to gain a fuller picture of gene expression variation among cell types. We identified marker genes using Seurat’s FindAllMarkers() method and assessed significance using a Wilcoxon Rank Sum test. We required marker genes to be expressed in at least 25% of focal cluster cells and set a minimal average log(fold-change) (logFC) requirement of 0.25. We filtered marker genes to those with Bonferroni-corrected p-values less than 0.05. To further investigate cell type specific expression bias of all Sfps, in addition to those strictly classified as marker genes, we did not impose minimum percent cells expressing and average logFC thresholds. We additionally identified markers distinguishing MC subpopulations from one another using the FindMarkers() method.

We used limma v3.42.2 (Ritchie et al. 2015) to infer differentially expressed (DE) genes for each cell type. We performed pairwise contrasts among the three species and classified genes as DE with an FDR of 5% (Benjamini and Hochberg 1995). Further details of the limma analysis can be found in our R scripts. To compare the rate of qualitative expression divergence across cell types, we calculated ratios of DE genes at various log2 fold change (logFC) cut-offs across the three cell types for each of the three pairwise species contrasts, and tested for differences in these ratios using a G-Test of goodness-of-fit (Sokal and Rohlf 2012). To test for differences in the magnitude of expression differences across cell types, we similarly compared distributions of absolute values of logFC using a Kruskal-Wallis test (Kruskal and Wallis 1952). Finally, we examined overall expression correlations between species within cell types by calculating average expression per gene and Pearson correlation coefficients.

To examine the relative level of concerted vs independent gene expression evolution across cell types, we subset the data to the set of DE genes exhibiting a logFC greater than one in at least one cell type-specific pairwise species contrast. We then calculated pairwise Pearson correlation coefficients of logFC across cell types within each of the three pairwise species contrasts. We permuted logFC values across genes 10,000 times to obtain a distribution of Pearson correlation coefficients under the null expectation of entirely cell type independent change within our set of DE genes.

### Population genetic inference of adaptive protein divergence of marker genes

To investigate potential differences in the prevalence of adaptive protein evolution across cell types, we used existing population data (2019) from *D. melanogaster* (Lack et al. 2015) with *D. simulans* as the outgroup. We considered two summaries of the role of adaptation in protein divergence: the proportion of marker genes with *α* > 0, and the distribution of *α* values amongst those genes with *α* > 0. The proportions of positive *α* values were compared using Fisher’s exact test, with post-hoc pairwise tests between cell types. The distributions of positive *α* values were visualized in ggplot2 v3.3.3 (Wickham 2016), and compared using a Kruskal-Wallis test with post-hoc pairwise Wilcoxon tests.

To determine whether the prevalence of positive selection in AG-expressed genes correlates with differential gene expression, we intersected *α* values with DE genes. We selected the set of all genes expressed in the AG and filtered out genes expressed at a level lower than the lowest-expressed DE gene, to account for power to detect DE. We tested whether DE genes and non-DE genes had different likelihoods of showing positive selection by comparing the fraction of positive *α* values in each class of genes using a G-test. We tested whether the fraction of sites with evidence of positive selection differed among classes of genes by comparing distributions of positive *α* values using a Kruskal-Wallis test.

De novo transcriptome assembly and identification of unannotated *D. melanogaster* transcripts

For de novo transcriptome assembly, we trimmed reads with TrimGalore! v0.6.5 (https://github.com/FelixKrueger/TrimGalore) and used Trinity v2.11.0 (Grabherr et al. 2011) to create the assembly. We augmented our assembly with two additional bulk RNA-Seq datasets (Leader et al. 2018; Immarigeon et al. 2021)—see <u>Supplemental Methods</u>. We quantified abundances of de novo-assembled transcripts in each cell type population with Salmon v0.12.0 (Patro et al. 2017). We used a BLAST-based strategy (Camacho et al. 2009) to identify candidate unannotated transcripts in *D. melanogaster*. We then took the set of transcripts that had at least one BLAST hit to the *mel* reference sequence but no BLAST hits to *mel* gene annotations. We also used the Ensembl Metazoa BLAST search tool to verify that these candidate transcripts do not overlap with any annotated features (Howe et al. 2020). We filtered out very lowly expressed transcripts using counts from Salmon. We created a GTF file based on the BLAST coordinates of our candidate transcripts, and aligned our raw sequencing reads with STAR, performed feature counting with HTSeq, and removed ambient RNA using SoupX, as described earlier for transcriptome-wide analysis.

We used Ensembl Metazoa BLAST and the *mel* genome browser (Howe et al. 2020) to identify transcript coordinates, strand, and neighboring annotated genes. For cell type specific analysis of unannotated-transcript expression we added transcript counts to the broader *mel* dataset, post-hoc. We used Seurat’s FindAllMarkers() method to identify cell type expression bias. and significance was assessed using a Wilcoxon Rank Sum test with Bonferroni multiple test correction. We assessed coding potential with CPAT v2.0.0 (L. Wang et al. 2013). To identify potential open reading frames (ORFs), we used the getorf function in the EMBOSS software package (http://emboss.sourceforge.net/apps/cvs/emboss/apps/getorf.html). We attempted to characterize these potential ORFs further using Ensembl Metazoa Protein BLAST (Howe et al. 2020) to the database of all *mel* proteins, NCBI’s Conserved Domain Database search tool (Lu et al. 2020), and SignalP v5.0 (Almagro Armenteros et al. 2019) to identify putative signal sequences.

## RESULTS

### Overview of single-nucleus RNA-Seq data

Following QC filtering to remove putative multiplets, we obtained a total of 4271 nuclei for single-cell analysis. The dataset comprised 1167 *mel*, 2116 *sim*, and 994 *yak* nuclei. While the overrepresentation of *sim* nuclei could be an artifact, given that tissue was pooled from nearly equal numbers of glands from each species prior to isolation of nuclei, it seems plausible that this difference results from divergence in cell number. Median counts per nucleus for *D. melanogaster*, *simulans*, and *yakuba* (hereafter referred to as *mel, sim, and yak*), were 1022, 1262.5, and 741.5, respectively, exhibiting the same species rank order as nuclei abundance, consistent with the idea of species differences in levels of seminal fluid production. We used k-nearest-neighbor based clustering with UMAP visualization to identify three primary clusters of cells in both the *mel* and three-species dataset (Fig. 1*A-C*). We then used marker gene identification along with the relative sizes of clusters to assign cell type identity to clusters, identifying MC, SC, and EDC. MC were identified as the cluster with the largest number of cells, and on the basis of markers *Sex Peptide* (*SP*) (Styger 1992), *Acp36DE* (Wolfner et al. 1997), and *Acp95EF* (DiBenedetto, Harada, and Wolfner 1990; Kalb, DiBenedetto, and Wolfner 1993) (Fig. 1*D*). SC and EDC were classified as relatively smaller clusters. SC were identified by expression of *lectin-46Ca* (*CG1652*) and *lectin-46Cb* (*CG1656*) (Maeda et al. 2018), *abd-A (Maeda et al. 2018)* and additionally by *iab-8* (Maeda et al. 2018) in the *mel*-only dataset (*iab-8* orthologues are not annotated in *sim* or *yak*) (Fig. 1*D*). EDC were identified by expression of *vvl* (Junell et al. 2010) and *Dup99B* (Rexhepaj et al. 2003) (Fig. 1*D*). We additionally used *Abd-B* to characterize both SC and EDC (Maeda et al. 2018; Gligorov et al. 2013). In the *mel* dataset, we identified 1056 MC, 51 SC, and 60 EDC, with 6444, 2596, and 3445 expressed genes, respectively. In the three species dataset, we identified a total of 3629 MC, 139 SC, and 509 EDC, with 6978, 3573, and 5978 expressed orthologous genes, respectively. While our results revealed no evidence of transcriptional heterogeneity among SC or EDC, we observed strong evidence of MC subpopulations (see <u>Transcriptome heterogeneity among main cells</u> below). For downstream analyses, we merged these sub-clusters into a single MC cluster.

**Figure 1.**
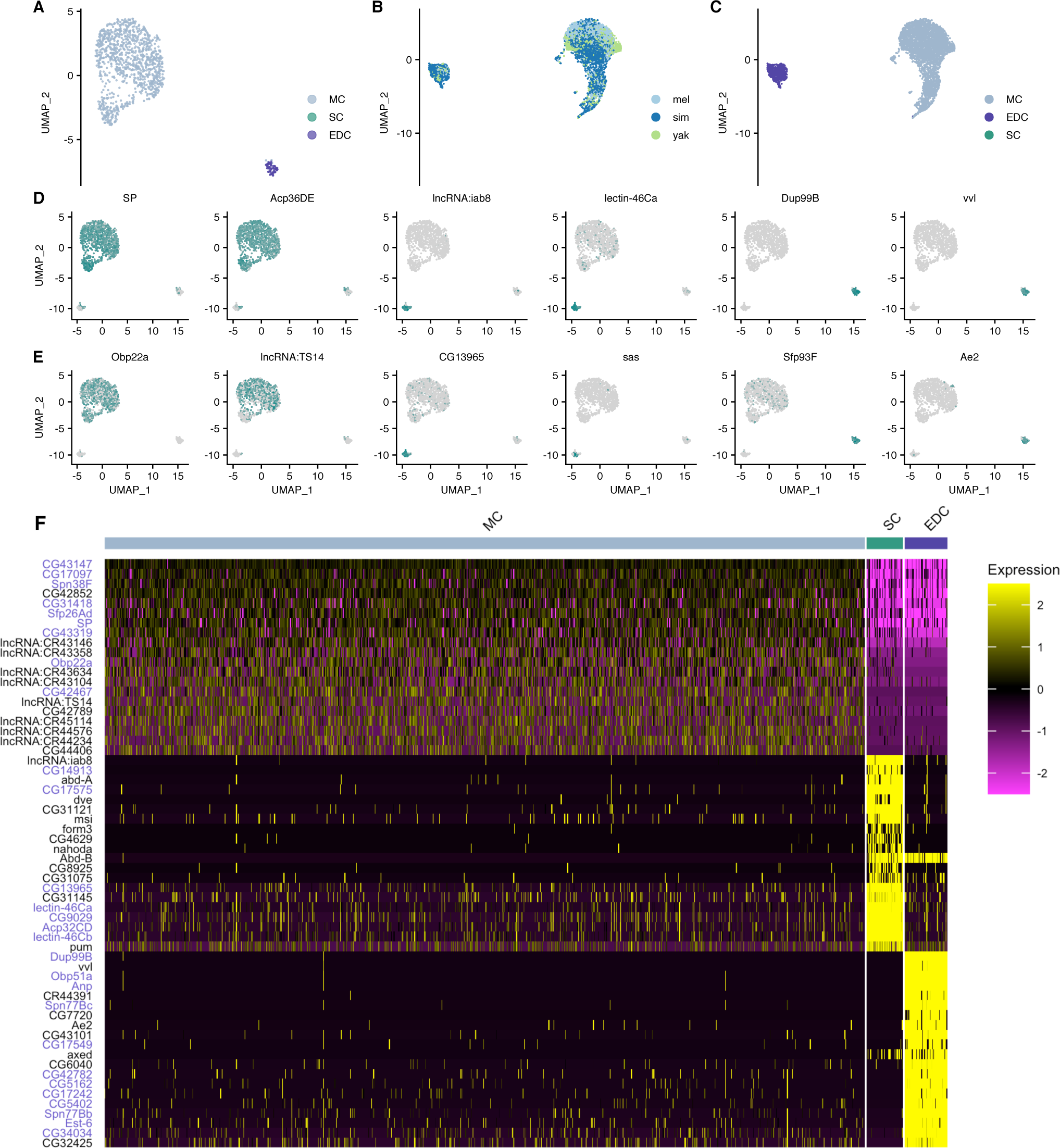
(*A*) UMAP showing clustering of *mel* single-nucleus transcriptomes into three major cell types: main cells (MC), secondary cells (SC), and ejaculatory duct cells (EDC). (*B*) Nuclei from three species cluster concordantly, (*C*) into the same three major cell types. Example marker genes in *mel*, with expression indicated in teal: (*D*) well known markers, (*E*) novel markers. (*F*) Heatmap showing scaled expression of the top 20 markers of each cell type. Seminal fluid proteins (Sfps) are highlighted in blue text.

### Cell type transcriptomes in the *Drosophila melanogaster* accessory gland

Thresholding marker genes as expressed in at least 25% of cells in the focal cell type and minimum logFC = 0.25, we identified 540 *mel* marker genes (Fig. 1*F*, Dataset S1). Of these, 123 are annotated Sfps identified from proteomic studies of the male ejaculate (Findlay, MacCoss, and Swanson 2009; Sepil et al. 2018), consistent with previous results that the majority of Sfps showing cell-type bias are expressed in MC (Swanson et al. 2001; Wolfner et al. 1997; Kalb, DiBenedetto, and Wolfner 1993). Of the 123 Sfp markers, 92 (75%) are MC markers, 10 (8%) are SC markers, and 21 (17%) are EDC markers. Marker Sfps for SC and EDC are summarized in Table S1. Among the 214 total MC markers, 43% are Sfps. Among the 82 SC markers, only 12% are Sfps, and among the 262 EDC markers, 8% are Sfps. MC marker genes are significantly enriched for Sfps relative to both SC and EDC (pairwise G-tests, p < 0.001), while SC and EDC are not significantly different (p = 0.26). Thus, in contrast to MC, the distinct natures of SC and EDC transcriptomes are not driven primarily by Sfp expression.

To investigate cell type expression bias for all Sfps in addition to that of marker genes, we calculated for each of 264 *mel* Sfps the log2(average expression) for the focal cell type and the average logFC vs. all other cells. Among the 219 Sfps detected in the data (Dataset S2), 158 (72%) show greatest expression in MC, 24 (11%) show greatest expression in SC, and 37 (17%) show greatest expression in EDC. Expressed Sfps generally exhibit cell-type expression bias, with relatively few Sfps showing comparable expression across types (Fig. 2). Highly MC-biased Sfps tend to also show expression in SC, though at a substantially lower level. Even among non-marker Sfps we observe a trend towards greater MC expression than SC expression (Fig. 2*A*, SC expression vs. MC expression gives a slope = 0.73, r^2^ = 0.82). EDC vs. MC comparison for non-marker Sfps exhibits a similar pattern (Fig. 2*B*, slope = 0.79, r^2^ = 0.75). Comparing SC vs. EDC suggests a relatively more even spread of expression across these cell types, with some bias towards SC (Fig. 2*C*; slope = 0.69, r^2^ = 0.59). Among the 98 non-marker Sfps, 67 (68%) show highest expression in MC, while 13 (13%) have highest expression in SC, and 18 (18%) have highest expression in EDC. Additionally, the distribution of average logFC of Sfps in MC. vs all other cells skews significantly greater than SC vs all others and EDC vs all others, respectively (Fig. 2*D*). The median logFC of MC vs all other cells is 0.75, while SC vs all others is -0.85, and EDC vs all others is -0.89.

**Figure 2.**
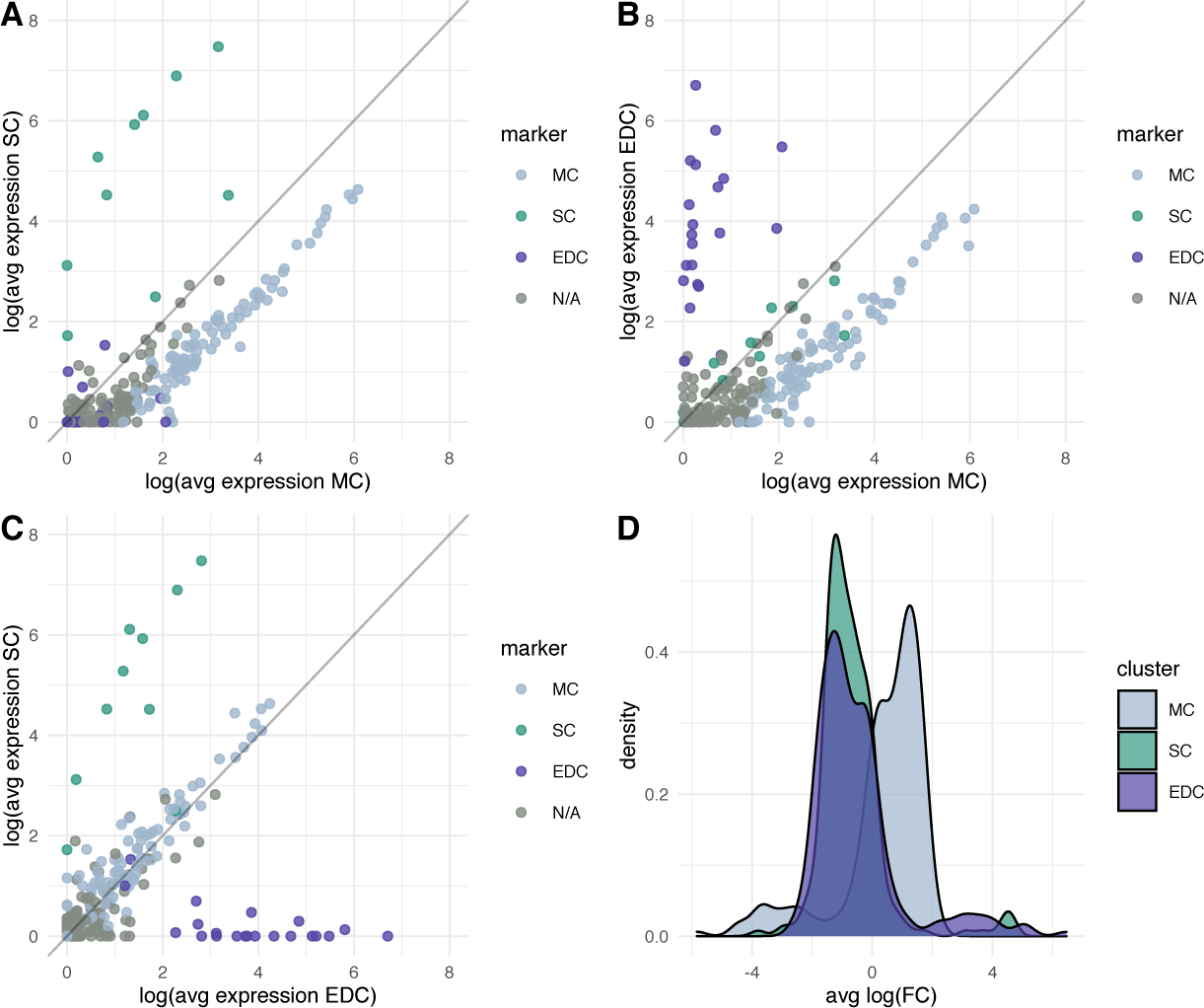
Expression of Sfps tends to be highly cell type-specific. (*A-C*) Expression levels of Sfps compared among cell types show a general pattern of MC enrichment and cell type bias. (*D*) The average log(fold-change) of expression between each cell type and the other two shows that most SFPs are most highly expressed in MC, with few Sfps showing highest expression in SC or EDC.

Using all annotated *mel* genes, marker genes for each *mel* cell type reveal both expected and novel markers (Dataset S1). In MC we identify many expected Sfps including *SP* (Fig 1*D*), *Acp36DE*, *Acp26Aa*, and *Acp95EF,* and relatively uncharacterized Sfps including *Obp22a* (Fig. 2*E*). The top non-Sfp markers of MC are generally functionally uncharacterized: *CG42852*, *CG43254*, *CG42481*, *CG43392*, *lncRNA:CR43146*, *lncRNA:CR45013*, *CG34041, lncRNA:TS14* (Fig. 1*E*), and the genes *CG44388* and *lncRNA:CR44389*, which are neighbors. Despite its annotation as a lncRNA, *CR44389* possesses a 41 amino acid ORF strongly predicted to have a signal sequence, suggesting it could be a secreted protein. *Ugt50B3,* a UDP-glycosyltransferase, is another strong marker of MC.

Among the 10 Sfps identified as SC markers (Table S1), three were previously known to be SC-specific: *Acp32CD*, *lectin-46Ca* and *lectin-46Cb* (Maeda et al. 2018). Previous work with MC-null mutants identified *Acp32CD* as expressed in SC (Swanson et al. 2001), and here we additionally show that it exhibits very low expression in MC. The Sfps *CG17575, CG3349, CG9029*, *CG13695* (Fig. 1*E*), and *mfas* have also been previously identified as SC-expressed (Gligorov et al. 2013; Sitnik et al. 2016; Immarigeon et al. 2021). Here we show that these Sfps show very low expression in MC and EDC. We also identify the Sfp *Pgant9* as a novel SC marker. We additionally recovered expected non-Sfp markers: *lncRNA:iab8*, *abd-A (Maeda et al. 2018)*, *abd-B*, and *defective proventriculus* (*dve*) (Minami et al. 2012). We also identify non-Sfp SC markers *stranded-at-second* (*sas*) (Fig. 1*E*), *musashi* (*msi*), *form3*, *nahoda*, *CG31121*, *CG4629*, and *CG46430*. Additionally, we discovered that the unannotated transcript *DN2695* (see <u>Identification of unannotated candidate genes in the AG</u>, Table S4, Fig. S3) is a strong SC marker.

We identified 21 Sfp EDC markers (Table S1). Of these, 12 had previously been identified as EDC-enriched in proteomics studies: *Dup99B*, *Obp51a*, *Spn77Bc*, *Spn77Bb*, *Est-6*, *Gld*, *CG18258*, *CG5162*, *CG17242*, *CG5402, CG34034,* and *CG31704* (Takemori and Yamamoto 2009; Sepil et al. 2018; Samakovlis et al. 1991; Cavener 1985; Saudan et al. 2002). The remainder have not been previously identified as EDC-specific Sfps: *Treh*, *betaggt-I*, *Sfp93F* (Fig. 1*E*)*, trx*, *NT5E-2, Anp, CG17549*, *CG42782*, and *CG15394*. *CG42782* was previously identified as a likely mating plug protein gene, consistent with origin in the ejaculatory duct or ejaculatory bulb (Avila et al. 2015). We also identified expected non-Sfps, *ventral veins lacking* (*vvl*) (Junell et al. 2010) and *Abd-B (Gligorov et al. 2013)*. Novel EDC markers are *anion exchanger 2* (*Ae2*) (Fig. 1*E*), *axundead* (*axed*), *single-minded* (*sim*), *CG7720*, *CG43101*, *CG7342*, and *CG13012,* and *CR44391*. *CR44391* is annotated as a pseudogene created by a tandem duplication of *CG11400* (an EDC-biased gene), however, it has a homologous ORF with a strongly predicted signal sequence.

Transcriptome heterogeneity among *D. melanogaster* main cell subpopulations

During initial analysis we discovered an apparent subcluster of main cells characterized by unique SNN clusters at *k* = 4 and clear separation in UMAP space (Fig. 3*A*). Of a total 1057 MC, 942 are in subcluster one (MCsp1) and 115 are in subcluster two (MCsp2). 349 significant markers (Bonferroni-corrected p < 0.05) distinguish these subclusters (Dataset S3). In all three species, these subclusters are apparent and appear in roughly equal proportions (Fig. S1, Dataset S4), strongly supporting the idea that they reflect a conserved, regulated phenomenon. Of the 349 markers distinguishing the MC subclusters, 32 are Sfps, all of which are MC markers and expressed in both subpopulations (Fig. 3*B*). Twenty-four show higher expression in MCsp2, while just eight show higher expression in MCsp1 (Dataset S3). Non-Sfps show the opposite pattern, with 108 genes showing increased expression in MCsp1, and 218 genes with higher expression in MCsp2 (Dataset S3). The most enriched non-Sfp genes for each subpopulation are shown in Fig. 3*D*. Genes significantly enriched in MCsp2 include 57 of the proteins comprising the large and small ribosomal subunits, along with *Eukaryotic Translation Elongation Factor 2* (*eEF2*), additional translation elongation factors *eEF5*, *eEF1δ*, and *eEF1α1*, and translation initiation factors *eIF3a*, *eIF3b*, and *eIF3c*. Notably, MCsp1 has a lower level of RNA counts per nucleus than MCsp1, with 832 vs. 1248 median counts (Fig. 3*C*, Wilcoxon rank sum test, p < 0.001). Together with the quantitatively greater level of Sfp expression, these markers suggest a higher level of transcription accompanied by greater expression of translational machinery. Markers of MCsp1 include *Golgi microtubule-associated protein* (*Gmap*), *easily shocked* (*eas*), *taiman* (*tai*), and lncRNAs including *roX1*, *Hsrω*, *CR43104*, *CR43146*, and *CR45114* (Fig. 3*D*). *roX1*, one of the strongest markers of MCsp1, plays a central role in dosage compensation (Mukherjee and Beermann 1965; Meller et al. 1997; Hallacli et al. 2012). We investigated patterns of broadly expressed genes using the methods of Mahadevaraju *et al*. (2021), but found no evidence of correlations between *roX1* abundance and X-to-autosome expression, or variation in X-to-autosome expression among subclusters or cell types. Finally, we observe evidence of finer functional divisions within MC (Fig. S1; Dataset S5) that deserve further investigation.

**Figure 3.**
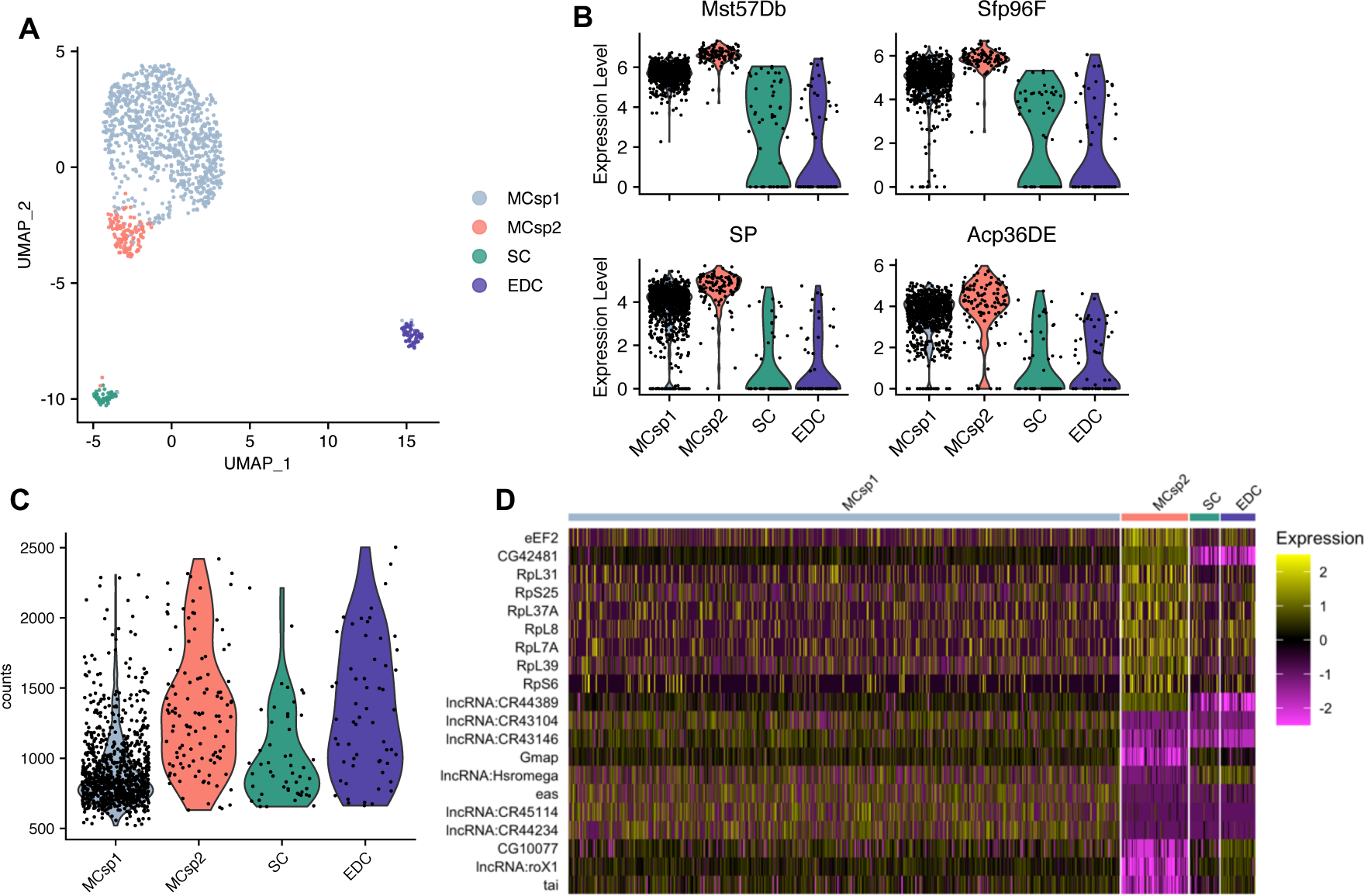
Transcriptome heterogeneity among subpopulations of MC in *mel*. (*A*) Subpopulations of MC are apparent in both UMAP space and SNN clustering with *k* = 4. (*B*) Examples of MC marker Sfps with greater expression in MCsp2. (*C*) MCsp1 has a significantly lower level of RNA counts per cell than MCsp2 or EDC (Kruskal-Wallis test and Wilcoxon rank sum tests, p < 0.001), but not SC (Wilcoxon rank sum test, p > 0.05). There is no significant difference between MCsp2 and EDC (Wilcoxon rank sum test, p > 0.05). (*D*) Heatmap showing scaled expression of the top 10 non-Sfp markers for each subpopulation, suggesting enrichment of translational machinery in MCsp2.

### Cell type-specific differential gene expression across species

We used our integrated three-species dataset to characterize differential gene expression (DE) across species. UMAP visualization reveals strongly concordant clustering of cell types across species (Fig. 1*B*,*C*). The top 12 DE genes for each cell type are summarized in Table S2, and expression of DE genes in all cell types can be found in Dataset S9. We found 131 genes that are DE (logFC > 1) in at least one pairwise species contrast among MC (Dataset S6), of which 31 (23%) are Sfps. Among SC we found 106 DE genes (Dataset S7), of which 21 (20%) are Sfps, while in EDC we found 221 (Dataset S8), of which only 29 (13%) are Sfps. The percentage of expressed genes that are DE for each species contrast and cell type (Fig. 4*A*) is significantly heterogeneous (G-test, p < 0.001, Table S3). Notably, EDC show a consistently greater fraction of DE genes than MC and SC for each species comparison, except for *sim*-*yak* EDC vs SC. The fraction of DE genes does not differ between MC and SC for any species contrasts. The fraction of DE genes in different cell types tends not to vary significantly over species contrasts, except for EDC, where the *mel-yak* fraction is significantly greater than *mel-sim,* but not significantly different from *sim-yak*. To determine the magnitude of DE among the genes that most distinguish each cell type we asked how many marker genes were DE in each cell type. In MC, 73 of 309 markers (24%) show DE, in SC, 25 of 121 markers (21%) show DE, and in EDC, 123 of 255 markers show DE (33%). EDC markers are significantly more likely to show DE than MC or SC (pairwise Fisher’s Exact Tests, p < 0.001), while MC and SC are not significantly different (p = 0.7). Together, the data suggest an elevated level of DE for EDC relative to MC and SC, and an effect of lineage on DE in EDC; the *mel*-*yak* EDC contrast has significantly more DE genes than *sim*-*yak*, suggesting that DE genes accumulated faster in the *mel* EDC than the *sim* EDC. These conclusions are robust to different logFC cutoffs (Fig S2*A-D*). There is a trend towards elevated MC enrichment compared to SC at particularly high and low cutoffs, however these differences are not statistically significant (Wilcoxon rank sum tests, p > 0.05). We found no evidence of differences in the magnitude of DE across cell types and lineages; distributions of logFC among DE genes are not significantly different (Fig. S2*E-F*).

**Figure 4.**
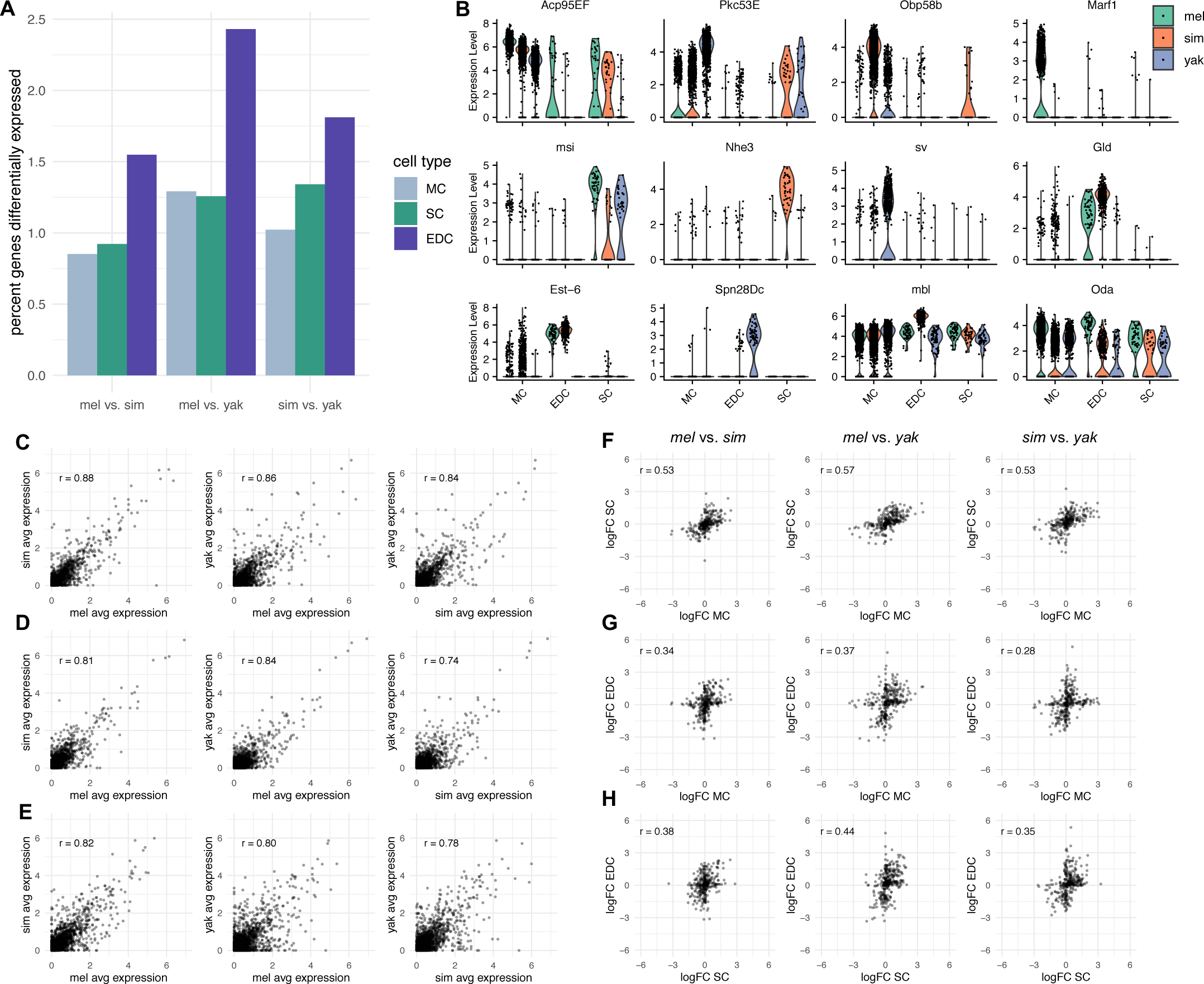
(*A*) Percentage of expressed genes DE by cell type and species contrast. For significance values see Table S3. (*B*) Examples of differential expression detected in this study. (*C-E*) Transcriptome-wide expression correlations show cell type- and species-specific patterns of divergence. (*C*) MC, (*D*) SC, (*E*) EDC. (*F-H*) Contrast correlations of logFC of DE genes reveal differences in the level of concerted vs independent DE events among cell type- and species-contrasts. (*F*) MC vs SC, (*G*) MC vs EDC, (*H*) SC vs EDC. Columns indicate each of three species-contrasts.

We used Pearson correlations of expression among all genes in species contrasts to investigate overall levels of transcriptome-wide divergence. MC have the greatest overall correlations (Fig. 4*C*; *rMCmel*-*sim* = 0.88, *rMCmel*-*yak* = 0.86, and *rMCsim*-yak = 0.84). Pearson correlations for SC and EDC are lower overall (Fig. 4*D-E*; *rSCmel*-*sim* = 0.81, *rSCmel*-*yak* = 0.84, *rSCsim*-yak = 0.74; *rEDCmel*-*sim* = 0.82, *rEDCmel*-*yak* = 0.80, *rEDCsim*-yak = 0.78). The data suggest an overall slower rate of expression evolution in MC than SC and EDC. Furthermore, the heterogeneous correlations for SC and EDC across species pairs suggest lineage by cell type interactions on rates of transcriptome evolution.

Examples of DE genes among MC include the Sfp *Acp95EF*, which has highest expression in *mel*, lower expression in *sim*, and lowest expression in *yak* (Fig 4*B*). The non-Sfp *Protein C kinase 53 E* (*Pkc53E*) shows the opposite pattern, expressed at the highest level in *yak*, and lowest in *mel* (Fig 4*B*). The transcription factor *shaven* (*sv*) is lowly expressed in *mel* and *sim*, but much more highly expressed in *yak*. *Meiosis regulator and mRNA stability factor 1* (*Marf1*) has low expression in *sim* and *yak*, but high expression and MC bias in *mel* (Fig 4*B*). Our previous work using bulk-tissue RNA-Seq described *Marf1* as gaining “neomorphic” expression in the *mel* AG (Cridland et al. 2020). *Odorant-binding protein 58b* (*Obp58b*) is highly expressed in *sim*, expressed moderately in *yak*, and rather lowly expressed in *mel* (Fig 4*B*). Findlay *et al*. (2009) detected peptides corresponding to *Obp58b* in a proteomic screen of *sim* seminal fluid, but did not detect any corresponding peptides in *mel* or *yak* seminal fluid. Taken together, these results suggest *Obp58b* is a MC-expressed Sfp in *sim*, but does not have a role as an Sfp in *mel*. The status of *Obp58b* in *yak* is less clear.

Examples of DE genes among SC include *Na+/H+ hydrogen exchanger 3* (*Nhe3*), with high expression in *sim* and near-zero expression in *mel* and *yak* (Fig 4*B*), consistent with *sim* gain-of-expression. Another top DE gene is *Peroxin 19* (*Pex19*), which exhibits what is likely gain-of-expression in *mel* SC and near-zero expression in *sim* and *yak* (Fig 4*B*). *Sex Peptide Receptor* (*SPR*), which is responsible for interactions with *SP* in the female reproductive tract (Yapici et al. 2008), is expressed in *yak* SC, but not in *sim* or *mel*, or in MC or EDC (Fig 4*B*). *musashi* (*msi*) is expressed most highly in *mel*, an intermediate level in *yak,* and at a relatively low level in *sim* (Fig 4*B*). In general, we observed little DE among SC-biased Sfps. While 24 Spfs exhibit SC DE, 22 of these are MC markers, with significantly lower expression in SC than MC. Two exceptions are *midline fascilin* (*mfas*) and *CG3349* (Dataset S7).

Examples of DE genes among EDC include the Sfp *Esterase 6* (*Est-6*), which is highly expressed in *mel* and *sim*, and much more lowly expressed in *yak* (Fig 4*B*). *Est-6* expression in the ejaculatory duct is specific to *mel*, *sim*, and *D. sechellia,* and notably absent in the rest of the *melanogaster* subgroup, including *yak (Richmond et al. 1990)*. *Serpin 28Dc* (*Spn28Dc*) has *yak*- specific EDC expression, with no expression in other cell types or species (Fig 4*B*). *Glucose dehydrogenase* (*Gld*) has a high level of expression in *sim*, a lower level in *mel*, and near-zero expression in *yak* (Fig 4*B*). This same species-specific pattern was previously observed in enzymatic GLD assays (Cavener 1985), suggesting that variation in GLD abundance in the ejaculatory duct is ultimately controlled at the transcriptional level.

To determine the ratio of markers to non-markers among DE genes, we used singlet markers (characterizing just one cell type) called independently for each species to filter our list of DE genes. We found 61% of DE genes were markers specific to a particular cell type. However, we find large differences in this ratio among cell types; 73% of genes differentially expressed in MC are MC markers, 75% of genes differentially expressed in EDC are EDC markers, while just 19% of DE genes in SC are SC markers. Thus, much DE is associated with cell-type biased expression for MC and EDC but not for SC. For example, *muscleblind* (*mbl*) exhibits high EDC expression in *sim* relative to both *mel* and *yak*, while showing no DE in MC or SC, despite high expression in these cell types (Fig 4*B*). Alternatively, DE may be correlated in the same direction across multiple cell types. For example, *Ornithine decarboxylase antizyme* (*Oda*) is broadly expressed and shows the same pattern of increased *mel* expression in each cell type (Fig 4*B*).

To investigate the degree of concerted vs. independent expression evolution across cell types we calculated pairwise Pearson correlation coefficients (*r*) of logFC of DE genes for each cell type for each of the three species contrasts. We find *r* ranges between 0.28 and 0.57 for each comparison (Fig. 4*F-H*). MC and SC have the highest correlations; *rmel*-*sim* = 0.53, *rmel*-*yak* = 0.57, and *rsim*-yak = 0.53. SC and EDC are less correlated; *rmel*-*sim* = 0.38, *rmel*-*yak* = 0.44, and *rsim*-yak = 0.35.

MC and EDC have the lowest correlations: *rmel*-*sim* = 0.34, *rmel*-*yak* = 0.37, and *rsim*-yak = 0.28. To determine the expected distribution of *r* under a null model of cell type-independent evolution, we permuted logFC 10,000 times and calculated values of *r,* as before. The 99th percentile of permuted *r* (0.123 to 0.133) was much lower than each observed *r*, supporting the hypothesis of correlated transcriptome divergence across cell types. Nevertheless, a gene is unlikely to pass our logFC ≥ 1 threshold for DE in multiple cell types; of 362 DE genes, 282 (78%) appear in a single cell type, 51 (14%) appear in two, and just 25 (7%) appear in all three cell types. This pattern is reflected in plots of logFC across cell types, with relatively few points falling near the line *x* = *y* (Fig. 4*F-H*). Thus, while the overall directionality of DE is similar among cell types, the largest interspecific expression differences tend to be limited to one cell-type.

### Protein sequence evolution in *melanogaster*

To investigate the evidence for protein adaptation among marker genes of each cell type, we used the McDonald-Kreitman test summary statistic *α*. A positive value of *α* suggests a history of directional selection. Among positive values, *α* provides an estimate of the proportion of amino acid differences between *mel and* sim attributable to directional selection. We obtained estimates of *α* for 561 of 691 marker genes (called from joint analysis of *mel*, *sim*, and *yak*), of which 265 (47%) were positive. The proportion of MC markers with positive *α* (61%) is significantly greater than SC (41%) or EDC (40%) (Fig. 5*A*; pairwise Fisher’s exact tests, p = 0.002, p < 0.001, respectively), suggesting that compared to SC and EDC, MC markers are more likely to have a history of adaptive protein divergence. Median values among positive *α* for SC, MC, and EDC are 0.30, 0.57, and 0.52, respectively (Kruskal-Wallis test, p = 0.01), with SC being significantly smaller than MC and EDC (pairwise Wilcoxon rank sum tests, p = 0.008). Overall, it appears MC-biased genes exhibit the greatest adaptive protein divergence and SC-biased genes the least. Given the enrichment for Sfp expression in MC we wanted to investigate whether this pattern of MC protein adaptation is driven by Sfp variation or is a general property of this cell type. Among marker genes, 91 of 125 Sfps (73%) have positive *α* values, while 177 of 437 non-Sfps (41%) have positive *α* values, a significant enrichment among Sfps (Fig. 5*B*; Fisher’s exact test, p < 0.001). However, medians of positive *α* values are not significantly different for Sfps vs. non-Sfps (Kruskal-Wallis test, p = 0.17).

**Figure 5.**
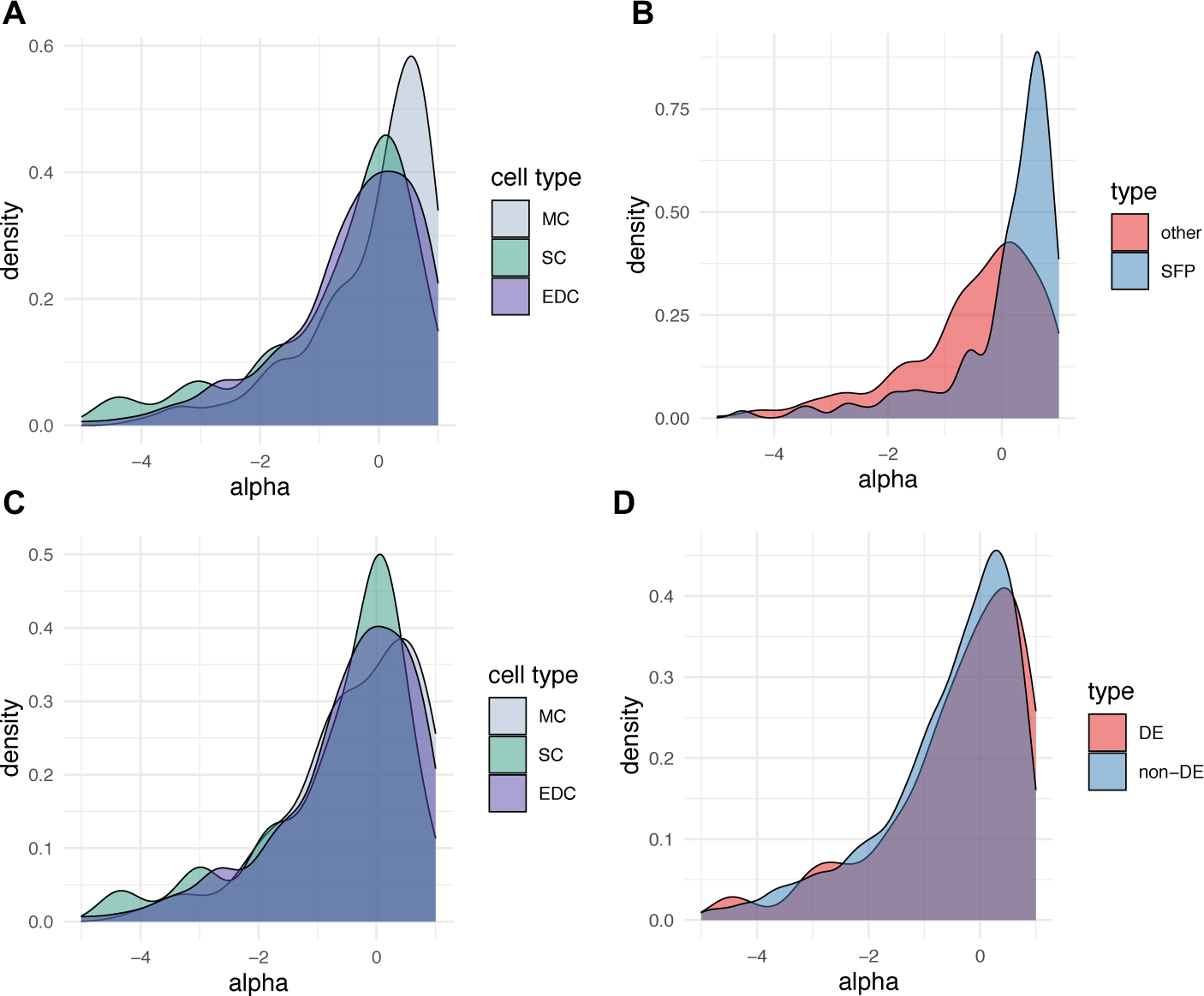
Distributions of *α* for marker genes. (*A*) *α* values by cell type show that MC markers are significantly greater than SC or EDC (Kruskal-Wallis test, p = 0.001). (*B*) Sfp markers are much more likely to have positive *α* than non-Sfps (Fisher test, p < 0.001). (*C*) Removing Sfps from the data shifts the distribution of MC *α* lower. MC and EDC are no longer significantly different, but SC is significantly less than MC and EDC (Kruskal-Wallis test, p = 0.006). (*D*) Genes that are DE between *mel* and *sim* have a modest but significantly greater *α* than non-DE markers (Kruskal-Wallis test, p < 0.001).

Among non-Sfp markers there is no significant difference in the proportion of positive vs. negative *α* among cell types (Fig. 5*C*; Fisher’s exact test, p = 0.41). However, non-Sfps show significant differences in distributions of positive *α*, with median *α* of 0.25 in SC, 0.54 in MC, and 0.51 in EDC (Kruskal-Wallis test, p = 0.006). Both MC and EDC are significantly greater than SC (pairwise Wilcoxon rank sum tests, p = 0.010 and p = 0.006, respectively). Thus, while the unequal distribution of Sfps among marker genes in different cell types accounts for some of the observed cell-type heterogeneity in the proportion of markers showing excess protein divergence, the reduced effect of directional selection on protein divergence in SC-biased genes remains apparent as a general phenomenon.

To investigate whether genes that are differentially expressed between *mel* and *sim* are also enriched for adaptive protein divergence for the *mel-sim* species pair, we compared *α* for genes that were DE vs. non-DE. While the proportion of DE vs. non-DE genes exhibiting *α* > 0 (42.5% and 38.7%, respectively) were not significantly different (Fig. 5*D*; G-test, p = 0.37), the median positive *α* value for DE genes, 0.59, was significantly greater than median positive *α* for non-DE genes 0.46 (Kruskal-Wallis test, p < 0.001). Thus, expression divergence appears to be more strongly correlated with the proportion of protein divergence explained by selection than with the probability of a protein having elevated levels of fixed nonsynonymous substitutions.

### Identification of unannotated genes expressed in the AG

Following stringent filtering (see <u>Supplemental Methods</u>), we identified 11 previously unannotated, single-exon genes (Table S4; Dataset S10). Transcript assemblies of FlyAtlas2 data were used to improve our annotation for seven of these candidates. Since *DN100097* and *DN2695* are SC-limited in expression (Fig. S3*A*), we used RNA-Seq data from FACS-sorted secondary cells (Immarigeon et al. 2021) to further improve our annotations. The median transcript length is 630 bp (range = 352 to 3,102 bp). None of these genes overlap annotated features in the *mel* genome. Among these genes, four show strong MC bias, two are SC-biased, and two are EDC-biased. In general, these candidates are expressed at a relatively high level compared to expressed annotated genes, but a relatively low level compared to marker genes (Fig. S3*B*). The two notable exceptions to this trend are *DN2695* in SC, and *DN818* in MC, which are expressed at a more intermediate level among markers. These two candidates additionally pass more stringent criteria (expressed in ≥ 25% of focal cells) to be considered marker genes (Dataset S1). *DN2695*, the 7th most significant SC marker, is expressed in 47% of SC yet shows no evidence of MC or EDC expression. Interestingly, the two candidate SC-biased genes, *DN2695* and *DN10097*, lie 5.4 kb apart within a 20.1 kb intergenic region on chromosome *2L*. Both EDC-biased candidates, *DN16089* and *DN10930*, are exclusively detected in EDC, although they do not meet our criteria for marker genes. *DN16089* is expressed in 18% of EDC, *DN10930* is expressed in 15% of EDC; neither exhibit SC or MC expression. *DN16089* is located just 79 bp from the EDC marker *sim*, but on the opposite strand. All 11 transcripts are predicted to be non-coding by CPAT. Although getorf identified many putative ORFs (Supplementary Files), BLAST comparisons of predicted proteins to the *D. melanogaster* protein database and the NCBI database of conserved domains returned no significant matches. SignalP revealed no evidence of signal sequences.

## DISCUSSION

Our single-nucleus transcriptome analysis of the primary Drosophila seminal fluid producing organs has validated conjectures in the literature and revealed several new findings. As expected, MC are the primary source of Sfp diversity and exhibit transcriptomes biased toward Sfp production. While several individual Sfps are produced in all three major cell types investigated here, it is notable that the majority of Sfps exhibit strong cell-biased expression, raising the question of why this occurs. Given that these three cell types are spatially separated along the reproductive tract, with the SC distal, the EDC proximal, and the MC intermediate, perhaps there are Sfp “order effects” in assembling the seminal fluid prior to transfer to the female. Order effects have been observed in assembly of the spermatophore in *Pieris rapae* butterflies (Meslin et al. 2017) and seminal fluid in tsetse flies (Odhiambo, Kokwaro, and Sequeira 1983). Such order effects could influence the details of how Sfps bind sperm or interact directly with the female reproductive tract. In spite of the important role for MC in Sfp production, many genes showing MC bias are not annotated as Sfps; their roles in AG function remain to be investigated. SC and EDC transcriptomes are much less biased toward Sfp expression. Indeed, most SC and EDC markers are not Sfps, and most of the genes exhibiting strongly biased expression in these cell types have no known functions in male reproduction. Thus, much of the biology of the AG and ejaculatory duct is still mysterious. Especially notable is the relatively small number of Sfps produced in SC, as first reported by Immarigeon *et al. (2021)*.

Our data confirm that expression of the ‘Sex Peptide network’—Sfps that interact with SP in the female reproductive tract and enhance the PMR (Ravi Ram and Wolfner 2007, 2009; LaFlamme, Ravi Ram, and Wolfner 2012; Singh et al. 2018; Findlay et al. 2014; McGeary and Findlay 2020)—is divided across cell types. *lectin-46Ca*, *lectin-46Cb*, and *CG17575* are SC markers, while *SP*, *aqrs*, *antr*, *intr*, *CG9997*, and *Sems* are MC markers, and *Esp* appears EDC- biased. *frma* and *hdly*, remaining members of the known Sex Peptide network, are not strongly expressed in our dataset. Discovery of the EDC marker *Anion exchanger 2* (*Ae2*), provides a clue about possible functions of the ejaculatory duct apart from Sfp production. In *D. melanogaster*, *Ae2* regulates intracellular pH through Cl^−^/HCO3^−^ exchange in the midgut (Overend et al. 2016) and ovary (Benitez et al. 2019; Ulmschneider et al. 2016). *Ae2* is a highly conserved membrane protein, responsible for pH regulation in the mouse epididymal epithelium, seminiferous tubules, and developing spermatocytes, and is essential for spermatogenesis (Medina et al. 2003). Thus, EDC-biased expression of *Ae2* suggests that the ejaculatory duct may regulate ejaculate pH.

Many of our strongest marker genes are lncRNAs, including markers of our newly defined MC subpopulations. Aside from *iab-8* and *msa* (Maeda et al. 2018), the roles of lncRNAs in AG biology and male reproduction more broadly are uncharacterized, though the possibility that some of these RNAs code for small proteins cannot be ruled out (Immarigeon et al. 2021). Our analysis revealed strong evidence of transcriptionally distinct main cell subclusters. The most obvious distinction between them is that one exhibits evidence of higher transcriptional and translational activity. Many of the markers for these MC subclusters are annotated as lncRNAs, further supporting the possible importance of non-coding RNAs in AG biology. Given that we observe no correlation between *roX1* expression and dosage compensation, *roX1* might have other, uncharacterized functions in the AG. Whether MC subpopulations represent cell subtypes, transitory states, or developmental states, and whether communication among these subclusters occurs, are important questions.

We found evidence for 11 unannotated *D. melanogaster* genes expressed in seminal fluid producing tissues, most of which exhibit strongly cell type-biased expression. Given the low coding potential of these transcripts, and that predicted ORFs exhibit no homology to known proteins and show no evidence of signal sequences required for secretion, their possible functions are mysterious, yet likely relevant to the biology of these three cell types. The two SC-biased genes *DN2695* and *DN10097*, located proximal to one another in a large intergenic region, are particularly interesting candidates for future research on their role in SC biology.

The transcriptomes of the three major cell types investigated here show many similarities between species, as expected given their recent common ancestor. Moreover, interspecific transcriptome divergence among cell types is not occurring independently, supporting the notion that these cell types have correlated functions. Nevertheless, each cell-type exhibits a distinct transcriptome and has distinct evolutionary properties. MC and SC, the two cell types of the AG proper, have less transcriptional divergence from each other than either has from EDC, consistent with more functional and developmental overlap between MC and SC. Overall, interspecific transcriptome divergence is substantially slower for MC than for SC or EDC. However, divergence rates are heterogenous among lineages.. For example, SC transcriptome divergence is substantially greater in the *sim* vs. *yak* comparison than the *mel* vs. *yak* comparison, consistent with the hypothesis accelerated transcriptome evolution along the *sim* lineage for this cell type.

A slightly different picture emerges if one focuses on the most strongly differentially expressed genes between species rather than on overall transcriptome divergence. While the directionality of DE is similar among cell types, the largest expression changes tend to be exhibited in a single cell-type, suggesting that whatever mechanism is driving divergence operates heterogeneously across cell types. EDC generally show the greatest interspecific divergence, though again, the data are consistent with the hypothesis of lineage differences in evolutionary rates. Whether the greater proportion of DE genes among EDC results from directional selection or. relaxed stabilizing selection (Dapper and Wade 2020) is an open question. Many DE genes are Sfps, as expected since Sfps are a major component of these transcriptomes, but notably, the majority of DE genes are not Sfps, raising important questions about the functional axes along which species differences are evolving in these cell types. Indeed, many of the most strongly differentiated genes, which include genes expressed at a high level in some species and apparently unexpressed in others, have unknown functions in these cells in any of the three species. Consistent with transcriptome-wide results, correlations of logFC for DE genes among cell types suggest concerted change, as expected given the closely shared developmental origins of these cell types (Musser and Wagner 2015; Liang et al. 2018) and short time-scales examined in this study. Indeed, correlations of logFC are greatest between MC and SC, which differentiate later in development (Minami et al. 2012; Xue and Noll 2000; Gligorov et al. 2013), compared to EDC cells. Given the limited inquiry into the phenomenon of DE across related cell types in *Drosophila*, however, it is difficult to establish a baseline expectation of concerted change.

Our investigation of the interaction of protein divergence with cell-biased expression and interspecific expression divergence revealed a few salient patterns. As expected given genome wide results (Begun et al. 2007; Langley et al. 2012), directional selection appears to play an important role in driving protein evolution for cell-biased genes. Indeed, *α* values for marker genes, though high, are not obviously different from genome-wide estimates (Fraïsse, Puixeu Sala, and Vicoso 2019), raising interesting questions about whether protein divergence of the AG is unusual in any way. Nevertheless, the relative importance of adaptive divergence appears to vary across cell-types. MC-biased genes are more likely than SC- or EDC-biased genes to show evidence of directional selection. Much of this enrichment results from the strongly Sfp-biased expression of MC, and cell-biased genes that are not Sfps are equally likely to show evidence of protein adaptation for all three cell types. However, conditioning on positive *α*, the relative importance of directional selection is much lower for SC-biased genes than for MC- or EDC-biased genes. Overall, it seems that while adaptive protein evolution is likely common for all cell types, it is most pronounced for MC and least for SC. A speculative hypothesis for this observation is that more beneficial non-synonymous mutations are associated with phenotypes related to establishment of the female PMR, which is primarily a MC function, than with long term maintenance of receptivity to remating, which is in part a SC function (Sitnik et al. 2016). However, it is difficult to make strong statements about the agents of selection driving protein divergence in marker genes without more information on their biological functions in the AG or other tissues and cell types. Finally, we found differentially expressed genes are not more likely than other genes to show evidence of protein adaptation, however, there is a small, significant elevation of positive *α* for DE genes vs. non-DE genes. Thus, while there appear to be some correlations between expression divergence and protein adaptation, the relationship is neither particularly strong nor simple.

While our analyses of single-nucleus transcriptomes in an evolutionary genetics framework has led to many functional and evolutionary findings and hypotheses, perhaps what is most apparent is how little we still understand the biology and evolution of these cells. Many open questions remain about the regulation and function of the seminal fluid producing cells, the biological consequences of species divergence in these cells, and the evolutionary mechanisms shaping this divergence. Continued investigation of closely related species for single-cell phenotypes and population genetic variation will facilitate the fruitful investigation of both functional and evolutionary mechanisms, and help to draw additional connections between these two research domains.

## ACKNOWLEDGEMENTS

This work was supported by the National Institutes of Health grant R35 GM134930 to DJB, a National Science Foundation Graduate Research Fellowship to ACM, and a Pilot and Feasibility Program award from the UC Davis Research Core Facilities Program. We thank the UC Davis Flow Cytometry Shared Resource for performing FACS and the UC Davis Genome Center for single-nucleus library preparation and sequencing. We thank Ben Hopkins and Rachel Thayer for feedback and thoughtful discussions, and the Chiu, Lott, and Kopp labs for sharing materials and equipment.

## DATA AVAILABILITY

Count data for single nuclei in each of the three species, fasta and GTF files for unannotated genes, R scripts, our orthology table, and the list of Sfps used in this study are available in Dataset S10.

## SUPPLEMENTAL METHODS

### Fly stocks and reproductive tract dissection

We used the following sequenced stocks to compare AG transcriptomes between three *melanogaster* subgroup species: *D. melanogaster* RAL 517 (Mackay et al. 2012), *D. simulans w^501^*, and *D. yakuba* Tai18E2 (hereafter referred to as *mel*, *sim*, and *yak*). All animals were raised on a cornmeal-molasses-agar medium at 25°C and 60% relative humidity, on a 12:12 light/dark cycle. For snRNA-Seq experiments, virgin male flies were collected and placed into fresh vials of food in groups of five males per vial. On the day of the experiment, 2-3 day old virgin males were anaesthetized with CO2 and their accessory glands plus anterior ejaculatory duct were dissected in cold 1X PBS and moved to 1X PBS + 2% BSA (Sigma SRE0036) on ice. Animals were dissected between Zeitgeber time (ZT) 0:30 and 2:30. Five animals from each species (derived from a single vial of food) were dissected before moving to the next species, in a repeating pattern, to prevent biasing interspecific sampling due to potential effects of circadian rhythms or the dissection protocol. Together, 23 *mel* (13 three day old, 10 two day old), 26 *sim* (15 three day old, 11 two day old), and 24 *yak* (14 three day old, 10 two day old) were dissected, and tissue from all three species was pooled together into a single microfuge tube.

### Nuclear isolation and purification

Our nuclear isolation protocol is based on Luciano Martelotto’s (2019) with some modification. For all steps we used non-stick, nuclease-free polypropylene tubes and pipette tips. All plastics, glassware, and filters were rinsed with PBS + 2% BSA before use. Pooled reproductive tract tissue was washed once with PBS + 2% BSA, then resuspended in 1 mL of lysis buffer (10 mM Tris-HCl pH 7.4, 10mM NaCl, 3 mM MgCl2) with 0.1% NP40 (Sigma 9002-93-1). The tissue and lysis buffer were moved to a glass dounce homogenizer on ice and incubated for five minutes, with intermittent low-speed vortexing. The tissue was then dounce homogenized with 25 strokes with pestle A (loose fit) and incubated for an additional 10 minutes with intermittent low-speed vortexing. The tissue was next dounce homogenized with 25 strokes with pestle B (tight fit). 0.5 mL of PBS + 2% BSA was added to the homogenate, and the mixture was triturated 15 times with a silanized fire-polished glass pasteur pipette (BrainBits FPP). The homogenate was filtered through a 35 micron mesh (Falcon 352235) to remove unlysed tissue, and an additional 100 uL of PBS + 2% BSA was rinsed through the filter afterwards. The filtrate was centrifuged at 500 rcf for eight minutes at 4°C. The supernatant was removed, 200 uL of PBS + 2% BSA was added to the pellet, and the pellet was incubated on ice for five minutes. The pellet was then resuspended before an additional 800 uL of PBS + 2% BSA was mixed into the solution. Nuclei were centrifuged again at 500 rcf for eight minutes at 4°C, the supernatant removed, and the pellet was incubated in 200 uL of PBS + 2% BSA with 10 ug/mL DAPI on ice for five minutes before being resuspended.

### FACS and snRNA-Seq

Following isolation of semi-pure nuclei, we used Fluorescence Activated Cell Sorting (FACS) to further purify singlet nuclei from clumps and cell debris. FACS was gated using DAPI and singlet nuclei were sorted into a new tube. Library preparation and sequencing was performed by the UC Davis Genome Center DNA Technologies & Expression Analysis Core Laboratory. The 10X Genomics 3’ Single Cell (v2) kit was used to create libraries for snRNA-Seq on purified nuclei, and libraries were then sequenced on one lane of an Illumina HiSeq4000.

### Bioinformatic assignment of species origin, RNA-Seq alignment, QC, and orthologue formatting

We used custom perl scripts to identify reads originating from droplets containing nuclei vs. those that were empty and therefore composed entirely of ambient background RNA. For 10X Chromium v3, the R1 fastq contains droplet barcode and UMI data, while the R2 fastq contains cDNA sequence (Zheng et al. 2017). We used the R1 fastq file to parse counts of unique molecular identifiers (UMIs) per droplet barcode. After gathering this count data, we examined the distribution of UMIs per barcode in descending rank-order, to identify the ‘knee’ inflection point separating true cell-containing barcodes from barcodes associated with empty droplets (Macosko et al. 2015). For initial alignment of reads we selected all barcodes above the inflection point, which we expected was an overestimate of the true number of singlet nuclei, prior to further downstream filtering of low-UMI nuclei and multiplet-containing droplets. Given that single-nucleus RNA-Seq typically has a higher background RNA content than single-cell RNA-Seq (Alvarez et al. 2020), we wanted to profile this RNA background for later downstream bioinformatic correction. To profile background RNA, we selected 1,000 empty droplets at random from rank-orders below the inflection point.

We used an alignment-based approach to determine species-of-origin for each cell. For each cell we aligned all reads to each species’ genome (*D. melanogaster* version 6.19, *D. simulans versio*n 2.02, *D. yakuba* version 1.05) respectively, using Hisat2 v2.1.0 (Kim et al. 2019). We sampled five million aligned reads and retained those with a best alignment match to a single species, dropping reads that aligned best equally well to two or more species. We used this subset of aligned reads to determine the percentage of reads originating from each species for each barcode. We selected those barcodes where ≥ 50% of the reads aligned best to a single genome and categorized them as barcodes for that species. Next, we selected the R1 and R2 reads corresponding to barcodes associated with putative nuclei, and a new set of fastq files was created for each barcode, removing duplicated UMIs from these files. For empty droplets, we selected the R1 and R2 reads corresponding to each of the randomly sampled barcodes and created new fastq file pairs for each empty droplet. We aligned raw R2 reads to the appropriate species’ genome (*D. melanogaster* version 6.33, *D. simulans versio*n 2.02, *D. yakuba* version 1.05) using STAR v2.7.5a (Dobin et al. 2013) with default parameters. We removed all transcripts of the gene *mod(mdg4)* from the *D. melanogaster* GTF file used for alignment, as this gene has trans-spliced transcripts without meaningful strand data, which causes a STAR error. We parsed the STAR logfile of each cell for percent of reads unmapped, percent of reads multi-mapped, and number of total mapped reads. We then used an R script to visualize these data and choose cut- offs to filter probable multiplets, as well as filtering low-quality/high background nuclei from the data.

In our first pass analysis of the data we discovered an apparent lack of *Abd-B* expression in secondary cells of *sim* and *yak*, which was highly unexpected due to the central role of *Abd-B* in *mel* secondary cell development. Investigating possible technical explanations, we found discrepancies in the completeness of exon annotation across the three species, with the annotated *Abd-B* orthologues in *sim* and *yak* significantly shorter than that of *mel*. Since the Chromium library prep produces reads beginning around 300-400 bases from 3’ end of expressed transcripts, rather than across the length of the transcript as in typical bulk mRNA-Seq library preps, differences in exon annotation of orthologous genes, especially at the 3’ end, may lead to artifactual inference of DE across species. To investigate this possibility we used de novo transcriptome assemblies from paired-end bulk tissue mRNA-Seq samples to improve the annotation of all AG-expressed transcripts, prior to counting features in our aligned single-nucleus data. Transcripts for *D. simulans* (*w^501^*) and *D. yakuba* (Tai18E2) were generated using Trinity v2.11.0 (Grabherr et al. 2011). We then aligned these transcripts back to the appropriate species genome and transcriptome to identify assembled transcripts that aligned to existing genes. Records for these new transcripts were added to the species’ GTF files from FlyBase. Custom GTF files with updated *sim* and *yak* exon annotation can be found in our Supplemental Data. Following this improved annotation, count data exhibited similar *Abd-B* expression in secondary cells across species, as expected.

Next, we counted features from BAM files using HTSeq-count v0.12.3 (Anders, Pyl, and Huber 2015) with default parameters. Given that intronic reads may be included in single-nucleus RNA-Seq (Lake et al. 2016), we counted reads mapping to either exons or introns. Empty droplets were treated similarly to nuclei, and were independently processed using each of the three species’ annotations. Although the “true” background RNA profile is not expected to vary by the species of the nucleus contained in each droplet, the alignment and feature counting steps will filter out some number of reads of discordant species origin that are sufficiently divergent, so background profiles at this step will depend on the species-specific genome and gene models used.

To remove background RNA from nuclei, we used the R package SoupX v1.4.8 (Young and Behjati 2020). We performed preliminary cell type clustering of uncorrected data to identify marker genes in Seurat v3.2.2 (Stuart et al. 2019), the basis for the background correction algorithm used by SoupX. We also did additional filtering of high UMI-count cells to remove outliers (potential doublets). This final filtering left us with 1167 *mel*, 2115 *sim*, and 989 *yak* nuclei. We performed preliminary Seurat analysis independently for each species using its full set of genes (not limited to 1-to-1-to-1 orthologues). The set of empty drops aligned to each respective species was used as the basis for estimation of the background transcriptome, and background correction was independently performed for each species.

For comparative analyses we created a set of 1-to-1-to-1 orthologues (11,481 genes) using the *D. melanogaster* orthologue table from Flybase (2020 version 2). We used an R script to identify the set of *mel* orthologues with single orthologues identified in *sim* and *yak* respectively. To maximize the number of Sfps included in our comparative analysis we compiled the unique set of Sfps published from two proteomic studies, Findlay *et al*. (2009), and Sepil *et al*. (2018). Of these 264 Sfps, 77 were not in our set of annotated 1-to-1-to-1 orthologues. Of these, we were able to manually curate 25 novel 1-to-1-to-1 orthologous genes (Table S5) using tblastx (Camacho et al. 2009) and Ensembl Metazoa genome browsers (Howe et al. 2020) to confirm synteny.

### Marker gene identification and differential expression among species

All analyses of single-nucleus gene expression data were performed in R v3.6.1 using Seurat v3.2.2 (Satija et al. 2015; Butler et al. 2018; Stuart et al. 2019). We used two parallel approaches. We did an integrated analysis of the data across species, using our set of *mel*, *sim*, and *yak* 1-to- 1-to-1 orthologues. We also performed an independent analysis of *mel* using all annotated genes to get a fuller picture of differences in gene expression among cell types. For both datasets we used a fairly standard Seurat workflow. We log-normalized UMI counts and z-score transformed counts on a gene-by-gene basis. We selected the top 2000 variable genes for downstream analysis by ranking their dispersion values. We chose not to use the usual variance-stabilizing transformation (VST) method, which normalizes dispersion by absolute expression to account for noise inherent in single-cell RNA-Seq (Hafemeister and Satija 2019). Our dataset has one dominant cell type (MC), and two rare cell types (SC and EDC). The nature of the data means that the most highly expressed, specific markers to MC tend not to show a very high variability over the entire dataset. As expected, VST did not include several important MC marker genes including *SP* and *Acp95EF*, while the dispersion method includes them. We clustered cells with the Shared Nearest Neighbor (SNN) algorithm and used UMAP dimensionality reduction to visualize clustering. We identified marker genes using Seurat’s FindAllMarkers() method and assessed significance using a Wilcoxon Rank Sum test. We required marker genes to be expressed in at least 25% of focal cluster cells, and set a minimal average logFC requirement of We filtered marker genes to those with Bonferroni-corrected p-values less than 0.05. To further investigate cell type specific expression bias of all SFPs, in addition to those strictly classified as marker genes, we did not impose minimum percent cells expressing and average logFC thresholds. We additionally identified markers distinguishing MC subpopulations from one another using the FindMarkers() method. Further details of the specific parameters and methods used to process data in Seurat can be found in our R scripts.

We used limma v3.42.2 (Ritchie et al. 2015) to infer DE genes for each cell type. We performed pairwise contrasts among the three species, and classified genes as DE with an FDR of 5% (Benjamini and Hochberg 1995). Further details of the limma analysis can be found in our R scripts. To compare the rate of qualitative expression divergence across cell types, we calculated ratios of DE genes at various log2 fold change (logFC) cut-offs across the three cell types, for each of the three species contrasts, and tested for differences in these ratios using a G-Test of goodness-of-fit (Sokal and Rohlf 2012). To test for differences in the magnitude of expression differences across cell types, we similarly compared distributions of absolute values of logFC using a Kruskal-Wallis test (Kruskal and Wallis 1952). Finally, we examined overall expression correlations between species, within cell types, by calculating average expression per gene and Pearson correlation coefficients.

To examine the relative level of concerted vs independent gene expression evolution across cell types, we subset the data to the set of differentially expressed genes exhibiting a logFC greater than one in at least one cell type specific pairwise species contrast (out of a total of nine contrasts: three cell types X three species). We then calculated pairwise Pearson correlation coefficients of logFC across cell types within each of the three pairwise species contrasts. We permuted logFC values across genes 10,000 times to obtain a distribution of Pearson correlation coefficients under the null expectation of entirely cell type independent change within our set of DE genes.

### Population genetic inference of adaptive protein divergence of marker genes

To investigate potential differences in the prevalence of adaptive protein evolution across cell types, we used existing population data (2019) from *D. melanogaster* (Lack et al. 2015) with *D. simulans* as the outgroup. The McDonald-Kreitman test (McDonald and Kreitman 1991) compares the ratios of polymorphic and fixed synonymous amino acid substitutions to nonsynonymous substitutions. The summary statistic *α* (Smith and Eyre-Walker 2002) represents an estimate of excess fixed amino acid substitutions relative to the expectation under strict neutrality, describing the predicted proportion of amino acid substitutions resulting from directional selection. A positive value of *α* suggests directional selection acting on a given gene. Among positive *α* values, a greater value for a given set of genes suggests a greater proportion of amino acid substitutions fixed under directional selection. We considered two summaries of the role of adaptation in protein divergence: the proportion of marker genes with *α* > 0, and the distribution of *α* values amongst those genes with *α* > 0. The proportions of positive *α* values were compared using Fisher’s exact test, with post-hoc pairwise tests between cell types. The distributions of positive *α* values were visualized in ggplot2 v3.3.3 (Wickham 2016), and compared using a Kruskal-Wallis test with post-hoc pairwise Wilcoxon tests.

### De novo transcriptome assembly and identification of unannotated *D. melanogaster* transcripts

We first identified the set of cell barcodes corresponding to each of the three cell types (MC, SC, EDC) based on marker gene identification as described earlier. Next, we concatenated raw fastq files corresponding to each barcode together to create a set of reads originating from each population of cells. We used TrimGalore! v2.11.0 (https://github.com/FelixKrueger/TrimGalore) to prep the raw reads, followed by de novo transcriptome assembly using Trinity v2.11.0 (Grabherr et al. 2011). Since transcript content is only contained in the R2 read of 10X Chromium libraries, our data is effectively single-end, so we used the ‘--single’ and ‘--SS_lib_type F’ options to Trinity. We used a BLAST-based strategy (Camacho et al. 2009) to identify candidate unannotated transcripts in *D. melanogaster*. First, using blastn we aligned all transcripts to two custom databases, A) the sequences of all annotated genes in *mel*, *sim*, and *yak*, and B) the whole-genome sequences of all three species. We aligned the *mel* de novo transcriptome to all three species’ concatenated databases to avoid having background transcripts from *sim* or *yak* identified as unannotated in *mel*. We then took the set of transcripts that had at least one BLAST hit to the genomic database, but no BLAST hits to the database of annotated genes.

In the course of this analysis we discovered template-switching oligo (TSO) concatemer occurrence at the 5’ end of some transcripts. TSO concatemers in 10X Chromium libraries have been previously documented (Svensson 2017). We removed the sequence (AAGCAGTGGTATCAACGCAGAGTACATGGG) from the 5’ end of assembled transcripts. We also checked the raw data for TSO occurrence and found that these sequences exclusively occur at the 5’ end, precluding the possibility of artifactual chimeric transcripts formed during Trinity assembly. Additionally, these concatemers are not expected to affect alignment of reads to the genome since STAR automatically trims non-aligning adapter sequences (Dobin et al. 2013).

We used counts from Salmon v0.12.0 (Patro et al. 2017) to filter the set of assembled transcripts to those expressed at a level of counts greater than 10% of the total number of cells in at least one cell type, to filter out very-lowly expressed transcripts. We removed poly-A sequence from the transcripts, and we aligned these filtered transcripts back to a blastn database containing solely the *mel* genome to remove background transcripts from other species. We converted tabular BLAST output to a GTF file using a python script, and used HTSeq to do feature counting on individual cell transcriptomes aligned with STAR (described earlier), as well as background drop transcriptomes. We used SoupX to remove background contamination as described above, with the global contamination fraction estimated from earlier all-genes analysis. Given the single-ended, 3’-biased nature of the 10X library prep, full-length transcript assemblies cannot be confidently constructed. Therefore, we augmented this analysis by making a de novo transcriptome assembly for the FlyAtlas2 (Leader et al. 2018) accessory gland data using Trinity (Grabherr et al. 2011). We identified homologous FlyAtlas2 transcripts using BLAST. These transcripts were used to improve the completeness of our candidate transcripts and eliminate a few candidates that overlapped annotated genes. Similarly, for SC-biased transcripts we used bulk RNA-Seq data from FACS-sorted secondary cells (Immarigeon et al. 2021) to improve our annotations. Both supplementary RNA-Seq datasets are unstranded, so we obtained strand information from our 10X-based transcript assemblies. We used FlyAtlas2 data to improve the annotation of *DN8354, DN35169, DN10930, DN16089, DN2736, DN5813,* and *DN818*. We found no matching FlyAtlas2 transcripts for *DN4707*, *DN11110*, and *DN10097*. We found a single, very lowly expressed FlyAtlas2 transcript homologous to *DN2695*, but the 10X transcript was longer than the FlyAtlas2 transcript. We searched for transcripts homologous to *DN2695* and *DN10097* in FACS-sorted secondary cell RNA-Seq data (Immarigeon et al. 2021). We found matching transcripts in this dataset expressed at significant levels: *DN2695* at a mean 20.8 TPM and *DN100097* at a mean 1.12 TPM. This improved our assembled *DN2965* transcript length from 463 to 2,176 bp, and took *DN10097* from 260 to 353 bp.

We used Ensembl Metazoa BLAST and the *mel* genome browser (Howe et al. 2020) to identify gene coordinates, strand, and neighboring annotated genes, and to verify that these candidate genes do not overlap with any existing annotated features. Though we selected our candidate gene sequences based on expression, Trinity groups sets of transcripts together based on shared k-mers—many of which are very lowly expressed. For each of the 11 candidates, we additionally verified that none of the clustered transcripts within the gene “family” overlapped with annotated features.

For cell type-specific analysis of unannotated gene expression, we added counts to the broader *mel* dataset post-hoc. We used Seurat’s FindAllMarkers() method to identify cell type bias of unannotated genes. To adjust the test for identification of cell type specific bias, rather than identification of canonical cell type markers, we relaxed the default thresholding requirements, including the requirement that a gene be expressed in at least 25% of focal cluster cells, and the minimal logFC requirement of 0.25. Significance of expression bias was assessed using a Wilcoxon Rank Sum test with Bonferroni multiple test correction. To identify potential ORFs, we used EMBOSS getorf (Rice et al. 2000). We attempted to characterize these potential ORFs further using Ensembl Metazoa Protein BLAST (Howe et al. 2020) to the database of all *mel* proteins, NCBI’s Conserved Domain Database search tool (Lu et al. 2020), and SignalP v5.0 (Almagro Armenteros et al. 2019) to identify putative signal sequences. We additionally assessed coding potential of our transcripts using CPAT v2.0.0 (L. Wang et al. 2013).

## SUPPLEMENTARY FIGURES AND TABLES

**Figure S1.**
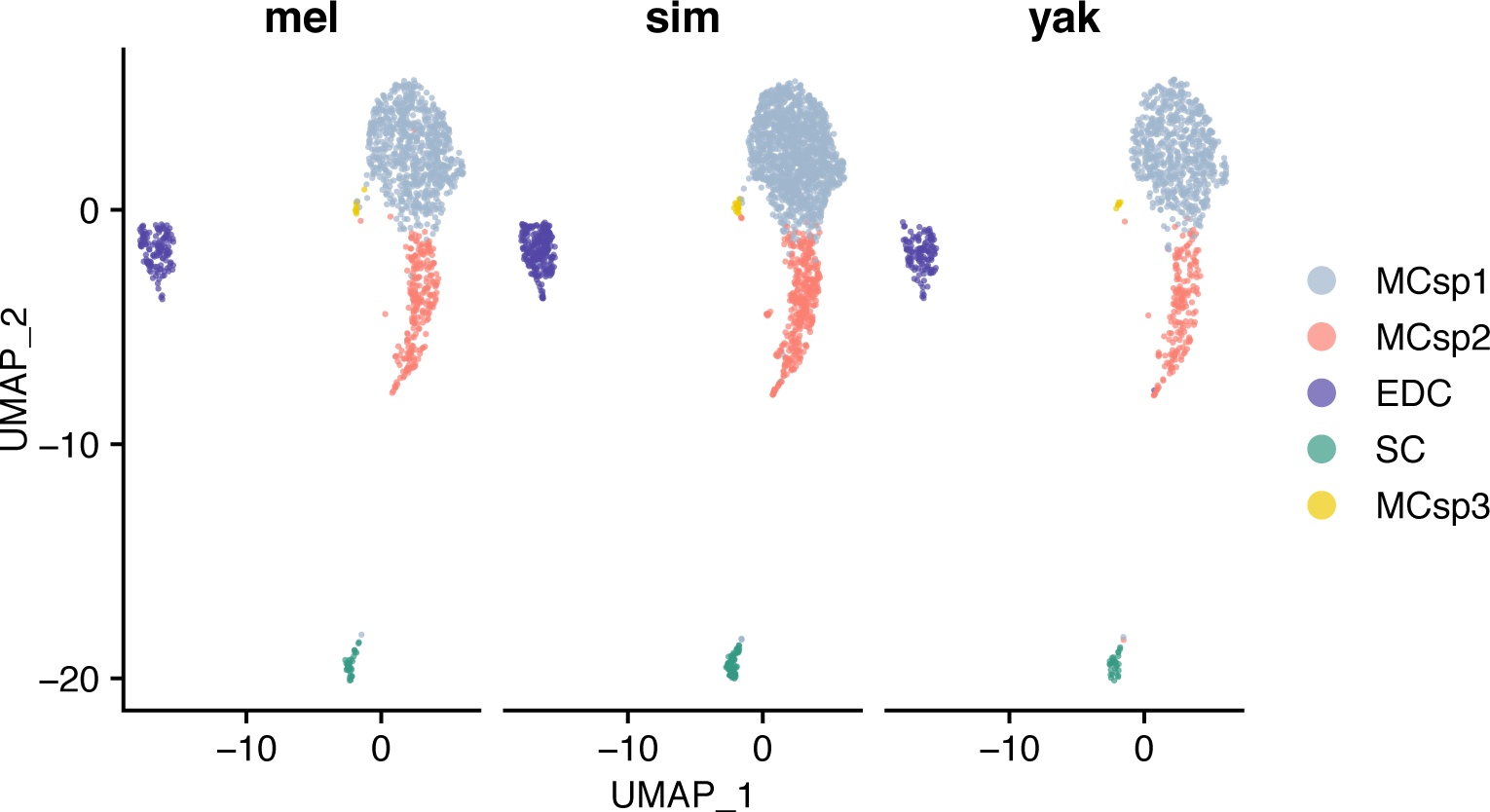
Subpopulations of MC appear conserved among *mel*, *sim*, and *yak*. MCsp2, explored in depth in *melanogaster*, is conserved with roughly equal proportionality among species. The data suggest an additional subpopulation of MC.

**Figure S2.**
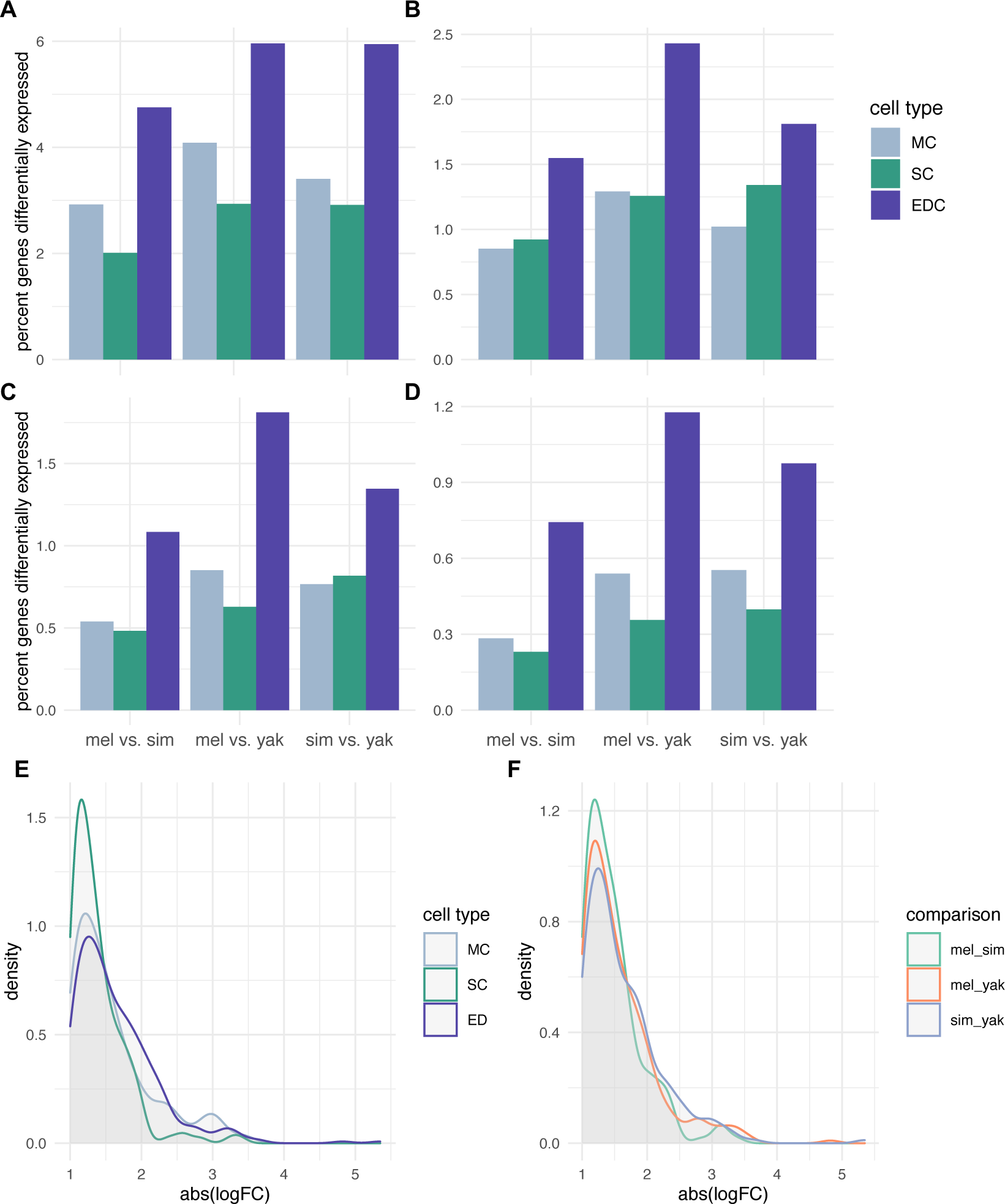
Patterns of DE among cell types are robust to a range of logFC cutoffs. (*A*): logFC *≥* 0.5, (*B*): *≥* 1, (*C*): *≥* 1.25, (*D*): *≥* 1.5. While there is a trend towards MC enrichment at the lowest and highest cutoffs, MC and SC are not significantly different in any intraspecific comparison (Wilcoxon rank sum tests, *p* > 0.05). The magnitude of DE does not vary among (*E*) cell types or (*F*) species contrasts (Kruskal-Wallis tests, *p* > 0.05).

**Figure S3.**
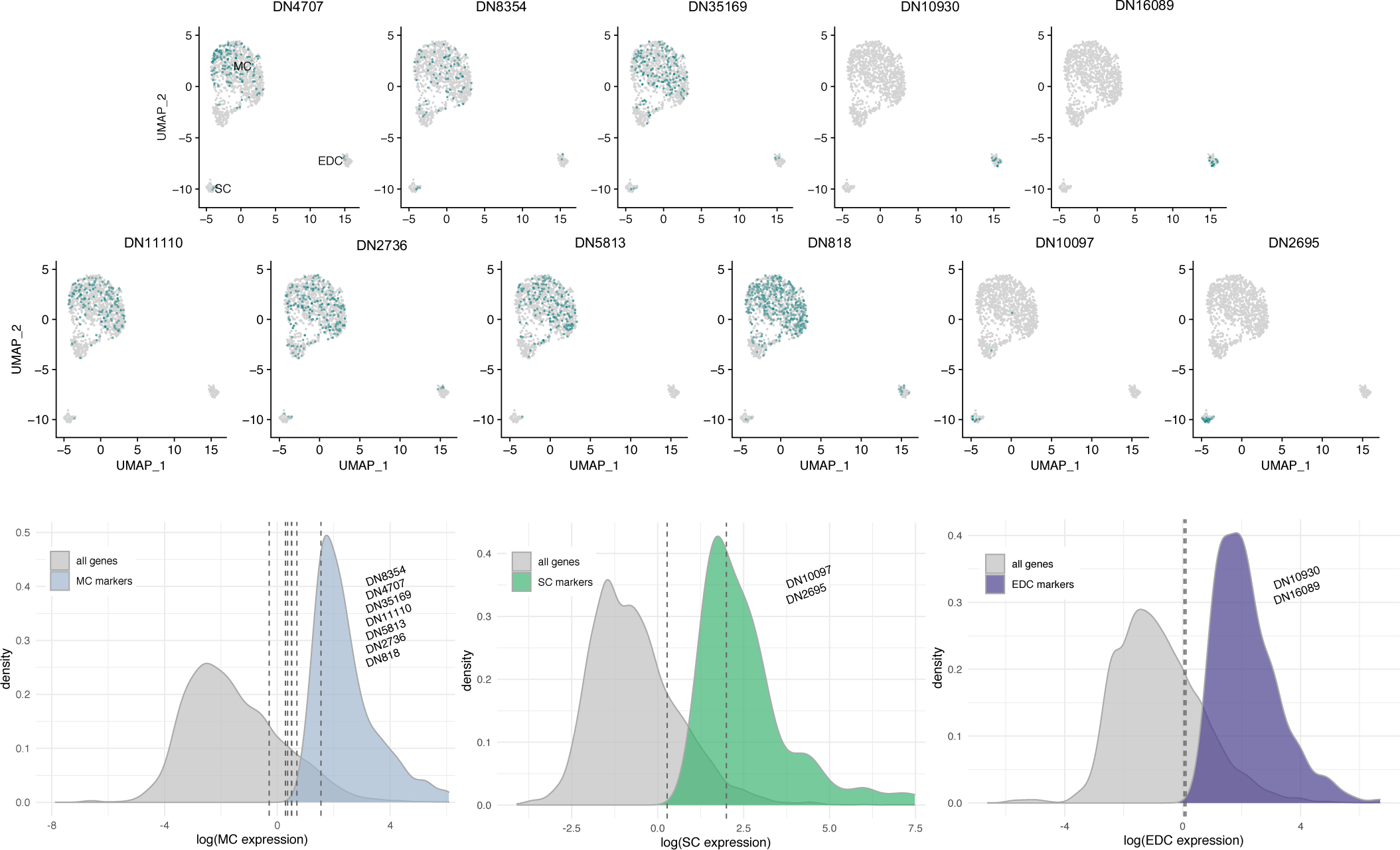
(*A*) UMAP showing three cell types in *mel*, along with expression of unannotated candidate genes (Table 1) indicated in teal. (*B*) Distributions of gene expression among all expressed genes (grey) and marker genes (color), for MC, SC, and EDC. Expression values for unannotated genes are overlayed. These data indicate that most unannotated genes are expressed at a relatively high level among all expressed genes, but a very low level among marker genes. Two exceptions to this trend are *DN818* in MC and *DN2695* in SC—the only two unannotated genes that are also characterized as marker genes.

**Table S1.** Sfp marker genes in *mel* SC and EDC, as described in Dataset S1. pct.1 refers to the fraction of focal cells expressing the marker, pct.2 refers to the fraction on non-focal cells expressing the marker. *p* is the result of a Wilcoxon rank-sum test with Bonferroni correction.

| gene | cluster | avg logFC | pct.1 | pct.2 | *p* |
| --- | --- | --- | --- | --- | --- |
| *CG14913* | SC | 3.109 | 0.780 | 0.001 | 5.37E-189 |
| *CG17575* | SC | 4.598 | 1.000 | 0.021 | 3.71E-172 |
| *CG3349* | SC | 1.711 | 0.320 | 0.001 | 6.09E-72 |
| *CG13965* | SC | 3.692 | 1.000 | 0.139 | 6.00E-68 |
| *lectin-46Ca* | SC | 4.504 | 1.000 | 0.169 | 2.11E-60 |
| *CG9029* | SC | 4.328 | 1.000 | 0.186 | 1.60E-56 |
| *Acp32CD* | SC | 4.606 | 1.000 | 0.207 | 3.82E-53 |
| *lectin-46Cb* | SC | 4.527 | 1.000 | 0.237 | 6.51E-49 |
| *mfas* | SC | 1.186 | 1.000 | 0.813 | 5.12E-16 |
| *Pgant9* | SC | 0.619 | 0.720 | 0.383 | 0.00102 |
| *Dup99B* | EDC | 6.457 | 1.000 | 0.002 | 1.32E-243 |
| *Obp51a* | EDC | 4.210 | 1.000 | 0.003 | 2.87E-239 |
| *Anp* | EDC | 4.879 | 1.000 | 0.003 | 3.40E-239 |
| *Spn77Bc* | EDC | 5.066 | 1.000 | 0.006 | 8.92E-226 |
| *CG18258* | EDC | 2.809 | 0.797 | 0.001 | 8.95E-194 |
| *CG17549* | EDC | 3.056 | 0.831 | 0.013 | 4.41E-161 |
| *CG42782* | EDC | 3.457 | 0.966 | 0.030 | 1.67E-150 |
| *CG5162* | EDC | 3.741 | 0.983 | 0.042 | 5.18E-142 |
| *CG17242* | EDC | 3.371 | 0.966 | 0.042 | 1.45E-136 |
| *CG5402* | EDC | 3.561 | 0.966 | 0.049 | 5.77E-129 |
| *CG15394* | EDC | 2.950 | 0.898 | 0.040 | 1.79E-124 |
| *Spn77Bb* | EDC | 5.154 | 1.000 | 0.084 | 8.22E-105 |
| *Est-6* | EDC | 3.979 | 0.983 | 0.082 | 1.59E-101 |
| *Treh* | EDC | 2.358 | 0.847 | 0.055 | 2.51E-93 |
| *Gld* | EDC | 2.441 | 0.763 | 0.046 | 4.62E-86 |
| *betaggt-I* | EDC | 2.132 | 0.559 | 0.017 | 1.14E-83 |
| *CG34034* | EDC | 4.023 | 1.000 | 0.175 | 3.77E-68 |
| *Sfp93F* | EDC | 3.023 | 0.966 | 0.161 | 8.77E-65 |
| *NT5E-2* | EDC | 1.118 | 0.254 | 0.009 | 7.80E-33 |
| *CG31704* | EDC | 1.937 | 0.983 | 0.450 | 1.13E-32 |
| *trx* | EDC | 0.491 | 0.356 | 0.106 | 0.00106 |

**Table S2.** The top 12 DE genes in each cell type. logFC refers to the magnitude of expression difference among species contrasts. F is the moderated F-statistic from limma. *p* is Benjamini-Hochberg corrected. Average expression values are provided for each species as log-transformed counts.

| gene | cluster | logFC(mel/sim) | logFC(mel/yak) | logFC(sim/yak) | F | *p* | mel | sim | yak |
| --- | --- | --- | --- | --- | --- | --- | --- | --- | --- |
| Acp95EF | MC | 0.762 | 1.751 | 0.989 | 1243 | 0.00E+00 | 6.48 | 5.75 | 4.92 |
| Pkc53E | MC | -0.600 | -3.114 | -2.514 | 1392 | 0.00E+00 | 1.96 | 2.45 | 4.38 |
| Cys | MC | -0.054 | -2.514 | -2.460 | 2154 | 0.00E+00 | 0.61 | 0.78 | 3.46 |
| sv | MC | -0.053 | -1.929 | -1.876 | 1242 | 0.00E+00 | 0.34 | 0.55 | 3.12 |
| Fkbp14 | MC | -2.159 | 0.430 | 2.589 | 1592 | 0.00E+00 | 1.69 | 3.54 | 1.04 |
| Acp63F | MC | 0.592 | 2.292 | 1.699 | 1020 | 0.00E+00 | 4.72 | 4.19 | 3.10 |
| Acp76A | MC | 0.585 | 3.490 | 2.905 | 1971 | 0.00E+00 | 4.13 | 3.64 | 0.23 |
| Acp62F | MC | 0.172 | 2.092 | 1.920 | 998 | 0.00E+00 | 4.93 | 4.82 | 3.40 |
| CG7329 | MC | -1.709 | 0.560 | 2.269 | 1055 | 0.00E+00 | 1.70 | 3.30 | 1.00 |
| Obp58b | MC | -3.148 | -0.837 | 2.311 | 2511 | 0.00E+00 | 0.60 | 3.84 | 1.93 |
| Marf1 | MC | 1.888 | 1.891 | 0.003 | 1568 | 0.00E+00 | 3.00 | 0.01 | 0.00 |
| Obp56f | MC | -1.693 | 0.650 | 2.343 | 970 | 0.00E+00 | 2.78 | 4.04 | 2.32 |
| Nhe3 | SC | -3.205 | -0.127 | 3.078 | 107.8 | 3.00E-25 | 1.28 | 4.17 | 1.44 |
| Pex19 | SC | 1.910 | 1.985 | 0.075 | 72.2 | 8.37E-19 | 2.75 | 0.44 | 0.13 |
| SPR | SC | 0.000 | -1.635 | -1.635 | 50.7 | 4.98E-14 | 0.00 | 0.00 | 2.57 |
| nahoda | SC | 1.044 | -1.600 | -2.645 | 48.4 | 1.42E-13 | 2.57 | 1.49 | 3.47 |
| MFS17 | SC | 2.371 | 1.090 | -1.281 | 47.0 | 2.61E-13 | 2.95 | 0.00 | 2.18 |
| msi | SC | 2.773 | 1.419 | -1.354 | 45.3 | 6.00E-13 | 4.00 | 1.87 | 3.00 |
| sba | SC | 0.791 | -1.812 | -2.603 | 43.2 | 1.87E-12 | 2.92 | 2.16 | 4.07 |
| CG31145 | SC | 1.789 | -0.132 | -1.921 | 42.6 | 2.28E-12 | 3.76 | 2.60 | 4.03 |
| ogre | SC | -0.099 | -1.828 | -1.729 | 39.7 | 1.26E-11 | 1.07 | 1.15 | 2.83 |
| obst-B | SC | -1.545 | -0.147 | 1.398 | 37.4 | 4.89E-11 | 0.04 | 2.51 | 0.71 |
| Slip1 | SC | -1.610 | -0.056 | 1.554 | 34.8 | 2.47E-10 | 0.72 | 2.60 | 0.84 |
| Cln7 | SC | 1.261 | 1.533 | 0.272 | 32.0 | 1.49E-09 | 2.45 | 0.94 | 0.00 |
| Est-6 | EDC | -0.563 | 4.782 | 5.345 | 2782 | 0.000 | 5.10 | 5.50 | 0.00 |
| CG6040 | EDC | -0.065 | 3.574 | 3.639 | 680 | 0.000 | 4.43 | 4.43 | 1.61 |
| mbl | EDC | -1.522 | 0.714 | 2.237 | 535 | 0.000 | 4.48 | 5.97 | 3.99 |
| Sfp24C1 | EDC | -3.121 | 0.125 | 3.246 | 513 | 0.000 | 0.46 | 3.69 | 0.00 |
| CG10031 | EDC | -3.161 | 0.000 | 3.161 | 508 | 0.000 | 0.00 | 3.57 | 0.00 |
| hdly | EDC | 0.036 | -2.351 | -2.387 | 443 | 0.000 | 0.24 | 0.04 | 3.21 |
| Gld | EDC | -1.575 | 1.775 | 3.350 | 407 | 0.000 | 3.13 | 4.17 | 1.64 |
| GlcAT-P | EDC | 0.096 | -2.196 | -2.292 | 394 | 0.000 | 0.27 | 0.00 | 3.16 |
| MFS17 | EDC | 2.357 | 0.378 | -1.979 | 389 | 0.000 | 2.94 | 0.00 | 2.84 |
| SK | EDC | 0.631 | -1.816 | -2.447 | 337 | 0.000 | 1.62 | 0.16 | 3.18 |
| CG43060 | EDC | 1.950 | 1.950 | 0.000 | 301 | 0.000 | 2.99 | 0.00 | 0.00 |
| Mi-2 | EDC | -0.565 | -3.330 | -2.765 | 288 | 0.000 | 0.00 | 1.29 | 3.71 |

**Table S3.** p values (Benjamini-Hochberg corrected) of pairwise *G*-tests for percentage of expressed genes DE among cell types and species contrasts. ms = *mel* vs *sim,* my = *mel* vs *yak,* sy = *sim* vs *yak*; MC = main cells, SC = secondary cells, EDC = ejaculatory duct cells. Significant values in bold face.

|  | *ms_MC* | *ms_SC* | *ms_ED* | *my_MC* | *my_SC* | *my_ED* | *sy_MC* | *sy_SC* |
| --- | --- | --- | --- | --- | --- | --- | --- | --- |
| *ms_SC* | 0.750 | - | - | - | - | - | - | - |
| *ms_ED* | **0.001** | **0.009** | - | - | - | - | - | - |
| *my_MC* | **0.027** | 0.099 | 0.276 | - | - | - | - | - |
| *my_SC* | 0.061 | 0.171 | 0.272 | 0.872 | - | - | - | - |
| *my_ED* | **0.000** | **0.000** | **0.001** | **0.000** | **0.000** | - | - | - |
| *sy_MC* | 0.352 | 0.662 | **0.018** | 0.193 | 0.301 | **0.000** | - | - |
| *sy_SC* | **0.027** | 0.090 | 0.423 | 0.839 | 0.760 | **0.000** | 0.171 | - |
| *sy_ED* | **0.000** | **0.000** | 0.303 | **0.031** | **0.037** | **0.031** | **0.000** | 0.087 |

**Table S4.** Unannotated candidate genes expressed in the *D. melanogaster* accessory gland. Coordinates are from BLAST results to *D. melanogaster* version 6.36. Length refers to the span of BLAST coordinates in base pairs. MC, SC, and EDC refer to the fraction of genes in each cell type with expression, respectively. Average logFC is the cell type with highest fraction of expression compared to the other two cell types. *p* is the result of a Wilcoxon Rank Sum test with Bonferroni correction.

| transcript | coordinates | strand | length | neighbor genes (dist., strand) | expression bias | MC | SC | EDC | logFC | *p* |
| --- | --- | --- | --- | --- | --- | --- | --- | --- | --- | --- |
| DN4707 | 3R:29346322-29346674 | F | 352 | *lncRNA:CR45547* (9.4 kb, R); *Cnx99A* (26.7 kb, R) | broad | 0.121 | 0.060 | 0.050 | 0.544 | 0.309 |
| DN8354 | 2R:20149969-20150499 | R | 530 | *lncRNA:CR45229* (0.2 kb, F); *CG1041* (3.2 kb, F) | broad | 0.074 | 0.060 | 0.017 | 0.255 | 1 |
| DN35169 | 3R:26745224-26745903 | R | 630 | *CG14247* (6.6 kb, F); *lncRNA:CR45562* (11.4 kb, R) | broad / MC | 0.141 | 0.060 | 0.050 | 0.595 | 0.087 |
| DN10930 | 3R:13056814-13057676 | R | 863 | *pic* (0.1 kb, F); *sim* (0.1 kb, F) | EDC | 0.000 | 0.000 | 0.150 | 0.750 | <0.001 |
| DN16089 | 3R:14099143-14099715 | R | 572 | *PK1-R* (0.1 kb, R); *CG9920* (1.1 kb, R) | EDC | 0.000 | 0.000 | 0.183 | 0.718 | <0.001 |
| DN11110 | X:9316379-9316731 | F | 352 | *lncRNA:CR44534* (1.2kb, F); *c1.21* (1.2kb, R) | MC | 0.149 | 0.020 | 0.017 | 0.923 | 0.001 |
| DN2736 | 2L:9747046-9747785 | R | 739 | *Cyp4e3* (0.05 kb, R); *Nckx30C* (0.5 kb, R) | MC | 0.177 | 0.060 | 0.050 | 0.856 | 0.006 |
| DN5813 | 2R:16922917-16924195 | F | 1278 | *Pkc53E* (0.5 kb, F); *inaC* (21 kb, F) | MC | 0.160 | 0.000 | 0.017 | 0.981 | <0.001 |
| DN818 | 3R:7341782-7344884 | R | 3102 | *lncRNA:CR44337* (9.7 kb, F); *CG31496* (0.2 kb, F) | MC | 0.353 | 0.080 | 0.167 | 1.170 | <0.001 |
| DN10097 | 2L:7597302-7597655 | R | 353 | *lncRNA:CR45370* (6.6 kb, F); *Spn28B* (14 kb, F) | SC | 0.002 | 0.100 | 0.000 | 0.826 | <0.001 |
| DN2695 | 2L:7601554-7603730 | R | 2176 | *lncRNA:CR45370* (0.5 kb, F); *Spn28B* (18 kb, F) | SC | 0.000 | 0.460 | 0.000 | 2.130 | <0.001 |

**Table S5.** Novel annotations of Sfps with 1-to-1-to-1 orthology among *mel*, *sim*, and *yak*. FBgn = Flybase gene number.

| FBgn *mel* | gene symbol *mel* | FBgn *sim* | FBgn *yak* |
| --- | --- | --- | --- |
| FBgn0000592 | *Est-6* | FBgn0012832 | FBgn0239133 |
| FBgn0003884 | *alphaTub84B* | FBgn0191393 | FBgn0068549 |
| FBgn0004629 | *Cys* | FBgn0191884 | FBgn0068171 |
| FBgn0011668 | *Mst57Da* | FBgn0270148 | FBgn0228508 |
| FBgn0011669 | *Mst57Db* | FBgn0268347 | FBgn0276545 |
| FBgn0011694 | *EbpII* | FBgn0196211 | FBgn0229194 |
| FBgn0015586 | *Acp76A* | FBgn0043398 | FBgn0239747 |
| FBgn0030589 | *CG9519* | FBgn0187177 | FBgn0233641 |
| FBgn0033868 | *S-Lap7* | FBgn0197024 | FBgn0231022 |
| FBgn0034471 | *Obp56e* | FBgn0183227 | FBgn0068110 |
| FBgn0036495 | *CG33259* | FBgn0186221 | FBgn0240340 |
| FBgn0043825 | *CG18284* | FBgn0250852 | FBgn0235891 |
| FBgn0050104 | *NT5E-2* | FBgn0183024 | FBgn0229466 |
| FBgn0051418 | *CG31418* | FBgn0190653 | FBgn0228001 |
| FBgn0051419 | *CG31419* | FBgn0190654 | FBgn0228003 |
| FBgn0051680 | *CG31680* | FBgn0270541 | FBgn0068381 |
| FBgn0051872 | *CG31872* | FBgn0195094 | FBgn0068366 |
| FBgn0052383 | *sphinx1* | FBgn0268539 | FBgn0237726 |
| FBgn0083965 | *CG34129* | FBgn0045605 | FBgn0228489 |
| FBgn0259958 | *Sfp24F* | FBgn0269971 | FBgn0275463 |
| FBgn0259961 | *CG42471* | FBgn0269795 | FBgn0276965 |
| FBgn0259964 | *Sfp33A3* | FBgn0269236 | FBgn0275933 |
| FBgn0259965 | *Sfp35C* | FBgn0268110 | FBgn0275630 |
| FBgn0259972 | *Sfp77F* | FBgn0269011 | FBgn0276285 |
| FBgn0262999 | *CG43307* | FBgn0270634 | FBgn0277370 |
| FBgn0053290 | *CG33290* | FBgn0186644 | FBgn0235434 |

