## Supplementary material for "Single-nucleus transcriptomes reveal functional and evolutionary properties of cell types in the Drosophila accessory gland": Dataset S10: README

gene X barcode expression matrices are contained in each directory in Matrix Market (MM) format. geneIDs.txt contains the row names - genes. barcodes.tsv 
contains the colnames - 10X cell barcodes. matrix.mtx contains the count data in MM format.

*_raw/ contain the raw output of HTSeq count for the cells of each species, and the 1000 sampled empty droplets
the first n columns are the cells, and the last 1000 barcodes are the empty droplets

*_clean/ contain the SoupX corrected counts for the cells of each species - some putative doublet cells also removed from raw - see SoupX in scripts/

*_ortho/ contain the counts for each species after SoupX correction, filtered to 1-to-1-to-1 orthologous genes

unannotated_v2_cleaned.tsv contains SoupX corrected counts for our unannotated Mel genes 
