## Supplementary material for "Single-nucleus transcriptomes reveal functional and evolutionary properties of cell types in the Drosophila accessory gland": Dataset S10: one_to_one_orthologs.R

```
# the following creates an ortholog table of all the 1:1:1 orthologs between
# melanogaster, simulans, and yakuba

library(dplyr)
library(purrr)
library(data.table)

# first just used grep to get out all lines of sim and yak orthologs from flybase ortholog tables, into two diff files

ortho_mel_yak <- read.table("ortho_mel_yak_2020_02.tsv", sep = "\t", quote = "", stringsAsFactors = T)
ortho_mel_sim <- read.table("ortho_mel_sim_2020_02.tsv", sep = "\t", quote = "", stringsAsFactors = T)

# keep only those entries that appear exactly once
ortho_mel_yak_one_to_one <- ortho_mel_yak[!(duplicated(ortho_mel_yak$V1) | duplicated(ortho_mel_yak$V1, fromLast = T)),] # this is a really clever peice of code from stack overflow
ortho_mel_yak_one_to_one <- ortho_mel_yak_one_to_one[!(duplicated(ortho_mel_yak_one_to_one$V6) | duplicated(ortho_mel_yak_one_to_one$V6, fromLast = T)),]

ortho_mel_sim_one_to_one <- ortho_mel_sim[!(duplicated(ortho_mel_sim$V1) | duplicated(ortho_mel_sim$V1, fromLast = T)),]
ortho_mel_sim_one_to_one <- ortho_mel_sim_one_to_one[!(duplicated(ortho_mel_sim_one_to_one$V6) | duplicated(ortho_mel_sim_one_to_one$V6, fromLast = T)),]

# join the tables to get one to one orthologs
ortho_mel_sim_yak_one_to_one <- inner_join(ortho_mel_sim_one_to_one, ortho_mel_yak_one_to_one, by = "V1")

# add headers
colnames(ortho_mel_sim_yak_one_to_one) <- c("FBgn_mel", "gene_symbol_mel", "scaffold_mel", "location_mel", "strand_mel", "FBgn_sim", "gene_symbol_sim", "scaffold_sim", "location_sim", "strand_sim", "orthoDB_ID_sim", "gene_symbol_yak", "scaffold_yak", "location_yak", "strand_yak", "FBgn_yak", "orthoDB_ID_yak")

ortho_mel_sim_yak_one_to_one <- ortho_mel_sim_yak_one_to_one[, c(1:17)]
names(ortho_mel_sim_yak_one_to_one)

write.table(ortho_mel_sim_yak_one_to_one, file = "ortho_MSY_1to1.tsv", quote = F, sep = "\t")
```
