## Supplementary material for "Single-nucleus transcriptomes reveal functional and evolutionary properties of cell types in the Drosophila accessory gland": Dataset S10: mel_sim_yak_seurat.R

```
##################################
## one to one ortholog analysis ##
##################################

# import the counts for each species
# these tables have been filtered for 1-to-1-orthologs, plus a smaller set of manually curated Sfps that are
# not technically 1-to-1-to-1 according to flybase annotation

#these counts for sim and yak also included some updated
# exon annotations from de novo transcriptome assembly to address differences in the annotation quality 
# from melanogaster
mel.counts <- read.table("mel_counts_ortho.tsv", row.names = 1, header = T)
sim.counts <- read.table("sim_counts_ortho.tsv", row.names = 1, header = T)
yak.counts <- read.table("yak_counts_ortho.tsv", row.names = 1, header = T)

# removing the SFP BG642312 as i suspect it has bad simulans annotation
mel.counts <- mel.counts[-grep("BG642312", rownames(mel.counts)),]
sim.counts <- sim.counts[-grep("BG642312", rownames(sim.counts)),]
yak.counts <- yak.counts[-grep("BG642312", rownames(yak.counts)),]

library(Seurat)
library(cowplot)
library(dplyr)
library(plotly)
library(ggplot2)
library(data.table)
library(magrittr)
library(plyr)

mel1.so <- CreateSeuratObject(counts = mel.counts, min.cells = 3, project = "msy_comparative")
sim.so <- CreateSeuratObject(counts = sim.counts, min.cells = 3, project = "msy_comparative")
yak.so <- CreateSeuratObject(counts = yak.counts, min.cells = 3, project = "msy_comparative")

# add species metadata
mel1.so <- AddMetaData(object = mel1.so, metadata = "mel", col.name = "species")
sim.so <- AddMetaData(object = sim.so, metadata = "sim", col.name = "species")
yak.so <- AddMetaData(object = yak.so, metadata = "yak", col.name = "species")

head(mel1.so@meta.data)
head(sim.so@meta.data)
head(yak.so@meta.data)

# plot num features, num counts
VlnPlot(mel1.so, features = c("nFeature_RNA", "nCount_RNA"), ncol = 3, cols = "slateblue")
VlnPlot(sim.so, features = c("nFeature_RNA", "nCount_RNA"), ncol = 3, cols = "slateblue")
VlnPlot(yak.so, features = c("nFeature_RNA", "nCount_RNA"), ncol = 3, cols = "slateblue")

# plot features v counts
FeatureScatter(mel1.so, feature1 = "nCount_RNA", feature2 = "nFeature_RNA", cols = "slateblue")
FeatureScatter(sim.so, feature1 = "nCount_RNA", feature2 = "nFeature_RNA", cols = "slateblue")
FeatureScatter(yak.so, feature1 = "nCount_RNA", feature2 = "nFeature_RNA", cols = "slateblue")

# Show 5% qunatiles for number of genes per cell
do.call("cbind", tapply(mel1.so@meta.data$nFeature_RNA,mel.so@active.ident,quantile,probs=seq(0,1,0.05)))
do.call("cbind", tapply(sim.so@meta.data$nFeature_RNA,sim.so@active.ident,quantile,probs=seq(0,1,0.05)))
do.call("cbind", tapply(yak.so@meta.data$nFeature_RNA,yak.so@active.ident,quantile,probs=seq(0,1,0.05)))
# Show 5% qunatiles for number of counts per cell per sample
do.call("cbind", tapply(mel1.so@meta.data$nCount_RNA,mel.so@active.ident,quantile,probs=seq(0,1,0.05)))
do.call("cbind", tapply(sim.so@meta.data$nCount_RNA,sim.so@active.ident,quantile,probs=seq(0,1,0.05)))
do.call("cbind", tapply(yak.so@meta.data$nCount_RNA,yak.so@active.ident,quantile,probs=seq(0,1,0.05)))

# plot cells per gene count
plot(sort(Matrix::rowSums(GetAssayData(mel1.so) >= 2)), xlab="gene rank", ylab="number of cells", main="Cells per genes ( >= 2 )")
plot(sort(Matrix::rowSums(GetAssayData(sim.so) >= 2)), xlab="gene rank", ylab="number of cells", main="Cells per genes ( >= 2 )")
plot(sort(Matrix::rowSums(GetAssayData(yak.so) >= 2)), xlab="gene rank", ylab="number of cells", main="Cells per genes ( >= 2 )")

median(mel1.so@meta.data$nCount_RNA)
median(sim.so@meta.data$nCount_RNA)
median(yak.so@meta.data$nCount_RNA)
median(mel1.so@meta.data$nFeature_RNA)
median(sim.so@meta.data$nFeature_RNA)
median(yak.so@meta.data$nFeature_RNA)

# now normalize the data
mel1.so <- NormalizeData(
  object = mel1.so,
  normalization.method = "LogNormalize",
  scale.factor = 10000)
sim.so <- NormalizeData(
  object = sim.so,
  normalization.method = "LogNormalize",
  scale.factor = 10000)
yak.so <- NormalizeData(
  object = yak.so,
  normalization.method = "LogNormalize",
  scale.factor = 10000)

# Gene selection for input to anchoring
mel1.so <- FindVariableFeatures(object = mel1.so, selection.method = "dispersion")
sim.so <- FindVariableFeatures(object = sim.so, selection.method = "dispersion")
yak.so <- FindVariableFeatures(object = yak.so, selection.method = "dispersion")

length(VariableFeatures(mel1.so))
head(VariableFeatures(mel1.so), 20)

length(VariableFeatures(sim.so))
head(VariableFeatures(sim.so), 20)

length(VariableFeatures(yak.so))
head(VariableFeatures(yak.so), 20)

# plot variable features with labels
top20 <- head(VariableFeatures(mel1.so), 20)
plot1 <- VariableFeaturePlot(mel1.so)
plot2 <- LabelPoints(plot = plot1, points = c(top20, "SP"))
plot2

top20 <- head(VariableFeatures(sim.so), 20)
plot1 <- VariableFeaturePlot(sim.so)
plot2 <- LabelPoints(plot = plot1, points = top20)
plot2

top20 <- head(VariableFeatures(yak.so), 20)
plot1 <- VariableFeaturePlot(yak.so)
plot2 <- LabelPoints(plot = plot1, points = top20)
plot2

# Scale the data (note this is absent from the Seurat tutorial however in the paper they state that you need to scale the data
# prior to performing integration)
mel1.so <- ScaleData(mel1.so)
sim.so <- ScaleData(sim.so)
yak.so <- ScaleData(yak.so)

### finishing each spp independently for marker gene calls
mel1.so <- RunPCA(object = mel1.so)
sim.so  <- RunPCA(object = sim.so)
yak.so  <- RunPCA(object = yak.so)

DimPlot(mel1.so)
DimPlot(sim.so)
DimPlot(yak.so)

mel1.so <- JackStraw(object = mel1.so, dims = 40)
mel1.so <- ScoreJackStraw(mel1.so, dims = 1:40)
JackStrawPlot(object = mel1.so, dims = 1:40)
use.pcs = c(1:8)
mel1.so <- RunUMAP(mel1.so, reduction = "pca", dims = use.pcs)
mel1.so <- FindNeighbors(mel1.so, reduction = "pca", dims = use.pcs)

sim.so <- JackStraw(object = sim.so, dims = 40)
sim.so <- ScoreJackStraw(sim.so, dims = 1:40)
JackStrawPlot(object = sim.so, dims = 1:40)
use.pcs = c(1:13)
sim.so <- RunUMAP(sim.so, reduction = "pca", dims = use.pcs)
sim.so <- FindNeighbors(sim.so, reduction = "pca", dims = use.pcs)

yak.so <- JackStraw(object = yak.so, dims = 40)
yak.so <- ScoreJackStraw(yak.so, dims = 1:40)
JackStrawPlot(object = yak.so, dims = 1:40)
use.pcs = c(1:8)
yak.so <- RunUMAP(yak.so, reduction = "pca", dims = use.pcs)
yak.so <- FindNeighbors(yak.so, reduction = "pca", dims = use.pcs)

mel1.so <- FindClusters(
  object = mel1.so,
  resolution = seq(0.2,1.4,0.1),
  verbose = FALSE)
sapply(grep("res",colnames(mel1.so@meta.data),value = TRUE),
       function(x) length(unique(mel1.so@meta.data[,x])))
Idents(mel1.so) <- "RNA_snn_res.0.2"

sim.so <- FindClusters(
  object = sim.so,
  resolution = seq(0.2,1.4,0.1),
  verbose = FALSE)
sapply(grep("res",colnames(sim.so@meta.data),value = TRUE),
       function(x) length(unique(sim.so@meta.data[,x])))
Idents(sim.so) <- "RNA_snn_res.0.2"

yak.so <- FindClusters(
  object = yak.so,
  resolution = seq(0.2,1.4,0.1),
  verbose = FALSE)
sapply(grep("res",colnames(yak.so@meta.data),value = TRUE),
       function(x) length(unique(yak.so@meta.data[,x])))
Idents(yak.so) <- "RNA_snn_res.0.2"

DimPlot(mel1.so, reduction = "umap", pt.size = 0.7) +
  theme_minimal(base_size = 15)
FeaturePlot(mel1.so, features = c("SP",
                                  "Acp36DE",
                                  "Acp32CD",
                                  "abd-A",
                                  "vvl",
                                  "Est-6"),
            reduction = "umap", pt.size = 0.7) +
  theme_minimal(base_size = 15)
mel1.so <- RenameIdents(mel1.so, "0" = "MC",
                                 "1" = "MC",
                                 "3" = "SC",
                                 "2" = "EDC")

DimPlot(sim.so, reduction = "umap", pt.size = 0.7) +
  theme_minimal(base_size = 15)
FeaturePlot(sim.so, features = c("SP",
                                  "Acp36DE",
                                  "Acp32CD",
                                  "abd-A",
                                  "vvl",
                                  "Est-6"),
            reduction = "umap", pt.size = 0.7) +
  theme_minimal(base_size = 15)
sim.so <- RenameIdents(sim.so, "0" = "MC",
                               "1" = "EDC",
                               "2" = "MC",
                               "3" = "SC")

DimPlot(yak.so, reduction = "umap", pt.size = 0.7) +
  theme_minimal(base_size = 15)
FeaturePlot(yak.so, features = c("SP",
                                  "Acp36DE",
                                  "Acp32CD",
                                  "abd-A",
                                  "vvl",
                                  "Est-6"),
            reduction = "umap", pt.size = 0.7) +
  theme_minimal(base_size = 15)
yak.so <- RenameIdents(yak.so, "0" = "MC",
                               "1" = "MC",
                               "2" = "EDC",
                               "3" = "SC")

### find independent markers for each species
markers_mel_ortho <- FindAllMarkers(mel1.so, min.pct = 0.25, logfc.threshold = 0.25, only.pos = T)
markers_mel_ortho <- subset(markers_mel_ortho, p_val_adj < 0.05)
markers_sim_ortho <- FindAllMarkers(sim.so, min.pct = 0.25, logfc.threshold = 0.25, only.pos = T)
markers_sim_ortho <- subset(markers_sim_ortho, p_val_adj < 0.05)
markers_yak_ortho <- FindAllMarkers(yak.so, min.pct = 0.25, logfc.threshold = 0.25, only.pos = T)
markers_yak_ortho <- subset(markers_yak_ortho, p_val_adj < 0.05)


############################################
########### INTEGRATED DATA ################
############################################

# integrate the data
anchors <- FindIntegrationAnchors(object.list = c(mel1.so, sim.so, yak.so), dims = 1:20)
msy.so <- IntegrateData(anchorset = anchors, dims = 1:20)

# now analyze combined data!
DefaultAssay(msy.so) <- "integrated"

# Run the standard workflow for visualization and clustering
msy.so <- ScaleData(object = msy.so)
msy.so <- RunPCA(object = msy.so)

DimPlot(msy.so, dims = c(1,2), reduction = "pca", cols = c("slateblue", "firebrick", "aquamarine"), group.by = "species")
DimPlot(msy.so, dims = c(1,3), reduction = "pca", cols = c("slateblue", "firebrick", "aquamarine"), group.by = "species")

ElbowPlot(msy.so)

# use JackStraw to determine significance of PCs
msy.so <- JackStraw(object = msy.so, dims = 40)
msy.so <- ScoreJackStraw(msy.so, dims = 1:40)
JackStrawPlot(object = msy.so, dims = 1:40)
use.pcs = c(1:14)

# UMAP + t-SNE and Clustering
msy.so <- RunUMAP(msy.so, reduction = "pca", dims = use.pcs)
msy.so <- FindNeighbors(msy.so, reduction = "pca", dims = use.pcs)

# look at cluster nums at a series of resolutions
msy.so <- FindClusters(
  object = msy.so,
  resolution = seq(0.2,1.4,0.1),
  verbose = FALSE)
sapply(grep("res",colnames(msy.so@meta.data),value = TRUE),
       function(x) length(unique(msy.so@meta.data[,x])))
# choose resolution
Idents(msy.so) <- "integrated_snn_res.0.3"

# UMAP Visualization

p1 <- DimPlot(msy.so, reduction = "umap", group.by = "species", pt.size = 0.05, shuffle = T) +
        scale_color_brewer(palette = "Paired") + theme(plot.title = element_text(size = 9), 
                                                     axis.title.x = element_text(size = 9),
                                                     axis.title.y = element_text(size = 9), 
                                                      legend.text = element_text(size = 9),
                                                        axis.text = element_text(size = 9))
p1[[1]]$layers[[1]]$aes_params$alpha = 0.7
p2 <- DimPlot(msy.so, reduction = "umap", pt.size = 0.05, cols = my_colors[c(1,3,2)]) + theme(plot.title = element_text(size = 9), 
                                                                                           axis.title.x = element_text(size = 9),
                                                                                           axis.title.y = element_text(size = 9),
                                                                                            legend.text = element_text(size = 9),
                                                                                              axis.text = element_text(size = 9))
p2[[1]]$layers[[1]]$aes_params$alpha = 0.7
plot_grid(p1, p2, labels = c('A', 'B'))

plot_grid(mel.umap, p1, p2, labels = c('A', 'B', 'C'), nrow = 1)

plot(p1)
plot(p2)

DimPlot(msy.so, reduction = "umap", split.by = "species", pt.size = 0.7) +
    theme_minimal(base_size = 15)
FeaturePlot(msy.so, features = c('nCount_RNA'), pt.size=0.7, split.by = "species", label = T)

# find markers
markers_all <- FindAllMarkers(
  object = msy.so, 
  only.pos = TRUE, 
  min.pct = 0.25, 
  thresh.use = 0.25
)

dim(markers_all)
head(markers_all)

markers_all_significant <- subset(markers_all, markers_all$p_val_adj < 0.05)
markers_all_single <- markers_all_significant[markers_all_significant$gene %in% names(table(markers_all_significant$gene))[table(markers_all_significant$gene) == 1],]

# subclusters
msy.so %<>% RenameIdents('0' = 'MCsp1',
                         '1' = 'MCsp2',
                         '2' = 'EDC',
                         '3' = 'SC',
                         '4' = 'MCsp3')

subpops_umap_msy <- DimPlot(msy.so, reduction = "umap", split.by = "species", pt.size = 0.1, shuffle = T,
                            cols = c(my_colors[1], "salmon", my_colors[3:2], "gold2")) +
                                             theme(plot.tag = element_text(size = 10, face = "plain"), 
                                                 axis.title.x = element_text(size = 10),
                                                 axis.title.y = element_text(size = 10), 
                                                  legend.text = element_text(size = 10),
                                                 axis.text.x = element_text(size = 10),
                                                 axis.text.y = element_text(size = 10) 
                                                  )
subpops_umap_msy[[1]]$layers[[1]]$aes_params$alpha = 0.7
subpops_umap_msy

markers_MC_subpops_MSY <- FindMarkers(msy.so, 'MCsp2', 'MCsp1')
markers_MC_subpops_MSY <- subset(markers_MC_subpops_MSY, p_val_adj <= 0.05)
markers_MC_subpops_MSY %<>% rownames_to_column("gene")

markers_MC_subpop4_MSY <- FindMarkers(msy.so, 'MCsp3', c('MCsp1','MCsp1'))
markers_MC_subpop4_MSY <- subset(markers_MC_subpop4_MSY, p_val_adj <= 0.05)
markers_MC_subpop4_MSY %<>% rownames_to_column("gene")

# change assay to RNA for vizualizing counts
DefaultAssay(msy.so) <- "integrated"

top5 <- markers_all_significant %>% group_by(cluster) %>% top_n(5, avg_logFC)
dim(top5)

msy.so <- ScaleData(msy.so)

DoHeatmap(
  object = msy.so, 
  features = top5$gene
) 

## save the unbiased "subpopulations" for a later analysis
msy_subpops.so <- msy.so

# look at marker genes
FeaturePlot(msy.so, 
            reduction = "umap", 
            pt.size = 1, 
            features = c("vvl"),
            split.by = "species")
FeaturePlot(msy.so, 
            reduction = "umap", 
            pt.size = 1.5, 
            features = c("abd-A"),
            split.by = "species")
FeaturePlot(msy.so, 
            reduction = "umap", 
            pt.size = 1.5, 
            features = c("Acp36DE"),
            split.by = "species")
FeaturePlot(msy.so, 
            reduction = "umap", 
            pt.size = 1.5, 
            features = c("mfas"),
            split.by = "species")
FeaturePlot(msy.so, 
            reduction = "umap", 
            pt.size = 1.5, 
            features = c("SP"),
            split.by = "species")

## merge our cells together for downstream analysis of markers
msy.so <- RenameIdents(
  object = msy.so,
  'MCsp1' = 'MC', 'MCsp2' = 'MC', 'MCsp3' = 'MC')

table(Idents(msy.so))
msy.so$CellType <- Idents(msy.so)

### get numbers of cells in cell types by species
table(Idents(subset(msy.so, species == "mel")))
table(Idents(subset(msy.so, species == "sim")))
table(Idents(subset(msy.so, species == "yak")))

### extract the indices of different cell types for trinity de novo assembly ##
###
##
#

MC_mel <- rownames(subset(msy.so@meta.data, CellType == 'main cells' & species == 'mel'))
MC_sim <- rownames(subset(msy.so@meta.data, CellType == 'main cells' & species == 'sim'))
MC_yak <- rownames(subset(msy.so@meta.data, CellType == 'main cells' & species == 'yak'))
SC_mel <- rownames(subset(msy.so@meta.data, CellType == 'secondary cells' & species == 'mel'))
SC_sim <- rownames(subset(msy.so@meta.data, CellType == 'secondary cells' & species == 'sim'))
SC_yak <- rownames(subset(msy.so@meta.data, CellType == 'secondary cells' & species == 'yak'))
EDC_mel <- rownames(subset(msy.so@meta.data, CellType == 'ejaculatory duct cells' & species == 'mel'))
EDC_sim <- rownames(subset(msy.so@meta.data, CellType == 'ejaculatory duct cells' & species == 'sim'))
EDC_yak <- rownames(subset(msy.so@meta.data, CellType == 'ejaculatory duct cells' & species == 'yak'))

barcode_celltype_index <- list(MC_mel, MC_sim, MC_yak, 
                               SC_mel, SC_sim, SC_yak, 
                               EDC_mel, EDC_sim, EDC_yak 
                            )
names(barcode_celltype_index) <- c("MC_mel", "MC_sim", "MC_yak",
                                   "SC_mel", "SC_sim", "SC_yak",
                                   "EDC_mel", "EDC_sim", "EDC_yak")

library(readr)
lapply(names(barcode_celltype_index), 
       function(x) write_lines(barcode_celltype_index[[x]], 
                               path = paste("~/Documents/Begun/STAR_counts/de_novo/", 
                                            x, ".txt", 
                                            sep = ""), 
                               sep = "\n"))

# run FindAllMarkers on the merged cell types
markers_merged <- FindAllMarkers(
  object = msy.so, 
  only.pos = TRUE, 
  min.pct = 0.25, 
  thresh.use = 0.25
)

dim(markers_merged)
head(markers_merged)

markers_merged_significant <- subset(markers_merged, markers_merged$p_val_adj < 0.05)
markers_merged_single <- markers_merged_significant[markers_merged_significant$gene %in% names(table(markers_merged_significant$gene))[table(markers_merged_significant$gene) == 1],]

top15 <- markers_merged_significant %>% group_by(cluster) %>% top_n(15, avg_logFC)
dim(top15)

heatmap <- DoHeatmap(object = msy.so, features = top15$gene, label = F, raster = F) 
plot(heatmap)
ggplot2::ggsave(filename = "~/Documents/Begun/STAR_counts/figs/publication_figures/MSY_heatmap.pdf",
                plot = heatmap, height = 7, width = 8)

write.table(x = markers_merged_significant, file = "~/Documents/Begun/STAR_counts/MSY_markers_12_20.tsv", quote = F, row.names = F, sep = "\t")
write.table(x = markers_merged_single, file = "~/Documents/Begun/STAR_counts/MSY_markers_single_12_20.tsv", quote = F, row.names = F, sep = "\t")

###################################
## DE across species using Limma ##
###################################

# note - MUST set default assay to RNA for the DE to work across integrated dataset 

DefaultAssay(msy.so) <- "RNA"

library(limma)
main.so <- subset(msy.so, idents = 'MC')
secondary.so <- subset(msy.so, idents = 'SC')
ED.so <- subset(msy.so, idents = 'EDC')

main.expr <- as.matrix(GetAssayData(main.so))
secondary.expr <- as.matrix(GetAssayData(secondary.so))
ED.expr <- as.matrix(GetAssayData(ED.so))

# Filter out genes that are 0 for every cell in this cluster
main.zeroes <- which(rowSums(main.expr) == 0)
main.expr <- main.expr[-main.zeroes,]
secondary.zeroes <- which(rowSums(secondary.expr) == 0)
secondary.expr <- secondary.expr[-secondary.zeroes,]
ED.zeroes <- which(rowSums(ED.expr) == 0)
ED.expr <- ED.expr[-ED.zeroes,]

# create limma matrix model
main.mm <- model.matrix(~0 + species, data = main.so@meta.data)
main.fit <- lmFit(main.expr, main.mm)  
head(coef(main.fit), 50) # means in each sample for each gene
secondary.mm <- model.matrix(~0 + species, data = secondary.so@meta.data)
secondary.fit <- lmFit(secondary.expr, secondary.mm)  
head(coef(secondary.fit), 50) # means in each sample for each gene
ED.mm <- model.matrix(~0 + species, data = ED.so@meta.data)
ED.fit <- lmFit(ED.expr, ED.mm)  
head(coef(ED.fit), 50) # means in each sample for each gene

# create the set of contrasts in expression across species
main.contr <- makeContrasts(speciesmel - speciessim, speciesmel - speciesyak, speciessim-speciesyak, levels = colnames(coef(main.fit)))
main.tmp <- contrasts.fit(main.fit, contrasts = main.contr)
main.tmp <- eBayes(main.tmp)
main.DE <- topTable(main.tmp, sort.by = "F", number = Inf) # top 50 DE genes
colnames(main.tmp)
head(main.DE)

# create subset datatables for each pairwise contrast
main.melvsim.DE <- topTable(main.tmp, coef = 1, sort.by = "P", number = Inf)
main.melvyak.DE <- topTable(main.tmp, coef = 2, sort.by = "P", number = Inf)
main.simvyak.DE <- topTable(main.tmp, coef = 3, sort.by = "P", number = Inf)

# create the set of contrasts in expression across species
secondary.contr <- makeContrasts(speciesmel - speciessim, speciesmel - speciesyak, speciessim-speciesyak, levels = colnames(coef(secondary.fit)))
secondary.tmp <- contrasts.fit(secondary.fit, contrasts = secondary.contr)
secondary.tmp <- eBayes(secondary.tmp)
secondary.DE <- topTable(secondary.tmp, sort.by = "F", number = Inf) # top 50 DE genes
head(secondary.DE)

# create subset datatables for each pairwise contrast
secondary.melvsim.DE <- topTable(secondary.tmp, coef = 1, sort.by = "P", number = Inf)
secondary.melvyak.DE <- topTable(secondary.tmp, coef = 2, sort.by = "P", number = Inf)
secondary.simvyak.DE <- topTable(secondary.tmp, coef = 3, sort.by = "P", number = Inf)

# create the set of contrasts in expression across species
ED.contr <- makeContrasts(speciesmel - speciessim, speciesmel - speciesyak, speciessim - speciesyak, levels = colnames(coef(ED.fit)))
ED.tmp <- contrasts.fit(ED.fit, contrasts = ED.contr)
ED.tmp <- eBayes(ED.tmp)
ED.DE <- topTable(ED.tmp, sort.by = "F", number = Inf) # top 50 DE genes
head(ED.DE)

# create subset datatables for each pairwise contrast
ED.melvsim.DE <- topTable(ED.tmp, coef = 1, sort.by = "P", number = Inf)
ED.melvyak.DE <- topTable(ED.tmp, coef = 2, sort.by = "P", number = Inf)
ED.simvyak.DE <- topTable(ED.tmp, coef = 3, sort.by = "P", number = Inf)

# subset out the significantly DE genes
main.DE.sig <- subset(main.DE, main.DE$adj.P.Val < 0.05)
secondary.DE.sig <- subset(secondary.DE, secondary.DE$adj.P.Val < 0.05)
ED.DE.sig <- subset(ED.DE, ED.DE$adj.P.Val < 0.05)

main.DE.melvsim.sig <- subset(main.melvsim.DE, main.melvsim.DE$adj.P.Val < 0.05 & abs(main.melvsim.DE$logFC) > 1)
main.DE.melvyak.sig <- subset(main.melvyak.DE, main.melvyak.DE$adj.P.Val < 0.05 & abs(main.melvyak.DE$logFC) > 1)
main.DE.simvyak.sig <- subset(main.simvyak.DE, main.simvyak.DE$adj.P.Val < 0.05 & abs(main.simvyak.DE$logFC) > 1)

secondary.DE.melvsim.sig <- subset(secondary.melvsim.DE, secondary.melvsim.DE$adj.P.Val < 0.05 & abs(secondary.melvsim.DE$logFC) > 1)
secondary.DE.melvyak.sig <- subset(secondary.melvyak.DE, secondary.melvyak.DE$adj.P.Val < 0.05 & abs(secondary.melvyak.DE$logFC) > 1)
secondary.DE.simvyak.sig <- subset(secondary.simvyak.DE, secondary.simvyak.DE$adj.P.Val < 0.05 & abs(secondary.simvyak.DE$logFC) > 1)

ED.DE.melvsim.sig <- subset(ED.melvsim.DE, ED.melvsim.DE$adj.P.Val < 0.05 & abs(ED.melvsim.DE$logFC) > 1)
ED.DE.melvyak.sig <- subset(ED.melvyak.DE, ED.melvyak.DE$adj.P.Val < 0.05 & abs(ED.melvyak.DE$logFC) > 1)
ED.DE.simvyak.sig <- subset(ED.simvyak.DE, ED.simvyak.DE$adj.P.Val < 0.05 & abs(ED.simvyak.DE$logFC) > 1)

# how many DE genes per cell type ?
melvsim <- c(
  100*(length(rownames(main.DE.melvsim.sig))/length(rownames(main.expr))),
  100*(length(rownames(secondary.DE.melvsim.sig))/length(rownames(secondary.expr))),
  100*(length(rownames(ED.DE.melvsim.sig))/length(rownames(ED.expr)))
)

melvyak <- c( 
  100*(length(rownames(main.DE.melvyak.sig))/length(rownames(main.expr))),
  100*(length(rownames(secondary.DE.melvyak.sig))/length(rownames(secondary.expr))),
  100*(length(rownames(ED.DE.melvyak.sig))/length(rownames(ED.expr)))
)

simvyak <- c(
  100*(length(rownames(main.DE.simvyak.sig))/length(rownames(main.expr))),
  100*(length(rownames(secondary.DE.simvyak.sig))/length(rownames(secondary.expr))),
  100*(length(rownames(ED.DE.simvyak.sig))/length(rownames(ED.expr)))
)

cell <- rep(c("MC", "SC", "EDC"), 3)
comparison <- c(rep("melvsim", 3), rep("melvyak", 3), rep("simvyak", 3))
percent_genes_DE <- c(melvsim, melvyak, simvyak)

DE_ratios <- data.frame(cell, comparison, percent_genes_DE)
DE_ratios

# factor cell types to order them in the bar plot
DE_ratios$cell <- factor(DE_ratios$cell, levels = c("MC", "SC", "EDC"))

DE_ratio_plot <- ggplot(DE_ratios, aes(comparison, percent_genes_DE, fill = cell)) + 
  geom_bar(position="dodge", stat="identity") +
  scale_fill_manual(values = my_colors) +
  theme_minimal(base_size =  11) +
  labs(y = "percent genes differentially expressed",
       fill = "cell type") +
  theme(axis.title.x = element_blank()) +
  scale_x_discrete(labels = c("mel vs. sim", "mel vs. yak", "sim vs. yak"))

DE_ratio_plot

DE_ratio_plot_125 <- DE_ratio_plot
DE_ratio_plot_150 <- DE_ratio_plot
DE_ratio_plot_050 <- DE_ratio_plot
DE_ratio_plot_100 <- DE_ratio_plot

plot_grid(DE_ratio_plot_050 + theme_minimal(base_size = 10) + theme(axis.title.x = element_blank(), legend.position = "none", axis.text.x = element_blank()), 
          DE_ratio_plot_100 + theme_minimal(base_size = 10) + theme(axis.title.x = element_blank(), legend.position = "none", axis.text.x = element_blank(), axis.title.y = element_blank()), 
          DE_ratio_plot_125 + theme_minimal(base_size = 10) + theme(axis.title.x = element_blank(), legend.position = "none", axis.text.x = element_text(size = 10)), 
          DE_ratio_plot_150 + theme_minimal(base_size = 10) + theme(axis.title.x = element_blank(), legend.position = "none", axis.title.y = element_blank(), axis.text.x = element_text(size = 10)))

### directionality of DE?
hist(main.DE.sig$speciesmel...speciessim, breaks = 100)
summary(main.DE.sig$speciesmel...speciessim)
hist(secondary.DE.sig$speciesmel...speciessim, breaks = 100)
summary(secondary.DE.sig$speciesmel...speciessim)
hist(ED.DE.sig$speciesmel...speciessim, breaks = 100)
summary(ED.DE.sig$speciesmel...speciessim)
hist(main.DE.sig$speciesmel...speciesyak, breaks = 100)
summary(main.DE.sig$speciesmel...speciesyak)
hist(secondary.DE.sig$speciesmel...speciesyak, breaks = 100)
summary(secondary.DE.sig$speciesmel...speciesyak)
hist(ED.DE.sig$speciesmel...speciesyak, breaks = 100)
summary(ED.DE.sig$speciesmel...speciesyak)
hist(main.DE.sig$speciessim...speciesyak, breaks = 100)
summary(main.DE.sig$speciessim...speciesyak)
hist(secondary.DE.sig$speciessim...speciesyak, breaks = 100)
summary(secondary.DE.sig$speciessim...speciesyak)
hist(ED.DE.sig$speciessim...speciesyak, breaks = 100)
summary(ED.DE.sig$speciessim...speciesyak)


# GTest
main.DEvNon.melvsim <- c(length(rownames(main.DE.melvsim.sig)), length(rownames(main.expr)) - length(rownames(main.DE.melvsim.sig)))
secondary.DEvNon.melvsim <- c(length(rownames(secondary.DE.melvsim.sig)), length(rownames(secondary.expr)) - length(rownames(secondary.DE.melvsim.sig)))
ED.DEvNon.melvsim <- c(length(rownames(ED.DE.melvsim.sig)), length(rownames(ED.expr)) - length(rownames(ED.DE.melvsim.sig)))
fraction.DE.table.melvsim <- data.frame(main.DEvNon.melvsim, secondary.DEvNon.melvsim, ED.DEvNon.melvsim, row.names = c("DE", "non"))
fraction.DE.table.melvsim <- t(fraction.DE.table.melvsim)

main.DEvNon.melvyak <- c(length(rownames(main.DE.melvyak.sig)), length(rownames(main.expr)) - length(rownames(main.DE.melvyak.sig)))
secondary.DEvNon.melvyak <- c(length(rownames(secondary.DE.melvyak.sig)), length(rownames(secondary.expr)) - length(rownames(secondary.DE.melvyak.sig)))
ED.DEvNon.melvyak <- c(length(rownames(ED.DE.melvyak.sig)), length(rownames(ED.expr)) - length(rownames(ED.DE.melvyak.sig)))
fraction.DE.table.melvyak <- data.frame(main.DEvNon.melvyak, secondary.DEvNon.melvyak, ED.DEvNon.melvyak, row.names = c("DE", "non"))
fraction.DE.table.melvyak <- t(fraction.DE.table.melvyak)

main.DEvNon.simvyak <- c(length(rownames(main.DE.simvyak.sig)), length(rownames(main.expr)) - length(rownames(main.DE.simvyak.sig)))
secondary.DEvNon.simvyak <- c(length(rownames(secondary.DE.simvyak.sig)), length(rownames(secondary.expr)) - length(rownames(secondary.DE.simvyak.sig)))
ED.DEvNon.simvyak <- c(length(rownames(ED.DE.simvyak.sig)), length(rownames(ED.expr)) - length(rownames(ED.DE.simvyak.sig)))
fraction.DE.table.simvyak <- data.frame(main.DEvNon.simvyak, secondary.DEvNon.simvyak, ED.DEvNon.simvyak, row.names = c("DE", "non"))
fraction.DE.table.simvyak <- t(fraction.DE.table.simvyak)

DE_table <- (rbind(fraction.DE.table.melvsim, fraction.DE.table.melvyak, fraction.DE.table.simvyak))
rownames(DE_table) <- c("ms_MC", "ms_SC", "ms_ED", "my_MC", "my_SC", "my_ED", "sy_MC", "sy_SC", "sy_ED")

DE_table

library("RVAideMemoire")
G.test(DE_table)

# logFC = 1
#G = 76.047, df = 8, p-value = 3.043e-13
DE_table_G_test_1 <- pairwise.G.test(DE_table, p.method = "fdr")
DE_table_G_test_1

# logFC = 1.25
#G = 77.133, df = 8, p-value = 1.843e-13
DE_table_G_test_125 <- pairwise.G.test(DE_table, p.method = "fdr")
DE_table_G_test_125

# logFC = 1.5
# G = 80.636, df = 8, p-value = 3.64e-14
DE_table_G_test_150 <- pairwise.G.test(DE_table, p.method = "fdr")
DE_table_G_test_150

# logFC = 0.5
# G = 80.636, df = 8, p-value = 3.64e-14
DE_table_G_test_150 <- pairwise.G.test(DE_table, p.method = "fdr")
DE_table_G_test_150


###
# look at fold change distributions

# main
FC_main <- rbind(main.DE.melvsim.sig,
                 main.DE.melvyak.sig,
                 main.DE.simvyak.sig
               )
FC_main %<>% cbind(CellType = rep("MC", length(rownames(FC_main))))
FC_main %<>% cbind(Comparison = c(rep("mel_sim", length(rownames(main.DE.melvsim.sig))), 
                                  rep("mel_yak", length(rownames(main.DE.melvyak.sig))), 
                                  rep("sim_yak", length(rownames(main.DE.simvyak.sig)))
                                  )
                   )

# secondary
FC_secondary <- rbind(secondary.DE.melvsim.sig,
                      secondary.DE.melvyak.sig,
                      secondary.DE.simvyak.sig
                  )
FC_secondary %<>% cbind(CellType = rep("SC", length(rownames(FC_secondary))))
FC_secondary %<>% cbind(Comparison = c(rep("mel_sim", length(rownames(secondary.DE.melvsim.sig))), 
                                      rep("mel_yak", length(rownames(secondary.DE.melvyak.sig))), 
                                      rep("sim_yak", length(rownames(secondary.DE.simvyak.sig)))
                                      )
                      )

# ejaculatory duct
FC_ED <- rbind(ED.DE.melvsim.sig,
               ED.DE.melvyak.sig,
               ED.DE.simvyak.sig
                ) 
FC_ED %<>% cbind(CellType = rep("ED", length(rownames(FC_ED))))
FC_ED %<>% cbind(Comparison = c(rep("mel_sim", length(rownames(ED.DE.melvsim.sig))), 
                                rep("mel_yak", length(rownames(ED.DE.melvyak.sig))), 
                                rep("sim_yak", length(rownames(ED.DE.simvyak.sig)))
                                )
                  )
# bind all together
FC_all <- rbind(FC_main, FC_secondary, FC_ED)

p1 <- ggplot(data = FC_all, aes(x = abs(logFC))) +
        geom_density(aes(color = CellType), alpha = 0.2, fill = "lightgrey") +
        scale_color_manual(values = my_colors) +
        theme_minimal(base_size = 9) + 
        labs(color= "cell type")

p2 <- ggplot(data = FC_all, aes(x = abs(logFC))) +
        geom_density(aes(color = Comparison), alpha = 0.2, fill = "lightgrey") +
       # scale_fill_manual(values = my_colors) +
        theme_minimal(base_size = 9) + 
        scale_color_brewer(palette = "Set2") +
        labs(color = "comparison")
plot_grid(p1, p2)

kruskal.test(logFC ~ interaction(CellType, Comparison), data = FC_all)
# Kruskal-Wallis chi-squared = 2.2601, df = 8, p-value = 0.972
kruskal.test(logFC ~ CellType, data = FC_all)
# Kruskal-Wallis chi-squared = 0.6925, df = 2, p-value = 0.7073
kruskal.test(logFC ~ Comparison, data = FC_all)
# Kruskal-Wallis chi-squared = 0.87377, df = 2, p-value = 0.646

# none of the logFC distributions are sig different 


### concerted vs independent cell-type evolution
# we want to know how frequently DE is shared across cells or if it generally occurs in just one
# cell-type

main.DE <- rownames_to_column(main.DE, "gene")
secondary.DE <- rownames_to_column(secondary.DE, "gene")
ED.DE <- rownames_to_column(ED.DE, "gene")

main.DE.sig <- rownames_to_column(main.DE.sig, "gene")
secondary.DE.sig <- rownames_to_column(secondary.DE.sig, "gene")
ED.DE.sig <- rownames_to_column(ED.DE.sig, "gene")

MC.DE.1 <- subset(main.DE.sig, 
                  abs(speciesmel...speciessim) >= 1 |
                  abs(speciesmel...speciesyak) >= 1 |
                  abs(speciessim...speciesyak) >= 1)$gene
SC.DE.1 <- subset(secondary.DE.sig, 
                  abs(speciesmel...speciessim) >= 1 |
                  abs(speciesmel...speciesyak) >= 1 |
                  abs(speciessim...speciesyak) >= 1)$gene
ED.DE.1 <- subset(ED.DE.sig, 
                  abs(speciesmel...speciessim) >= 1 |
                  abs(speciesmel...speciesyak) >= 1 |
                  abs(speciessim...speciesyak) >= 1)$gene

length(c(MC.DE.1, SC.DE.1, ED.DE.1))
# 459 total DE genes 
length(unique(c(MC.DE.1, SC.DE.1, ED.DE.1)))
# 358 unique genes

# what is the frequency of occurances in 1, 2, or 3 cell types?
table(table(c(MC.DE.1, SC.DE.1, ED.DE.1)))
#  1   2   3 
# 282  51  25

282 / 358
51 /  358
25 /  358

# so 78% of genes DE at logFC >= 1 are so in obly 1 cell type, 16% are in 2, and 7% are in 3

## the genes DE in all three 
triplets <- names(table(c(MC.DE.1, SC.DE.1, ED.DE.1))[table(c(MC.DE.1, SC.DE.1, ED.DE.1)) == 3])

VlnPlot(msy.so, features = triplets, split.by = "species",  cols = my_colors)

# combine all genes w. DE in at least one cell type
all.DE.1 <- unique(c(MC.DE.1, SC.DE.1, ED.DE.1))

MC.DE.0.5 <- subset(main.DE.sig, 
                  abs(speciesmel...speciessim) >= 0.5 |
                  abs(speciesmel...speciesyak) >= 0.5 |
                  abs(speciessim...speciesyak) >= 0.5)$gene
SC.DE.0.5 <- subset(secondary.DE.sig, 
                  abs(speciesmel...speciessim) >= 0.5 |
                  abs(speciesmel...speciesyak) >= 0.5 |
                  abs(speciessim...speciesyak) >= 0.5)$gene
ED.DE.0.5 <- subset(ED.DE.sig, 
                  abs(speciesmel...speciessim) >= 0.5 |
                  abs(speciesmel...speciesyak) >= 0.5 |
                  abs(speciessim...speciesyak) >= 0.5)$gene
all.DE.0.5 <- c(MC.DE.0.5, SC.DE.0.5, ED.DE.0.5) # all total occurances of DE genes at logFC=0.5 cutoff 

# get the genes at logFC >= 1 that occur just in one cell type
singleton_DE.1 <- all.DE.1[table(c(MC.DE.1, SC.DE.1, ED.DE.1)) == 1]

# how many times do we see these singletons once the logFC filter is dropped down to 0.5?
table(table(all.DE.0.5[all.DE.0.5 %in% singleton_DE.1]))
#   1   2   3 
#   143  80  60

143 / (143+80+60)
# so 51% of the singleton genes at logFC >= 1 are still only DE in one cell type at logFC >= 0.5

dim(subset(main.DE, gene %in% all.DE.1))
dim(subset(secondary.DE, gene %in% all.DE.1))
dim(subset(ED.DE, gene %in% all.DE.1))

# what are the given genes that do NOT show up in a particular set of all DE by cell-type
all.DE.1[!all.DE.1 %in% subset(ED.DE, gene %in% all.DE.1)$gene]
all.DE.1[!all.DE.1 %in% subset(main.DE, gene %in% all.DE.1)$gene]
all.DE.1[!all.DE.1 %in% subset(secondary.DE, gene %in% all.DE.1)$gene]

###########################################################################################################
###########################################################################################################
###########################################################################################################
#### permutation of logFC values - want to ask what the null distribution of coincidence of logFC changes is
#### across cells

all.DE.1 <- setNames(data.frame(unique(c(MC.DE.1, SC.DE.1, ED.DE.1))), "gene") # combine all genes w. DE logFC > 1 in at least one cell type
all.DE.0.5 <- setNames(data.frame(unique(c(MC.DE.0.5, SC.DE.0.5, ED.DE.0.5))), "gene") # all total occurances of DE logFC > 0.5 in at least one cell type

DE_list <- list(all.DE.1, 
                subset(main.DE, gene %in% all.DE.1$gene),
                subset(secondary.DE, gene %in% all.DE.1$gene),
                subset(ED.DE, gene %in% all.DE.1$gene))
all.DE.1 <- setNames(join_all(DE_list, by = "gene", type = "inner")[,c(1:5,9:12,16:19)],
                             c("gene", "MC_ms", "MC_my", "MC_sy", "avg_exp_MC",
                                       "SC_ms", "SC_my", "SC_sy", "avg_exp_SC",
                                       "EDC_ms", "EDC_my", "EDC_sy", "avg_exp_EDC"))

DE_list <- list(all.DE.0.5, 
                subset(main.DE, gene %in% all.DE.0.5$gene),
                subset(secondary.DE, gene %in% all.DE.0.5$gene),
                subset(ED.DE, gene %in% all.DE.0.5$gene))
all.DE.0.5 <- setNames(join_all(DE_list, by = "gene", type = "inner")[,c(1:5,9:12,16:19)],
                             c("gene", "MC_ms", "MC_my", "MC_sy", "avg_exp_MC",
                               "SC_ms", "SC_my", "SC_sy", "avg_exp_SC",
                               "EDC_ms", "EDC_my", "EDC_sy", "avg_exp_EDC"))

pearson_results <- data.frame(
                      c("MC_SC", "MC_EDC", "SC_EDC"),
                      c(cor(all.DE.1$MC_ms, all.DE.1$SC_ms, method = 'pearson'),
                           cor(all.DE.1$MC_ms, all.DE.1$EDC_ms, method = 'pearson'),
                           cor(all.DE.1$SC_ms, all.DE.1$EDC_ms, method = 'pearson')),
                      c(cor(all.DE.1$MC_my, all.DE.1$SC_my, method = 'pearson'),
                           cor(all.DE.1$MC_my, all.DE.1$EDC_my, method = 'pearson'),
                           cor(all.DE.1$SC_my, all.DE.1$EDC_my, method = 'pearson')),
                      c(cor(all.DE.1$MC_sy, all.DE.1$SC_sy, method = 'pearson'),
                           cor(all.DE.1$MC_sy, all.DE.1$EDC_sy, method = 'pearson'),
                           cor(all.DE.1$SC_sy, all.DE.1$EDC_sy, method = 'pearson'))
                      )

pearson_results %<>% setNames(c("cell_type_comparison", 
                                "mel_sim",
                                "mel_yak",
                                "sim_yak"))

pearson_results

pearson_results_05 <- data.frame(
  c("MC_SC", "MC_EDC", "SC_EDC"),
  c(cor(all.DE.0.5$MC_ms, all.DE.0.5$SC_ms, method = 'pearson'),
    cor(all.DE.0.5$MC_ms, all.DE.0.5$EDC_ms, method = 'pearson'),
    cor(all.DE.0.5$SC_ms, all.DE.0.5$EDC_ms, method = 'pearson')),
  c(cor(all.DE.0.5$MC_my, all.DE.0.5$SC_my, method = 'pearson'),
    cor(all.DE.0.5$MC_my, all.DE.0.5$EDC_my, method = 'pearson'),
    cor(all.DE.0.5$SC_my, all.DE.0.5$EDC_my, method = 'pearson')),
  c(cor(all.DE.0.5$MC_sy, all.DE.0.5$SC_sy, method = 'pearson'),
    cor(all.DE.0.5$MC_sy, all.DE.0.5$EDC_sy, method = 'pearson'),
    cor(all.DE.0.5$SC_sy, all.DE.0.5$EDC_sy, method = 'pearson'))
)

pearson_results_05 %<>% setNames(c("cell_type_comparison", 
                                "mel_sim",
                                "mel_yak",
                                "sim_yak"))
pearson_results_05
pearson_results

## permutation test for these pearson correlations of logfold changes

# permute the logFC changes 1000 times
permutation <- lapply(1:10000, function(x){set.seed(x)
                                          setNames(data.frame(all.DE.1$gene,
                                                              sample(all.DE.1$MC_ms),
                                                              sample(all.DE.1$MC_my),
                                                              sample(all.DE.1$MC_sy),
                                                              sample(all.DE.1$SC_ms),
                                                              sample(all.DE.1$SC_my),
                                                              sample(all.DE.1$SC_sy),
                                                              sample(all.DE.1$EDC_ms),
                                                              sample(all.DE.1$EDC_my),
                                                              sample(all.DE.1$EDC_sy)),
                                                   c("gene",
                                                     "MC_ms",
                                                     "MC_my",
                                                     "MC_sy",
                                                     "SC_ms",
                                                     "SC_my",
                                                     "SC_sy",
                                                     "EDC_ms",
                                                     "EDC_my",
                                                     "EDC_sy"))})

# do pearson correlations on the permuted logFC changes on all combinations of 
# cell-types across branches
permutation_pearson <- rbindlist(lapply(1:10000, function(x){
                                          data.frame(
                                             cor(permutation[[x]]$MC_ms,
                                               permutation[[x]]$SC_ms, 
                                               method = "pearson"),
                                              cor(permutation[[x]]$MC_ms,
                                               permutation[[x]]$EDC_ms, 
                                               method = "pearson"),
                                              cor(permutation[[x]]$SC_ms,
                                               permutation[[x]]$EDC_ms, 
                                               method = "pearson"),
                                              cor(permutation[[x]]$MC_my,
                                               permutation[[x]]$SC_my, 
                                               method = "pearson"),
                                              cor(permutation[[x]]$MC_my,
                                               permutation[[x]]$EDC_my, 
                                               method = "pearson"),
                                              cor(permutation[[x]]$SC_my,
                                               permutation[[x]]$EDC_my, 
                                               method = "pearson"),
                                              cor(permutation[[x]]$MC_sy,
                                               permutation[[x]]$SC_sy, 
                                               method = "pearson"),
                                              cor(permutation[[x]]$MC_sy,
                                               permutation[[x]]$EDC_sy, 
                                               method = "pearson"),
                                              cor(permutation[[x]]$SC_sy,
                                               permutation[[x]]$EDC_sy, 
                                               method = "pearson") 
                                              )}))

permutation_pearson %<>% setNames(c("ms_MC_SC",
                                    "ms_MC_EDC",
                                    "ms_SC_EDC",
                                    "my_MC_SC",
                                    "my_MC_EDC",
                                    "my_SC_EDC",
                                    "sy_MC_SC",
                                    "sy_MC_EDC",
                                    "sy_SC_EDC"))

quants <- c(0,0.01,0.05,0.25,0.50,0.75,0.90,0.95,0.99,1)
apply(permutation_pearson, 2, quantile, probs = quants, na.rm = TRUE)


# plotting the data 
plot_ms_MC_SC  <- ggplot(data = all.DE.1, aes(x = MC_ms, y = SC_ms)) + 
                    geom_point(size = 0.4, alpha = 0.4) + 
                    labs(x = "logFC MC", y = "logFC SC") + 
                    lims(x = c(-6,6), y = c(-6,6)) + 
                    annotate(geom = "text", y = 5, x = -4.5, label = "r = 0.53", size = 2.8) +
                    theme_minimal(base_size = 9)

plot_ms_MC_SC 
plot_ms_MC_EDC <- ggplot(data = all.DE.1, aes(x = MC_ms, y = EDC_ms)) +
                    geom_point(size = 0.4, alpha = 0.4) + 
                    labs(x = "logFC MC", y = "logFC EDC") + 
                    lims(x = c(-6,6), y = c(-6,6))+ 
                    annotate(geom = "text", y = 5, x = -4.5, label = "r = 0.34", size = 2.8) +
                    theme_minimal(base_size = 9)

plot_ms_SC_EDC <- ggplot(data = all.DE.1, aes(x = SC_ms, y = EDC_ms)) +
                    geom_point(size = 0.4, alpha = 0.4) + 
                    labs(x = "logFC SC", y = "logFC EDC") + 
                    lims(x = c(-6,6), y = c(-6,6)) + 
                    annotate(geom = "text", y = 5, x = -4.5, label = "r = 0.38", size = 2.8) +
                    theme_minimal(base_size = 9)

plot_my_MC_SC  <- ggplot(data = all.DE.1, aes(x = MC_my, y = SC_my)) + 
                    geom_point(size = 0.4, alpha = 0.4) + 
                    labs(x = "logFC MC", y = "logFC SC") + 
                    lims(x = c(-6,6), y = c(-6,6)) + 
                    annotate(geom = "text", y = 5, x = -4.5, label = "r = 0.57", size = 2.8) +
                    theme_minimal(base_size = 9)

plot_my_MC_EDC <- ggplot(data = all.DE.1, aes(x = MC_my, y = EDC_my)) +
                    geom_point(size = 0.4, alpha = 0.4) + 
                    labs(x = "logFC MC", y = "logFC EDC") + 
                    lims(x = c(-6,6), y = c(-6,6)) + 
                    annotate(geom = "text", y = 5, x = -4.5, label = "r = 0.37", size = 2.8) +
                    theme_minimal(base_size = 9)

plot_my_SC_EDC <- ggplot(data = all.DE.1, aes(x = SC_my, y = EDC_my)) +
                    geom_point(size = 0.4, alpha = 0.4) + 
                    labs(x = "logFC SC", y = "logFC EDC") + 
                    lims(x = c(-6,6), y = c(-6,6)) + 
                    annotate(geom = "text", y = 5, x = -4.5, label = "r = 0.44", size = 2.8)  +
                    theme_minimal(base_size = 9)

plot_sy_MC_SC  <- ggplot(data = all.DE.1, aes(x = MC_sy, y = SC_sy)) + 
                    geom_point(size = 0.4, alpha = 0.4) + 
                    labs(x = "logFC MC", y = "logFC SC") + 
                    lims(x = c(-6,6), y = c(-6,6)) + 
                    annotate(geom = "text", y = 5, x = -4.5, label = "r = 0.53", size = 2.8)+
                    theme_minimal(base_size = 9)
                    
plot_sy_MC_EDC <- ggplot(data = all.DE.1, aes(x = MC_sy, y = EDC_sy)) +
                    geom_point(size = 0.4, alpha = 0.4) + 
                    labs(x = "logFC MC", y = "logFC EDC") + 
                    lims(x = c(-6,6), y = c(-6,6)) + 
                    annotate(geom = "text", y = 5, x = -4.5, label = "r = 0.28", size = 2.8) +
                    theme_minimal(base_size = 9)

plot_sy_SC_EDC <- ggplot(data = all.DE.1, aes(x = SC_sy, y = EDC_sy)) +
                    geom_point(size = 0.4, alpha = 0.4) + 
                    labs(x = "logFC SC", y = "logFC EDC") + 
                    lims(x = c(-6,6), y = c(-6,6)) + 
                    annotate(geom = "text", y = 5, x = -4.5, label = "r = 0.35", size = 2.8) +
                    theme_minimal(base_size = 9)

plot_grid(plot_ms_MC_SC,
          plot_my_MC_SC,
          plot_sy_MC_SC,
          plot_ms_MC_EDC,
          plot_my_MC_EDC,
          plot_sy_MC_EDC,
          plot_ms_SC_EDC,
          plot_my_SC_EDC,
          plot_sy_SC_EDC,
          labels = c(NULL))

## here is code to add gene labels to outliers if desired
library(ggrepel)
    geom_text_repel(
    data = subset(all.DE.1, abs(MC_ms) > 1.5 | abs(SC_ms) > 1.5),
    aes(label = gene),
    size = 5,
    box.padding = unit(0.35, "lines"),
    point.padding = unit(0.3, "lines"))

### here are the discordant DE examples (opposite direction logFC >1 in both cell types in a comparison)
subset(all.DE.1, MC_sy > 1  & SC_sy < -1 |
                 MC_sy < -1 & SC_sy > 1)
discordant_DE <- c("SP", "CG16713")
    
msy.so %>% VlnPlot(features = discordant_DE, split.by = "species")       

#### what about corellations of contrasts over the entire shared transcriptome? we would expect to see
#### a higher degree of corellation over cell-types in the mel-sim contrast, right?
    
all.DE <- join_all(list(main.DE, secondary.DE, ED.DE), by = "gene", type = "inner")[,c(1:4,9:11,16:18)]
all.DE %<>% setNames(c("gene", "MC_ms", "MC_my", "MC_sy", "SC_ms", "SC_my", "SC_sy", "EDC_ms", "EDC_my", "EDC_sy"))    
    
pearson_results_all <- data.frame(c("MC_SC", "MC_EDC", "SC_EDC"),
                                  c(cor(all.DE$MC_ms, all.DE$SC_ms, method = 'pearson'),
                                    cor(all.DE$MC_ms, all.DE$EDC_ms, method = 'pearson'),
                                    cor(all.DE$SC_ms, all.DE$EDC_ms, method = 'pearson')),
                                  c(cor(all.DE$MC_my, all.DE$SC_my, method = 'pearson'),
                                    cor(all.DE$MC_my, all.DE$EDC_my, method = 'pearson'),
                                    cor(all.DE$SC_my, all.DE$EDC_my, method = 'pearson')),
                                  c(cor(all.DE$MC_sy, all.DE$SC_sy, method = 'pearson'),
                                    cor(all.DE$MC_sy, all.DE$EDC_sy, method = 'pearson'),
                                    cor(all.DE$SC_sy, all.DE$EDC_sy, method = 'pearson')))

pearson_results_all %<>% setNames(c("cell_type_comparison", 
                                   "mel_sim",
                                   "mel_yak",
                                   "sim_yak"))
pearson_results_all

### do overall transcriptome (counts per gene) correlations among species

# get the average expression values for all genes by species
MC_mel <- rownames(subset(msy.so@meta.data, CellType == 'MC' & species == 'mel'))
MC_sim <- rownames(subset(msy.so@meta.data, CellType == 'MC' & species == 'sim'))
MC_yak <- rownames(subset(msy.so@meta.data, CellType == 'MC' & species == 'yak'))

SC_mel <- rownames(subset(msy.so@meta.data, CellType == 'SC' & species == 'mel'))
SC_sim <- rownames(subset(msy.so@meta.data, CellType == 'SC' & species == 'sim'))
SC_yak <- rownames(subset(msy.so@meta.data, CellType == 'SC' & species == 'yak'))

EDC_mel <- rownames(subset(msy.so@meta.data, CellType == 'EDC' & species == 'mel'))
EDC_sim <- rownames(subset(msy.so@meta.data, CellType == 'EDC' & species == 'sim'))
EDC_yak <- rownames(subset(msy.so@meta.data, CellType == 'EDC' & species == 'yak'))

main.expr.mel <- data.frame(main.expr[,colnames(main.expr) %in% MC_mel])
main.expr.sim <- data.frame(main.expr[,colnames(main.expr) %in% MC_sim])
main.expr.yak <- data.frame(main.expr[,colnames(main.expr) %in% MC_yak])

secondary.expr.mel <- data.frame(secondary.expr[,colnames(secondary.expr) %in% SC_mel])
secondary.expr.sim <- data.frame(secondary.expr[,colnames(secondary.expr) %in% SC_sim])
secondary.expr.yak <- data.frame(secondary.expr[,colnames(secondary.expr) %in% SC_yak])

ED.expr.mel <- data.frame(ED.expr[,colnames(ED.expr) %in% EDC_mel])
ED.expr.sim <- data.frame(ED.expr[,colnames(ED.expr) %in% EDC_sim])
ED.expr.yak <- data.frame(ED.expr[,colnames(ED.expr) %in% EDC_yak])

# calculate means
avg_MC_mel <- rowMeans(main.expr.mel)
avg_MC_sim <- rowMeans(main.expr.sim)
avg_MC_yak <- rowMeans(main.expr.yak)
avg_MC_all <- data.frame(avg_MC_mel, avg_MC_sim, avg_MC_yak)

avg_SC_mel <- rowMeans(secondary.expr.mel)
avg_SC_sim <- rowMeans(secondary.expr.sim)
avg_SC_yak <- rowMeans(secondary.expr.yak)
avg_SC_all <- data.frame(avg_SC_mel, avg_SC_sim, avg_SC_yak)

avg_EDC_mel <- rowMeans(ED.expr.mel)
avg_EDC_sim <- rowMeans(ED.expr.sim)
avg_EDC_yak <- rowMeans(ED.expr.yak)
avg_EDC_all <- data.frame(avg_EDC_mel, avg_EDC_sim, avg_EDC_yak)

## get pearson corellations for average expression values across species
pearson_avg_exp <- data.frame(c("MC", "SC", "EDC"),
                              c(round(cor(avg_MC_mel, avg_MC_sim), 2),
                                round(cor(avg_MC_mel, avg_MC_yak), 2),
                                round(cor(avg_MC_sim, avg_MC_yak), 2)),
                              c(round(cor(avg_SC_mel, avg_SC_sim), 2),
                                round(cor(avg_SC_mel, avg_SC_yak), 2),
                                round(cor(avg_SC_sim, avg_SC_yak), 2)),
                              c(round(cor(avg_EDC_mel, avg_EDC_sim), 2),
                                round(cor(avg_EDC_mel, avg_EDC_yak), 2),
                                round(cor(avg_EDC_sim, avg_EDC_yak), 2)))

pearson_avg_exp %<>% setNames(c("celltype", 
                                "mel_sim",
                                "mel_yak",
                                "sim_yak"))
pearson_avg_exp

### visualize corellations
plot_MC_melsim <- ggplot(avg_MC_all, aes(x = avg_MC_mel, y = avg_MC_sim)) + 
  geom_point(size = 0.4, alpha = 0.4) +
  labs(x = "mel avg expression", y = "sim avg expression") + 
  lims(x = c(0,7), y = c(0,7)) +
  theme_minimal(base_size = 9) +
  annotate(geom = "text", y = 5.75, x = 1.3, label = "r = 0.88", size = 2.8)

plot_MC_melyak <- ggplot(avg_MC_all, aes(x = avg_MC_mel, y = avg_MC_yak)) + 
  geom_point(size = 0.4, alpha = 0.4) +
  labs(x = "mel avg expression", y = "yak avg expression") + 
  lims(x = c(0,7), y = c(0,7)) +
  theme_minimal(base_size = 9) +
  annotate(geom = "text", y = 5.75, x = 1.3, label = "r = 0.86", size = 2.8)


plot_MC_simyak <- ggplot(avg_MC_all, aes(x = avg_MC_sim, y = avg_MC_yak)) + 
  geom_point(size = 0.4, alpha = 0.4) +
  labs(x = "sim avg expression", y = "yak avg expression") + 
  lims(x = c(0,7), y = c(0,7)) +
  theme_minimal(base_size = 9) +
  annotate(geom = "text", y = 5.75, x = 1.3, label = "r = 0.84", size = 2.8)


plot_SC_melsim <- ggplot(avg_SC_all, aes(x = avg_SC_mel, y = avg_SC_sim)) + 
  geom_point(size = 0.4, alpha = 0.4) +
  labs(x = "mel avg expression", y = "sim avg expression") + 
  lims(x = c(0,7), y = c(0,7)) +
  theme_minimal(base_size = 9) +
  annotate(geom = "text", y = 5.75, x = 1.3, label = "r = 0.81", size = 2.8)


plot_SC_melyak <- ggplot(avg_SC_all, aes(x = avg_SC_mel, y = avg_SC_yak)) + 
  geom_point(size = 0.4, alpha = 0.4) +
  labs(x = "mel avg expression", y = "yak avg expression") + 
  lims(x = c(0,7), y = c(0,7)) +
  theme_minimal(base_size = 9) +
  annotate(geom = "text", y = 5.75, x = 1.3, label = "r = 0.84", size = 2.8)


plot_SC_simyak <- ggplot(avg_SC_all, aes(x = avg_SC_sim, y = avg_SC_yak)) + 
  geom_point(size = 0.4, alpha = 0.4) +
  labs(x = "sim avg expression", y = "yak avg expression") + 
  lims(x = c(0,7), y = c(0,7)) +
  theme_minimal(base_size = 9) +
  annotate(geom = "text", y = 5.75, x = 1.3, label = "r = 0.74", size = 2.8)


plot_EDC_melsim <- ggplot(avg_EDC_all, aes(x = avg_EDC_mel, y = avg_EDC_sim)) + 
  geom_point(size = 0.4, alpha = 0.4) +
  labs(x = "mel avg expression", y = "sim avg expression") + 
  lims(x = c(0,7), y = c(0,7)) +
  theme_minimal(base_size = 9) +
  annotate(geom = "text", y = 5.75, x = 1.3, label = "r = 0.82", size = 2.8)


plot_EDC_melyak <- ggplot(avg_EDC_all, aes(x = avg_EDC_mel, y = avg_EDC_yak)) + 
  geom_point(size = 0.4, alpha = 0.4) +
  labs(x = "mel avg expression", y = "yak avg expression") + 
  lims(x = c(0,7), y = c(0,7)) +
  theme_minimal(base_size = 9) +
  annotate(geom = "text", y = 5.75, x = 1.3, label = "r = 0.80", size = 2.8)


plot_EDC_simyak <- ggplot(avg_EDC_all, aes(x = avg_EDC_sim, y = avg_EDC_yak)) + 
  geom_point(size = 0.4, alpha = 0.4) +
  labs(x = "sim avg expression", y = "yak avg expression") + 
  lims(x = c(0,7), y = c(0,7)) +
  theme_minimal(base_size = 9) +
  annotate(geom = "text", y = 5.75, x = 1.3, label = "r = 0.78", size = 2.8)


plot_grid(plot_MC_melsim,
          plot_MC_melyak,
          plot_MC_simyak,
          plot_SC_melsim,
          plot_SC_melyak,
          plot_SC_simyak,
          plot_EDC_melsim,
          plot_EDC_melyak,
          plot_EDC_simyak, labels = NULL)

## pearson corr for Sfps vs non-Sfps
 avg_MC_mel_sfp  <-  avg_MC_mel[sfps$V1]
 avg_MC_sim_sfp  <-  avg_MC_sim[sfps$V1]
 avg_MC_yak_sfp  <-  avg_MC_yak[sfps$V1]
 avg_SC_mel_sfp  <-  avg_SC_mel[sfps$V1]
 avg_SC_sim_sfp  <-  avg_SC_sim[sfps$V1]
 avg_SC_yak_sfp  <-  avg_SC_yak[sfps$V1]
avg_EDC_mel_sfp <-  avg_EDC_mel[sfps$V1]
avg_EDC_sim_sfp <-  avg_EDC_sim[sfps$V1]
avg_EDC_yak_sfp <-  avg_EDC_yak[sfps$V1]

pearson_avg_exp_sfp <- data.frame(c("MC", "SC", "EDC"),
                              c(round(cor(avg_MC_mel_sfp,  avg_MC_sim_sfp), 2),
                                round(cor(avg_MC_mel_sfp,  avg_MC_yak_sfp), 2),
                                round(cor(avg_MC_sim_sfp,  avg_MC_yak_sfp), 2)),
                              c(round(cor(avg_SC_mel_sfp,  avg_SC_sim_sfp), 2),
                                round(cor(avg_SC_mel_sfp,  avg_SC_yak_sfp), 2),
                                round(cor(avg_SC_sim_sfp,  avg_SC_yak_sfp), 2)),
                             c(round(cor(avg_EDC_mel_sfp, avg_EDC_sim_sfp), 2),
                               round(cor(avg_EDC_mel_sfp, avg_EDC_yak_sfp), 2),
                               round(cor(avg_EDC_sim_sfp, avg_EDC_yak_sfp), 2)))

pearson_avg_exp_sfp %<>% setNames(c("celltype", 
                                    "mel_sim",
                                    "mel_yak",
                                    "sim_yak"))
pearson_avg_exp_sfp
 
 avg_MC_mel_nonsfp  <-  avg_MC_mel[!names(avg_MC_mel)  %in% sfps$V1]
 avg_MC_sim_nonsfp  <-  avg_MC_sim[!names(avg_MC_sim)  %in% sfps$V1]
 avg_MC_yak_nonsfp  <-  avg_MC_yak[!names(avg_MC_yak)  %in% sfps$V1]
 avg_SC_mel_nonsfp  <-  avg_SC_mel[!names(avg_SC_mel)  %in% sfps$V1]
 avg_SC_sim_nonsfp  <-  avg_SC_sim[!names(avg_SC_sim)  %in% sfps$V1]
 avg_SC_yak_nonsfp  <-  avg_SC_yak[!names(avg_SC_yak)  %in% sfps$V1]
avg_EDC_mel_nonsfp <-  avg_EDC_mel[!names(avg_EDC_mel) %in% sfps$V1]
avg_EDC_sim_nonsfp <-  avg_EDC_sim[!names(avg_EDC_sim) %in% sfps$V1]
avg_EDC_yak_nonsfp <-  avg_EDC_yak[!names(avg_EDC_yak) %in% sfps$V1]

pearson_avg_exp_nonsfp <- data.frame(c("MC", "SC", "EDC"),
                                  c(round(cor(avg_MC_mel_nonsfp,  avg_MC_sim_nonsfp), 2),
                                    round(cor(avg_MC_mel_nonsfp,  avg_MC_yak_nonsfp), 2),
                                    round(cor(avg_MC_sim_nonsfp,  avg_MC_yak_nonsfp), 2)),
                                  c(round(cor(avg_SC_mel_nonsfp,  avg_SC_sim_nonsfp), 2),
                                    round(cor(avg_SC_mel_nonsfp,  avg_SC_yak_nonsfp), 2),
                                    round(cor(avg_SC_sim_nonsfp,  avg_SC_yak_nonsfp), 2)),
                                  c(round(cor(avg_EDC_mel_nonsfp, avg_EDC_sim_nonsfp), 2),
                                    round(cor(avg_EDC_mel_nonsfp, avg_EDC_yak_nonsfp), 2),
                                    round(cor(avg_EDC_sim_nonsfp, avg_EDC_yak_nonsfp), 2)))

pearson_avg_exp_nonsfp %<>% setNames(c("celltype", 
                                    "mel_sim",
                                    "mel_yak",
                                    "sim_yak"))
pearson_avg_exp_nonsfp
pearson_avg_exp_sfp

#######################################
## looking at top DE genes "by hand" ##
#######################################

#  it is useful to gather all the filtered DE genes together, 
# across contrasts within a cell type, so I can look at them manually

MC.DE.filtered <- subset(main.DE, gene %in% MC.DE.1)
SC.DE.filtered <- subset(secondary.DE, gene %in% SC.DE.1)
EDC.DE.filtered <- subset(ED.DE, gene %in% ED.DE.1)

### plot top 20 DE genes of each cell type
VlnPlot(msy.so, c(MC.DE.filtered[c(1:20),]$gene), split.by = "species",  cols = my_colors)
VlnPlot(msy.so, c(SC.DE.filtered[c(1:20),]$gene), split.by = "species",  cols = my_colors)
VlnPlot(msy.so, c(EDC.DE.filtered[c(1:20),]$gene), split.by = "species", cols = my_colors)

### examples for paper
DE_examples <- c("Acp95EF",
                 "Pkc53E",
                 "Obp58b",
                 "Marf1",
                 "msi",
                 "Nhe3",
                 "sv",
                 "Gld",
                 "Est-6",
                 "Spn28Dc",
                 "mbl",
                 "Oda")

DE_example_fig <- VlnPlot(msy.so, DE_examples, split.by = "species", pt.size = 0.1, ncol = 2, assay = "RNA", combine = F, cols = RColorBrewer::brewer.pal(3, "Set2"))

DE_example_fig %<>% lapply(FUN = function(x) x + theme(plot.title = element_text(size = 10, face = "plain"), 
                                                   axis.title.x = element_blank(),
                                                   axis.title.y = element_text(size = 9),
                                                   axis.text = element_text(size = 9),
                                                   legend.text = element_text(size = 9)))
DE_example_fig[c(2:length(DE_examples))] %<>% lapply(FUN = function(x) x + theme(legend.position = ""))
DE_example_fig[1] %<>% lapply(FUN = function(x) x + theme(legend.position = c(NA,4)))
DE_example_fig[c(2:4,6:8,10:12)] %<>% lapply(FUN = function(x) x + theme(axis.title.y = element_blank()))
DE_example_fig[c(1:8)] %<>% lapply(FUN = function(x) x + theme(axis.text.x = element_blank()))

plot_grid(plotlist = DE_example_fig, ncol = 4)
DE_example_fig %<>% lapply(FUN = function(x) x + theme(legend.position =  "none"))

#### wondering how often DE genes are non-marker genes of focal cell type?

## need to use markers called from all three species. That way, if there was a gain of expression
## specific to one species in one cell type, it also shows up as a marker. As is with the integrated dataset,
## markers are not called if expressed in only one of three species

markerlist_allspp_MC  <- unique(c(subset(markers_mel_ortho, cluster == "MC")$gene,
                                  subset(markers_sim_ortho, cluster == "MC")$gene,
                                  subset(markers_yak_ortho, cluster == "MC")$gene))
markerlist_allspp_SC  <- unique(c(subset(markers_mel_ortho, cluster == "SC")$gene,
                                  subset(markers_sim_ortho, cluster == "SC")$gene,
                                  subset(markers_yak_ortho, cluster == "SC")$gene))
markerlist_allspp_EDC <- unique(c(subset(markers_mel_ortho, cluster == "EDC")$gene,
                                  subset(markers_sim_ortho, cluster == "EDC")$gene,
                                  subset(markers_yak_ortho, cluster == "EDC")$gene))

# make the markers singlets so we can include things that are markers of two cell types
markerlist_allspp_MC_singlet <- markerlist_allspp_MC[!markerlist_allspp_MC %in% c(markerlist_allspp_SC, markerlist_allspp_EDC)]
markerlist_allspp_SC_singlet <- markerlist_allspp_SC[!markerlist_allspp_SC %in% c(markerlist_allspp_MC, markerlist_allspp_EDC)]
markerlist_allspp_EDC_singlet <- markerlist_allspp_EDC[!markerlist_allspp_EDC %in% c(markerlist_allspp_MC, markerlist_allspp_SC)]

MC.DE.nonmarkers <- subset(MC.DE.filtered, !gene %in% markerlist_allspp_MC_singlet)
36 / 132
VlnPlot(msy.so, c(MC.DE.nonmarkers[c(1:20),]$gene), split.by = "species",  cols = my_colors)

SC.DE.nonmarkers <- subset(SC.DE.filtered, !gene %in% markerlist_allspp_SC_singlet)
90 / 111
VlnPlot(msy.so, c(SC.DE.nonmarkers[c(1:20),]$gene), split.by = "species",  cols = my_colors)

EDC.DE.nonmarkers <- subset(EDC.DE.filtered, !gene %in% markerlist_allspp_EDC_singlet)
56/224
VlnPlot(msy.so, c(SC.DE.nonmarkers[c(1:20),]$gene), split.by = "species",  cols = my_colors)

(36+90+56)/(132+111+224)

# ok finally. What about percent of marker genes that show DE?
table(markerlist_allspp_MC %in% MC.DE.1)
table(markerlist_allspp_SC %in% SC.DE.1)
table(markerlist_allspp_EDC %in% ED.DE.1)

table(subset(markers_merged_significant, cluster == "MC")$gene %in% MC.DE.1)
73/(73+236)
table(subset(markers_merged_significant, cluster == "SC")$gene %in% SC.DE.1)
25/(25+96)
table(subset(markers_merged_significant, cluster == "EDC")$gene %in% ED.DE.1)
123/(123+255)

markers_DE <- data.frame(c(309,121,378), c(73,25,12), row.names = c("MC", "SC", "EDC"))
fisher.multcomp(as.matrix(markers_DE))

####### tables for the paper #######

### MC DE table ###
# join to average expression - the ppl want it

avg.msy <- AverageExpression(msy.so, return.seurat = T, add.ident = "species")[["RNA"]]
avg.msy <- data.frame(GetAssayData(avg.msy, slot = "data"))
avg.msy %<>% tibble::rownames_to_column("gene")
head(avg.msy)

MC.DE.filtered.avgs <- inner_join(MC.DE.filtered, avg.msy[,c(1,2,6,9)], by = "gene")[,c(1:4,6,8:11)]
write.table(MC.DE.filtered.avgs, "~/Documents/Begun/STAR_counts/MC_DE.tsv", quote = F, sep = "\t", row.names = F)

SC.DE.filtered.avgs <- inner_join(SC.DE.filtered, avg.msy[,c(1,3,7,10)], by = "gene")[,c(1:4,6,8:11)]
write.table(SC.DE.filtered.avgs, "~/Documents/Begun/STAR_counts/SC_DE.tsv", quote = F, sep = "\t", row.names = F)

EDC.DE.filtered.avgs <- inner_join(EDC.DE.filtered, avg.msy[,c(1,4,5,8)], by = "gene")[,c(1:4,6,8:11)]
write.table(EDC.DE.filtered.avgs, "~/Documents/Begun/STAR_counts/EDC_DE.tsv", quote = F, sep = "\t", row.names = F)

all.DE.filtered.avgs <- inner_join(all.DE.1, avg.msy, by = "gene")[,-c(5,9,13)]
```
