## Supplementary material for "Single-nucleus transcriptomes reveal functional and evolutionary properties of cell types in the Drosophila accessory gland": Dataset S10: melanogaster_seurat.R

```
library(Seurat)
library(cowplot)
library(dplyr)
library(plotly)
library(ggplot2)
library(data.table)
library(magrittr)
library(RVAideMemoire)
library(tibble)
library(ggtext)

my_colors <- c("slategray3", "mediumseagreen","darkslateblue")
my_colors <- c("slategray3", "#349A83","#5346A6")
# import the counts
counts <- read.table("mel.cleaned.tsv", stringsAsFactors = F)
counts <- counts[,order(colnames(counts))]
counts[1:5,1:5]

# swap FBgn for gene names
synonyms <- fread("fb_synonym_fb_2020_02.tsv") # this is the synonym table from flybased
rownames(counts) <- synonyms$current_symbol[match(rownames(counts), synonyms$`##primary_FBid`)]
 
# import unannotated counts
counts_unannotated <- read.table("unannotated_v2_cleaned.tsv")

# catalog the IDs for later analyses of these genes
unannotated_geneIDs <- gsub(rownames(counts_unannotated),pattern = "_", replacement = "-")

# merge all genes together
counts_all <- rbind(counts, counts_unannotated)
counts_all[16850:16864,1:3]

# remove mitochondrial genome genes (background expression)
counts_all <- counts_all[-grep("mt:", rownames(counts_all)),]
# remove CR332222, pseudogene overlaping with RpS6 (MC subpop 2 marker)
counts_all <- counts_all[-grep("CR33222", rownames(counts_all)),]

# create the seurat object
mel.so <- CreateSeuratObject(counts_all, min.cells = 3, min.features = 100)

# overview of the data
VlnPlot(mel.so, features = c("nFeature_RNA", "nCount_RNA"), cols = "slateblue")
FeatureScatter(mel.so, feature1 = "nCount_RNA", feature2 = "nFeature_RNA")

# pre-processing / transformation of the data
mel.so %<>% NormalizeData(normalization.method = "LogNormalize", scale.factor = 10000)

mel.so %<>% FindVariableFeatures(selection.method = "dispersion")
top60 <- head(VariableFeatures(mel.so), 60)
top60
plot1 <- VariableFeaturePlot(mel.so)
plot2 <- LabelPoints(plot = plot1, points = top60, repel = T, xnudge = 0, ynudge = 0)
plot2

# which of the de novo and unannotated genes are variable features?
unannotated_geneIDs_variable <- VariableFeatures(mel.so)[
                                        VariableFeatures(mel.so) %in% unannotated_geneIDs]
unannotated_geneIDs_variable
# just 3 of these count as variable features, but keep in mind that they are lowly expressed, and
# there could be others that show a clear pattern without being a top 2000 variable feature. 
# so manual investigation is critical

# Scale data to have equal variance. here scaling every gene not just var features, so we 
# can analyze cell-type bias of any gene we want, and plot any gene.
mel.so %<>% ScaleData(features = rownames(mel.so@assays$RNA@counts))

# PCA
mel.so %<>% RunPCA()
DimPlot(object = mel.so, dims = c(1,2), reduction = "pca")
DimPlot(object = mel.so, dims = c(1,3), reduction = "pca")
ElbowPlot(mel.so)
# elbow plot shows significance of first 4 to 6 PCs

# JackStraw permutation analysis of PC significance
mel.so <- JackStraw(
  object = mel.so, dims = 40)
mel.so <- ScoreJackStraw(mel.so, dims = 1:40)

JackStrawPlot(object = mel.so, dims = 1:40)
# observe plot to decide what PCs to use
use.pcs = c(1:7,11,16)

#UMAP
mel.so <- RunUMAP(mel.so, reduction = "pca", dims = use.pcs)
mel.so <- FindNeighbors(mel.so, reduction = "pca", dims = use.pcs)

# look at cluster nums at a series of resolutions
mel.so <- FindClusters(
  object = mel.so,
  resolution = seq(0.2,1.4,0.1),
  verbose = FALSE)
sapply(grep("res",colnames(mel.so@meta.data),value = TRUE),
       function(x) length(unique(mel.so@meta.data[,x])))
# choose resolution - simple dataset, avoid over-splitting clusters
Idents(mel.so) <- "RNA_snn_res.0.2"

# UMAP Visualization
DimPlot(mel.so, reduction = "umap", label = T, pt.size = 2) +
  theme_minimal(base_size = 15)

FeaturePlot(mel.so, features = c('nCount_RNA'), pt.size=2)
FeaturePlot(mel.so, features = c('nFeature_RNA'), pt.size=2)

## assign cell types
mel.so %<>% RenameIdents('0' = 'MCsp1',
                         '1' = 'MCsp2',
                         '3' = 'SC', 
                         '2' = 'EDC')

subpops.umap <- DimPlot(mel.so, reduction = "umap", label = F, pt.size = 0.3, cols = c(my_colors[1], "salmon", my_colors[2:3])) + theme(plot.title = element_text(size = 10), 
                                                                                                       axis.title.x = element_text(size = 9),
                                                                                                       axis.title.y = element_text(size = 9),
                                                                                                        legend.text = element_text(size = 9),
                                                                                                       axis.text.x = element_text(size = 9),
                                                                                                       axis.text.y = element_text(size = 9))
subpops.umap[[1]]$layers[[1]]$aes_params$alpha = 0.8
subpops.umap

# marker genes 
markers_all <- FindAllMarkers(
  object = mel.so, 
  only.pos = TRUE, 
  min.pct = 0.25, 
  thresh.use = 0.25
)

dim(markers_all)
head(markers_all)

FeaturePlot(mel.so, features = c('Acp76A'), pt.size=2)
FeaturePlot(mel.so, features = c('abd-A'), pt.size=2)

head(subset(markers_all, cluster == 2))

markers_all_sig <- subset(markers_all, p_val_adj < 0.05)

# find markers defining the subpops
markers_MC_subpops <- FindMarkers(mel.so, ident.1 = 'MCsp2',
                                          ident.2 = 'MCsp1')
markers_MC_subpops  %<>% rownames_to_column("gene")
markers_MC_subpops_sig <- subset(markers_MC_subpops, p_val_adj < 0.05)

# most significantly marker gene of cluster 1 is Gmap 
mel.so %>% VlnPlot("Gmap")
mel.so %>% VlnPlot("eEF2")
mel.so %>% VlnPlot("lncRNA:roX1")

# it appeasrs that SFPs are more highly expressed in the small subpoplution of Main cells
sfp.ridge <- VlnPlot(mel.so, c("Mst57Db", 
                                 "Sfp96F", 
                                 "SP",
                                 "Acp36DE"), 
                       cols = c(my_colors[1], "salmon", my_colors[2:3]), combine = F, pt.size = 0.1)

sfp.ridge <- lapply(X = sfp.ridge, FUN = function(x) x + theme(legend.position = "none", 
                                                               plot.title =   element_text(size = 11.5, face = "plain"),
                                                               axis.title.y = element_text(size = 10),
                                                               axis.text.y  = element_text(size = 10),
                                                               axis.title.x = element_blank(),
                                                               axis.text.x  = element_text(size = 10)))
sfp.ridge[c(2,4)] %<>% lapply(FUN = function(x) x + theme(axis.title.y = element_blank()))
sfp.ridge[c(1,2)] %<>% lapply(FUN = function(x) x + theme(axis.text.x = element_blank()))
plot_grid(plotlist = sfp.ridge, ncol = 2, rel_heights = c(1,1.15))

## some nice non-SFPs
nonsfp.ridge <- VlnPlot(mel.so ,c("eEF2", 
                                    "RpS25",
                                    "RpL31", 
                                    "Gmap",
                                    "lncRNA:roX1",
                                    "lncRNA:Hsromega"), 
                       cols = c(my_colors[1], "salmon", my_colors[2:3]), combine = F, pt.size = 0.1)  

nonsfp.ridge <- lapply(X = nonsfp.ridge, FUN = function(x) x + theme(legend.position = "none", 
                                                               plot.title =   element_text(size = 13, face = "plain"),
                                                               axis.title.x = element_text(size = 10.4),
                                                               axis.text.x  = element_text(size = 10.4),
                                                               axis.title.y = element_blank(),
                                                               axis.text.y  = element_text(size = 10.4)))

plot_grid(plotlist = nonsfp.ridge, ncol = 3, labels = c("A", "", "", "B", "", ""))

# subset the non-sfps so we can get a gene list for DAVID GO analysis
sfps <- read.table("alex_all_IDs.txt")

markers_MC_subpops_sig_nonSFP <- subset(markers_MC_subpops_sig, !gene %in% sfps$V1)
markers_MC_subpops_sig_SFP <- subset(markers_MC_subpops_sig, gene %in% sfps$V1)

top_10 <- markers_MC_subpops_sig_nonSFP %>% slice_max(order_by = avg_logFC, n =10)
bot_10 <- markers_MC_subpops_sig_nonSFP %>% slice_min(order_by = avg_logFC, n =10)

write.table(top_20$gene, "top20_MCsubcluster", col.names = F, row.names = F, quote = F)
write.table(bot_20$gene, "bot20_MCsubcluster", col.names = F, row.names = F, quote = F)

mel.so %>% VlnPlot('nCount_RNA', cols = c(my_colors[1], "salmon", my_colors[2:3]), pt.size = 0.3) + theme(text = element_text(size = 9),
                                                                                           axis.text = element_text(size = 9),
                                                                                           axis.title.x = element_blank(),
                                                                                           legend.position = "none") +
                                                                                      ylab("counts")
mel.so %>% VlnPlot('nFeature_RNA', cols = c(my_colors[1], "salmon", my_colors[2:3]))
mel.so %>% RidgePlot(c(top_20$gene[1:10], bot_20$gene[1:10]), cols = c(my_colors[1], "salmon", my_colors[2:3]))

heatmap <- DoHeatmap(mel.so, features = c(top_10$gene, bot_10$gene), group.colors = c(my_colors[1], "salmon", my_colors[2:3]), size = 2.5) + 
            theme(text = element_text(size = 9),
                  axis.text.y = element_text(color = "black"))
ggplot2::ggsave(filename = "~/Documents/Begun/AG_nuclei_pub_figs_tables/transcriptome_heterogeneity_among_main_cell_subpopulations/subpop_heatmap.tiff",
                plot = heatmap, height = 3.3, width = 6.95, dpi = "retina")

# look at SFP and nonSFP logFC distributions
markers_MC_subpops_sig_nonSFP$type <- 'non-SFP'
markers_MC_subpops_sig_SFP$type <- 'SFP'
markers_MC_subpops_sig_labelled <- rbind(markers_MC_subpops_sig_SFP, markers_MC_subpops_sig_nonSFP)

## export the labelled subpopulation data for paper
write.table(x = markers_MC_subpops_sig_labelled, file = "~/Documents/Begun/STAR_counts/mel_labelled_subpopMc1Mc2_markers_5_09_21.tsv", quote = F, row.names = F, sep = "\t")

ggplot(markers_MC_subpops_sig_labelled, aes(x = avg_logFC))  +
  geom_density(aes(fill = type), alpha = 0.6, size = 0.3) +
  theme_minimal(base_size = 8) +
  xlab(expression(paste("average logFC ", frac(MCsp2, MCsp1))))

MCsp1 <- names(Idents(mel.so)[Idents(mel.so) == "MCsp1"])
MCsp2 <- names(Idents(mel.so)[Idents(mel.so) == "MCsp2"])
SC <- names(Idents(mel.so)[Idents(mel.so) == "SC"])
EDC <- names(Idents(mel.so)[Idents(mel.so) == "EDC"])

MCsp1_meta <- subset(mel.so@meta.data, rownames(mel.so@meta.data) %in% MCsp1)
MCsp2_meta <- subset(mel.so@meta.data, rownames(mel.so@meta.data) %in% MCsp2)
SC_meta <- subset(mel.so@meta.data, rownames(mel.so@meta.data) %in% SC)
EDC_meta <- subset(mel.so@meta.data, rownames(mel.so@meta.data) %in% EDC)

plot(density(MCsp1_meta$nCount_RNA))
plot(density(MCsp2_meta$nCount_RNA))
plot(density(SC_meta$nCount_RNA))
plot(density(EDC_meta$nCount_RNA))

median(MCsp1_meta$nCount_RNA)
median(MCsp2_meta$nCount_RNA)
median(SC_meta$nCount_RNA)
median(EDC_meta$nCount_RNA)

# these are not normal so use non parametric test
kruskal.test(list(MCsp1_meta$nCount_RNA, MCsp2_meta$nCount_RNA, SC_meta$nCount_RNA, EDC_meta$nCount_RNA))
pairwise.wilcox.test(list(MCsp1_meta$nCount_RNA, MCsp2_meta$nCount_RNA, SC_meta$nCount_RNA, EDC_meta$nCount_RNA))

ncounts.df <- rbind(setNames(data.frame(MCsp1_meta$nCount_RNA, "MCsp1"), c("counts", "celltype")),
                    setNames(data.frame(MCsp2_meta$nCount_RNA, "MCsp2"), c("counts", "celltype")),
                    setNames(data.frame(SC_meta$nCount_RNA, "SC"), c("counts", "celltype")),
                    setNames(data.frame(EDC_meta$nCount_RNA, "EDC"), c("counts", "celltype")))

aggregate(counts ~ celltype, ncounts.df, median)
kruskal.test(counts ~ celltype, ncounts.df)
pairwise.wilcox.test(ncounts.df$counts, ncounts.df$celltype)

###########################################
# merge the types for downstream analysis #
###########################################

mel.so <- RenameIdents(object = mel.so, 'MCsp1' = 'MCsp2')
mel.so <- RenameIdents(object = mel.so, 'MCsp2' = 'MC')

table(Idents(mel.so))

mel.umap <- DimPlot(mel.so, reduction = "umap", label = F, pt.size = 0.3, cols = c(my_colors)) +   theme(plot.title = element_text(size = 9), 
                                                                                                       axis.title.x = element_text(size = 9),
                                                                                                       axis.title.y = element_text(size = 9),
                                                                                                        legend.text = element_text(size = 9),                                                                                                      axis.text = element_text(size = 9))
mel.umap[[1]]$layers[[1]]$aes_params$alpha = 0.7

# characteristic markers
char.markers <- c("SP", "Acp36DE", "Acp95EF",
                  "lectin-46Ca", "abd-A", "lncRNA:iab8",
                  "vvl", "Dup99B", "Abd-B")
marker.umap <- FeaturePlot(mel.so, char.markers, cols = c("lightgrey", "darkcyan"), pt.size = 0.1, combine = F)
marker.umap <- lapply(X = marker.umap, FUN = function(x) x + theme(plot.title = element_text(size = 9, face = "plain"), 
                                                                 axis.title.x = element_text(size = 9),
                                                                 axis.title.y = element_text(size = 9)))
marker.umap[c(2,3,5,6,8,9)] %<>% lapply(FUN = function(x) x + theme(axis.title.y = element_blank()))
marker.umap[c(1:6)] %<>% lapply(FUN = function(x) x + theme(axis.title.x = element_blank()))
plot_grid(plotlist = c(marker.umap), ncol = 3)

## a useful thing to know is the total expressed genes per cell type
MC.expr  <- as.matrix(GetAssayData(subset(msy.so, idents = 'MC')))
SC.expr  <- as.matrix(GetAssayData(subset(msy.so, idents = 'SC')))
EDC.expr <- as.matrix(GetAssayData(subset(msy.so, idents = 'EDC')))

length(which(rowSums(MC.expr) > 3))
length(which(rowSums(SC.expr) > 3))
length(which(rowSums(EDC.expr) > 3))

# merged markers
markers_merged <- FindAllMarkers(
  object = mel.so, 
  only.pos = TRUE, 
  min.pct = 0.25, 
  thresh.use = 0.25
)

# subset significant bonferoni
markers_merged_sig <- subset(markers_merged, markers_merged$p_val_adj < 0.05)

# get the markers characterizing MC + SC vs EDC
markers_AG <- FindMarkers(mel.so, ident.1 = c("MC", "SC"),
                                  ident.2 = "EDC")
markers_AG_sig <- subset(markers_AG, p_val_adj < 0.05)

#### heatmap for top marker genes 
top20 <- markers_merged_sig %>% group_by(cluster) %>% top_n(20, avg_logFC)
dim(top20)

# create a label list to highlight SFPs
genes.to.label <- as.vector(sfps$V1)
labels <- rep(x = "black", times = length(x = top20$gene))
labels[match(x = genes.to.label, table = top20$gene)] <- "slateblue"

heatmap <- DoHeatmap(object = mel.so, 
  features = top20$gene, group.colors = my_colors, size = 2) + 
    theme(axis.text.y = element_text(color = rev(x = labels)),
          text = element_text(size = 6))
heatmap
ggplot2::ggsave(filename = "~/Documents/Begun/AG_nuclei_pub_figs_tables/characterization_of_cell_type_tanscriptomes_in_the_drosophila_melanogaster_accessory_gland/mel_heatmap_nolabels.png",
                plot = heatmap, height = 3.95, width = 6.43)

### how do SFPs break down across the markers?

# join SFPs to the markers
mel_markers_SFPs <- left_join(markers_merged_sig, sfps, by = c("gene" = "V1"))

summary(!is.na(mel_markers_SFPs$V2)) # TRUE = SFP
summary(!is.na(subset(mel_markers_SFPs, cluster == "MC")$V2))
summary(!is.na(subset(mel_markers_SFPs, cluster == "SC")$V2))
summary(!is.na(subset(mel_markers_SFPs, cluster == "EDC")$V2))

SFPmarkers_summary <- setNames(data.frame(c(124, 72, 241),
                                          c(92, 10, 21)), 
                               c("nonSFP", "SFP"))

rownames(SFPmarkers_summary) <- c("MC", "SC", "EDC")
SFPmarkers_summary

G.test(as.matrix(SFPmarkers_summary))
pairwise.G.test(as.matrix(SFPmarkers_summary))

## get the actual SFPs by the cell type
SFP_markers_only <- inner_join(markers_merged_sig, sfps, by = c("gene" = "V1"))
MC_SFPs <- subset(SFP_markers_only, cluster == "MC")$gene
SC_SFPs <- subset(SFP_markers_only, cluster == "SC")$gene
EDC_SFPs <- subset(SFP_markers_only, cluster == "EDC")$gene

# SC + EDC marker table
SFP_markers_SC_EDC <- subset(SFP_markers_only, cluster == "SC" | cluster == "EDC")

# now what about bias over cell-types? Not just marker genes, but let's look at the spread
# of expression bias over all SFPs

bias_exp <- FindAllMarkers(
  object = mel.so, 
  only.pos = FALSE, 
  min.pct = 0, 
  logfc.threshold = 0,
  return.thresh = 1
)

bias_exp$gene[bias_exp$gene %in% sfps$V1]

# here are SFPs apparently not expressed in the dataset:
sfps_notexpressed <- as.list(sfps$V1[!sfps$V1 %in% bias_exp$gene])
sfps_expressed <- unique(as.list(sfps$V1[sfps$V1 %in% bias_exp$gene]))

# SFP bias
sfp_bias_exp <- bias_exp %>% subset(gene %in% sfps$V1)
sfp_bias_exp_pos <- sfp_bias_exp %>% subset(avg_logFC > 0)

# SP network Sfps
spnetwork <- c("intr", "lectin-46Ca", "lectin-46Cb", "aqrs", "antr", "CG9997", "CG17575", "Sems", "frma", "hdly", "Esp", "SP", "SPR")
bias_exp %>% subset(gene %in% spnetwork)

# where are the positive values?
summary(sfp_bias_exp_pos$cluster)

# write table outputs
write.table(sfp_bias_exp_pos,  file = "~/Documents/Begun/STAR_counts/mel_sfp_bias_5_06_21.tsv", quote = F, row.names = F, sep = "\t")

# here are the genes that have a + value in more than one cell type
sfp_pos_dup <- sfp_bias_exp_pos$gene[duplicated(sfp_bias_exp_pos$gene)]
View(sfp_bias_exp_pos %>% subset(gene %in% sfp_pos_dup)) #& p_val_adj < 0.05))

#####################################
# fetch the acerage expression data #
#####################################
cluster.averages <- AverageExpression(mel.so)[["RNA"]]
cluster.averages %<>% rownames_to_column("gene")
head(cluster.averages)

sfp.cluster.averages <- cluster.averages %>% subset(gene %in% sfps$V1)
head(sfp.cluster.averages)

# mark marker genes in the expression table
sfp.cluster.averages %<>% mutate(marker = case_when(gene %in% MC_SFPs ~ "MC",
                                                   gene %in% SC_SFPs ~ "SC",
                                                   gene %in% EDC_SFPs ~ "EDC",
                                                   !gene %in% c(MC_SFPs, SC_SFPs, EDC_SFPs) ~ "N/A"))

# plot avg exp in different cells against one another
pMCSC <- ggplot(sfp.cluster.averages, aes(x = log(MC+1), y = log(SC+1))) + 
            geom_point(aes(color = marker), alpha = 0.8, size = 1.5) + 
            theme_minimal(base_size = 9) + 
            labs(x = "log(avg expression MC)", y = "log(avg expression SC)") +
            scale_color_manual(values = c(my_colors, "honeydew4"),
                               breaks=c("MC","SC","EDC", "N/A")) + 
            lims(x = c(0,8), y = c(0,8)) +
            geom_abline(slope = 1, intercept = c(0,0), alpha = 0.3)
          
pMCEDC <- ggplot(sfp.cluster.averages, aes(x = log(MC+1), y = log(EDC+1))) + 
            geom_point(aes(color = marker), alpha = 0.8, size = 1.5) + 
            theme_minimal(base_size = 9) + 
            labs(x = "log(avg expression MC)", y = "log(avg expression EDC)") +
            scale_color_manual(values = c(my_colors, "honeydew4"),
                               breaks=c("MC","SC","EDC", "N/A"))+
            lims(x = c(0,8), y = c(0,8)) +
            geom_abline(slope = 1, intercept = c(0,0), alpha = 0.3) 
          
pSCEDC <- ggplot(sfp.cluster.averages, aes(x = log(EDC+1), y = log(SC+1))) + 
            geom_point(aes(color = marker), alpha = 0.8, size = 1.5) + 
            theme_minimal(base_size = 9) + 
            labs(x = "log(avg expression EDC)", y = "log(avg expression SC)") +
            scale_color_manual(values = c(my_colors, "honeydew4"),
                               breaks=c("MC","SC","EDC", "N/A"))+
            lims(x = c(0,8), y = c(0,8)) +
            geom_abline(slope = 1, intercept = c(0,0), alpha = 0.3)

plot_grid(pMCSC, pMCEDC, pSCEDC, nrow = 3, labels = c("A", "B", "C"), label_x = -0.0075)

# get some more stats about expression
summary(lm(SC ~ MC, subset(sfp.cluster.averages, marker == "N/A")))
summary(lm(EDC ~ MC, subset(sfp.cluster.averages, marker == "N/A")))
summary(lm(SC ~ EDC, subset(sfp.cluster.averages, marker == "N/A")))

sfp.na.averages <- sfp.cluster.averages %>% subset(marker == "N/A")
dim(subset(sfp.na.averages, max.col(sfp.na.averages[,2:4]) == 1))
dim(subset(sfp.na.averages, max.col(sfp.na.averages[,2:4]) == 2))
dim(subset(sfp.na.averages, max.col(sfp.na.averages[,2:4]) == 3))

dim(subset(sfp.cluster.averages, max.col(sfp.cluster.averages[,2:4]) == 1))
dim(subset(sfp.cluster.averages, max.col(sfp.cluster.averages[,2:4]) == 2))
dim(subset(sfp.cluster.averages, max.col(sfp.cluster.averages[,2:4]) == 3))

## do metric MDS on SFP expression
# get our scaled SFP data 
scaled.sfp <- as.data.frame(t(mel.so@assays$RNA@scale.data[rownames(mel.so@assays$RNA@scale.data) %in% sfps$V1,]))
dist.sfp <- dist(scaled.sfp,method = "euclidian")
mat.sfp <- as.matrix(dist.sfp)
mds.sfp <- cmdscale(mat.sfp, eig = TRUE, k = 2)  # Perform the actual MDS

mds.sfp.df <- data.frame(xcor = mds.sfp$points[,1],
                         ycor = mds.sfp$points[,2],
                         celltype = Idents(mel.so))

ggplot(mds.sfp.df, aes(x = xcor, y = ycor)) + 
  geom_point(aes(color = celltype)) + 
  theme_minimal()

## logFC distributions by cell type
plogfc <- ggplot(sfp_bias_exp, aes(x = avg_logFC, fill = cluster)) + 
            geom_density(alpha = 0.7) +
            scale_fill_manual(values = my_colors)+
            theme_minimal(base_size = 9) +
            xlab(element_text("avg log(FC)"))
kruskal.test(avg_logFC ~ cluster, sfp_bias_exp)
pairwise.wilcox.test(sfp_bias_exp$avg_logFC, sfp_bias_exp$cluster, p.adjust.method = "BH")
aggregate(avg_logFC ~ cluster, sfp_bias_exp, median)

plot_grid(pMCSC, pMCEDC, pSCEDC, plogfc, nrow = 2, labels = c("A", "B", "C", "D"), label_x = -0.0075)

## now visualizing the SFPs specific to SC and EDC
FeaturePlot(mel.so, as.vector(subset(SFP_markers_only, cluster == "EDC")$V3), cols = c("lightgrey", "darkcyan"))
FeaturePlot(mel.so, as.vector(subset(SFP_markers_only, cluster == "SC")$V3), cols = c("lightgrey", "darkcyan"))

## and get the markers that are NOT SFPs
nonSFP_markers <- anti_join(markers_merged_sig, sfps, by = c("gene" = "V1"))

## plots of some interesting marker genes
FeaturePlot(mel.so, cols = c("lightgrey", "darkcyan"), pt.size = 1.5, ncol = 3,
            features = c("SP", "Acp32CD", "Dup99B", "Acp26Aa", "lectin-46Ca", "Est-6"))

FeaturePlot(mel.so, cols = c("lightgrey", "darkcyan"), pt.size = 1.5, ncol = 3,
            features = c("lncRNA:iab8", "abd-A", "dve"))

FeaturePlot(mel.so, cols = c("lightgrey", "darkcyan"), pt.size = 1.5, ncol = 3,
            features = c("sas", "msi", "form3"))

FeaturePlot(mel.so, cols = c("lightgrey", "darkcyan"), pt.size = 1.5, ncol = 3,
            features = c("vvl", "Abd-B", "Anp"))

FeaturePlot(mel.so, cols = c("lightgrey", "darkcyan"), pt.size = 1.5, ncol = 3,
            features = c("Sfp93F", "Ae2", "axed"))

### for the paper 

markers_paper_fig <- c("SP",
                       "Acp36DE",
                       "lncRNA:iab8",
                       "lectin-46Ca",
                       "Dup99B",
                       "vvl",
                       "Obp22a",
                       "lncRNA:TS14",
                       "CG13965",
                       "sas",
                       "Sfp93F",
                       "Ae2")

eg.marker.fig <- FeaturePlot(mel.so, cols = c("lightgrey", "darkcyan"), pt.size = 0.1,
                    features = markers_paper_fig, combine = F)

eg.marker.fig %<>% lapply(FUN = function(x) x + theme(plot.title = element_text(size = 9, face = "plain"), 
                                                                   axis.title.x = element_text(size = 9),
                                                                   axis.title.y = element_text(size = 9),
                                                                    axis.text.y = element_text(size = 9),
                                                                    axis.text.x = element_text(size = 9),
                                                                    legend.position = "none"))
eg.marker.fig[c(2:6,8:12)] %<>% lapply(FUN = function(x) x + theme(axis.title.y = element_blank()))
eg.marker.fig[c(1:6)] %<>% lapply(FUN = function(x) x + theme(axis.title.x = element_blank()))
plot_grid(plotlist = eg.marker.fig, labels = c("D", "", "", "", "", "",
                                               "E", "", "", "", "", ""), ncol = 6)

###############################################
## evolutionary  analysis of marker proteins ##
###############################################

### marker genes

main_markers <- subset(markers_merged_sig, cluster == "MC")
secondary_markers <- subset(markers_merged_sig, cluster == "SC")
ED_markers <- subset(markers_merged_sig, cluster == "EDC")

# MK test #

# import MK and calc alpha, malawi unpolarized. MK results from Fraisse, Sala, and Vicoso 2019
# https://academic.oup.com/mbe/article/36/3/500/5261346#131592134
library(data.table)
mk <- fread("MK.tsv")
colnames(mk)

# joining marker genes to alpha values
markers_merged <- inner_join(markers_merged, synonyms, by = c("gene" = "current_symbol"))[,1:8]
markers_alpha <- inner_join(markers_merged, mk, by = c("##primary_FBid" = "geneId"))
table(markers_alpha$cluster)

# calculate alpha ourselves because there are a few mistakes in the precalculated data
markers_alpha %<>% mutate(alpha = 1 - (Ds*Pn)/(Dn*Ps))
markers_alpha <- markers_alpha[,-18]

library(ggplot2)
markers_alpha_pos <- subset(markers_alpha, alpha > 0 & alpha <= 1 )
table(markers_alpha_pos$cluster)

alpha_plot_mel <- ggplot(markers_alpha, aes(x = alpha))
alpha_plot_mel +
  geom_density(aes(fill = cluster),size = 0.3, alpha = 0.5)+
  theme(legend.position="right")+
  labs(fill = "cell type") +
  scale_x_continuous(limits = c(-5, 1)) +
  scale_fill_manual(values = my_colors)+
  theme_minimal(base_size = 10.5)

kruskal.test(alpha ~ cluster, data = markers_alpha)
pairwise.wilcox.test(markers_alpha_pos$alpha, markers_alpha_pos$cluster,
                     p.adjust.method = "BH")
aggregate(alpha ~ cluster, markers_alpha, mean)

## how do non-SFPs look w/ alpha values
mk_nonSFPs <- subset(markers_alpha, !`##primary_FBid` %in% sfps$V2)
table(mk_nonSFPs$cluster)

mk_nonSFPs_pos <- subset(mk_nonSFPs, alpha > 0 )
table(mk_nonSFPs_pos$cluster)

alpha_plot_nonSFP <- ggplot(mk_nonSFPs, aes(x = alpha))
alpha_plot_nonSFP +
  geom_density(aes(fill = cluster), size = 0.3, alpha = 0.5)+
  theme(legend.position="right")+
  scale_x_continuous(limits = c(-5, 1))+
  scale_fill_manual(values = my_colors)+
  theme_minimal(base_size = 10.5) + labs(fill = "cell type") 
kruskal.test(alpha ~ cluster, data = mk_nonSFPs)
pairwise.wilcox.test(mk_nonSFPs$alpha, mk_nonSFPs$cluster,
                     p.adjust.method = "BH")
aggregate(alpha ~ cluster, mk_nonSFPs, median)

# comparing SFPs vs non-SFPs in alpha values
mk_SFP <- subset(markers_alpha, `##primary_FBid` %in% SFPs$V1)
table(mk_SFP$cluster)

mk_SFP$type <- "SFP"
mk_nonSFPs$type <- "other"

SFPvsOther_alpha <- rbind(mk_SFP, mk_nonSFPs)
ggplot(SFPvsOther_alpha, aes(x = alpha)) +
  geom_density(aes(fill = type), size = 0.3, alpha = 0.5)+
  theme(legend.position="right")+
  scale_x_continuous(limits = c(-5, 1))+
  theme_minimal(base_size = 10.5) +
  scale_fill_brewer(palette = "Set1")

table(subset(SFPvsOther_alpha, alpha > 0)$type)
table(subset(SFPvsOther_alpha, alpha < 0)$type)

91 / (91 + 34) # SFP
177 / (177 + 260) # nonSFP

fisher.test(as.matrix(data.frame(c(91, 91 + 34), c(177, 177 + 260))))

aggregate(alpha ~ type, subset(SFPvsOther_alpha, alpha > -5), median)
kruskal.test(alpha ~ type, data = subset(SFPvsOther_alpha, alpha > 0))
### examining the ratios of positive to negative alpha values

MC_pos <- length(markers_alpha$alpha[markers_alpha$alpha > 0 & markers_alpha$cluster == "MC"])
MC_neg <- length(markers_alpha$alpha[markers_alpha$alpha <= 0 & markers_alpha$cluster == "MC"])
SC_pos <- length(markers_alpha$alpha[markers_alpha$alpha > 0 & markers_alpha$cluster == "SC"])
SC_neg <- length(markers_alpha$alpha[markers_alpha$alpha <= 0 & markers_alpha$cluster == "SC"])
ED_pos <- length(markers_alpha$alpha[markers_alpha$alpha > 0 & markers_alpha$cluster == "EDC"])
ED_neg <- length(markers_alpha$alpha[markers_alpha$alpha <= 0 & markers_alpha$cluster == "EDC"])

alpha_proportions <- t(as.matrix(data.frame(c(MC_pos, 
                                              MC_neg), 
                                            c(SC_pos, 
                                              SC_neg), 
                                            c(ED_pos, 
                                              ED_neg), 
                                            row.names = c("positive", "negative")))) 
rownames(alpha_proportions) <- c("MC", "SC", "ED")

as.data.frame(alpha_proportions) %>% mutate(ratio = 
                                              round(positive/(negative + positive), 2))

library(RVAideMemoire)
G.test(alpha_proportions)
fisher.test(alpha_proportions)
# G = 21.36, df = 2, p-value = 2.3e-05
fisher.multcomp(alpha_proportions)
# MC have elevated ratio of genes w/ evidence of positive selection

# what if we remove the SFPs?
MC_noSFP_pos <- length(mk_nonSFPs$alpha[mk_nonSFPs$alpha > 0 & mk_nonSFPs$cluster == "MC"])
MC_noSFP_neg <- length(mk_nonSFPs$alpha[mk_nonSFPs$alpha <= 0 & mk_nonSFPs$cluster == "MC"])
SC_noSFP_pos <- length(mk_nonSFPs$alpha[mk_nonSFPs$alpha > 0 & mk_nonSFPs$cluster == "SC"])
SC_noSFP_neg <- length(mk_nonSFPs$alpha[mk_nonSFPs$alpha <= 0 & mk_nonSFPs$cluster == "SC"])
ED_noSFP_pos <- length(mk_nonSFPs$alpha[mk_nonSFPs$alpha > 0 & mk_nonSFPs$cluster == "EDC"])
ED_noSFP_neg <- length(mk_nonSFPs$alpha[mk_nonSFPs$alpha <= 0 & mk_nonSFPs$cluster == "EDC"])

alpha_proportions_noSFP <- t(as.matrix(data.frame(c(MC_noSFP_pos, 
                                                    MC_noSFP_neg), 
                                                  c(SC_noSFP_pos, 
                                                    SC_noSFP_neg), 
                                                  c(ED_noSFP_pos, 
                                                    ED_noSFP_neg), 
                                                  row.names = c("positive", "negative")))) 
rownames(alpha_proportions_noSFP) <- c("MC", "SC", "ED")

as.data.frame(alpha_proportions_noSFP) %>% mutate(ratio = 
                                                    round(positive/(negative + positive), 2))

fisher.test(alpha_proportions_noSFP)
# G = 2.3818, df = 2, p-value = 0.4076

# so take-away is that MC pattern is driven by SFP enrichments
# in the MC. but that the distribution of the number of sites
# estimated to be fixed under selection is not driven by SFPs.

#######################################################
#### intersect DE with alpha values and SFP status ####
## wondering if genes that are DE tend to have higher alpha values than
## those that do not show DE?
#######################################################

## get list of genes DE in mel v sim 
all.DE.1.melvsim <- unique(c(rownames(main.DE.melvsim.sig),
                             rownames(secondary.DE.melvsim.sig),
                             rownames(ED.DE.melvsim.sig)))

## find out expression level of the DE genes, to get comparable non-DE genes
DE.genes.data <- as.data.frame(GetAssayData(msy.so))[rownames(as.data.frame(GetAssayData(msy.so))) %in% all.DE.1.melvsim,]
DE.genes.data %<>% rownames_to_column("gene")
DE.genes.data %<>% mutate(counts = rowSums(.[-1]))
summary(DE.genes.data$counts)
### so the min number of counts for a DE gene is 62.87
table(rowSums(as.data.frame(GetAssayData(msy.so))) > 62.87)
## that gives us a set of 4948 expressed genes expressed at least
## as highly as the lowest-expressed DE gene
expr.genes.data <- as.data.frame(GetAssayData(msy.so))[rowSums(as.data.frame(GetAssayData(msy.so))) > 62.87,]
expr.genes.data %<>% rownames_to_column("gene")
expr.genes.data %<>% mutate(counts = rowSums(.[-1]))
summary(expr.genes.data$counts)
## so we have two groups to compare for alpha values and dN + omega too if we want.
## DE genes, and all genes expressed at that level. thinking this is analogous to a GO 
## or any other enrichment type analysis

DE.genes.names <- data.frame(DE.genes.data$gene, DE.genes.data$counts, "DE") %>% setNames(c("gene", "counts", "type"))
nonDE.genes.names <- data.frame(expr.genes.data$gene[!expr.genes.data$gene %in% DE.genes.data$gene], 
                                expr.genes.data[!expr.genes.data$gene %in% DE.genes.data$gene,]$counts,
                                "non-DE") %>% setNames(c("gene", "counts", "type"))
DE.vs.expr <- rbind(DE.genes.names, nonDE.genes.names)

# join gene name to FBid and FBid to alpha vals
mk %<>% mutate(alpha = 1 - (Ds*Pn)/(Dn*Ps))

DE.vs.expr <- inner_join(DE.vs.expr, synonyms, by = c("gene" = "current_symbol"))
DE.vs.expr <- inner_join(DE.vs.expr, mk, by = c("##primary_FBid" = "geneId"))

## look at the data 
aggregate(alpha ~ type, subset(DE.vs.expr), median)
kruskal.test(subset(DE.vs.expr, alpha > 0), alpha ~ type)
ggplot(DE.vs.expr, aes(x = alpha)) +
  geom_density(aes(fill = type), size = 0.3, alpha = 0.5)+
  theme(legend.position="right")+
  scale_x_continuous(limits = c(-5, 1))+
  theme_minimal(base_size = 10.5)+
  scale_fill_brewer(palette = "Set1")

## DE gene positive alpha median is 0.53 vs nonDE gene 0.45,
## sig diff w/ kruskal test

DE_pos    <- nrow(subset(DE.vs.expr, type == "DE" & alpha >  0))
DE_neg    <- nrow(subset(DE.vs.expr, type == "DE" & alpha <= 0))
nonDE_pos <- nrow(subset(DE.vs.expr, type == "non-DE" & alpha >  0))
nonDE_neg <- nrow(subset(DE.vs.expr, type == "non-DE" & alpha <= 0))

contingency <- data.frame(c(DE_pos, nonDE_pos), c(DE_neg, nonDE_neg))
contingency %<>% setNames(c("positive", "negative"))

contingency %>% mutate(ratio = positive / (positive + negative))
G.test(as.matrix(contingency))
# and we can see that DE genes are not more likely to have a positive alpha than nonDE genes

# is expression level confounded with alpha
ggplot(subset(DE.vs.expr, alpha > 0 & alpha <= 1), aes(x = log(counts), y = alpha)) +
  geom_point()

lm(scale(alpha) ~ scale(log(counts)), subset(DE.vs.expr, alpha > 0 & alpha <= 1))
# not very the slope is 0.145?

### ok what about logFC mel vs sim and alpha
DE.vs.expr.logFCs <- inner_join(DE.vs.expr, all.DE.1, by = "gene")[,c(1:4,9,12:19,23,27)]
DE.alpha.logFC <- subset(DE.vs.expr.logFCs, type == "DE")

ggplot(subset(DE.alpha.logFC, alpha > 0), aes(x = alpha, y = abs(MC_ms))) +
  geom_point() +
  theme_minimal() +
  labs(x = "alpha", y = "abs(log(mel/sim)) MC")
ggplot(subset(DE.alpha.logFC, alpha > 0), aes(x = alpha, y = abs(SC_ms))) +
  geom_point() +
  theme_minimal() +
  labs(x = "alpha", y = "abs(log(mel/sim)) SC")
ggplot(subset(DE.alpha.logFC, alpha > 0), aes(x = alpha, y = abs(EDC_ms))) +
  geom_point() +
  theme_minimal() +
  labs(x = "alpha", y = "abs(log(mel/sim)) EDC")

cor(subset(DE.alpha.logFC, alpha > -10)$alpha, abs(subset(DE.alpha.logFC, alpha > -10)$MC_ms))
cor(subset(DE.alpha.logFC, alpha > -10)$alpha, abs(subset(DE.alpha.logFC, alpha > -10)$SC_ms))
cor(subset(DE.alpha.logFC, alpha > -10)$alpha, abs(subset(DE.alpha.logFC, alpha > -10)$EDC_ms))


#######################################
#### unannotated and de novo genes ####
#######################################

# merged markers
markers_merged_nothresh <- FindAllMarkers(
  object = mel.so, 
  only.pos = TRUE, 
  min.pct = 0, 
  logfc.threshold = 0,
  return.thresh = 1
)

## as marker genes?
markers_merged_DU <- subset(markers_merged, gene %in% denovo_unannotated_geneIDs)
markers_merged_DU

markers_merged_sig_DU <- subset(markers_merged_sig, gene %in% denovo_unannotated_geneIDs)
markers_merged_sig_DU

markers_merged_DU_nothresh <- subset(markers_merged_nothresh, gene %in% unannotated_geneIDs)
markers_merged_DU_nothresh

## get proper adjusted p-value, rather than bonferonni correcting by divinding by all genes,
## multiply by number of unannotated genes investigated, 11

markers_merged_DU_nothresh %<>% mutate(p_val_adj = p_val * 11)

## get percent expressing in each cell type
# first get brcodes of each cell type
MC<- names(Idents(mel.so)[Idents(mel.so) == "MC"])
SC<- names(Idents(mel.so)[Idents(mel.so) == "SC"])
EDC<- names(Idents(mel.so)[Idents(mel.so) == "EDC"])

counts_unannotated %<>% rownames_to_column("gene")

counts_unannotated %<>% mutate(pct.MC=rowSums(.[MC]!=0)/length(MC))
counts_unannotated %<>% mutate(pct.SC=rowSums(.[SC]!=0)/length(SC))
counts_unannotated %<>% mutate(pct.EDC=rowSums(.[EDC]!=0)/length(EDC))

View(counts_unannotated[,c(1,1169:1171)])

## UMAP
FeaturePlot(mel.so, features = denovo_unannotated_geneIDs, pt.size = 0.8)
FeaturePlot(mel.so, features = c("TRINITY-DN2695-c0-g1-i1"), pt.size = 0.8)
VlnPlot(mel.so, features = "TRINITY-DN2695-c0-g1-i1", cols = c("slategrey", "slategrey","slategrey")) + theme(legend.position = 'none') 

# SC marker TRINITY-DN2695-c0-g1-i1 is 650 bp from a lncRNA. Is it expressed similarly?
FeaturePlot(mel.so, features = "lncRNA:CR45370")
FeaturePlot(mel.so, features = "Spn28B")
# no, it is apparently not expressed in the dataset, nice

# how does the avg expression level of these candidate genes compare to the distributions of
# expression among all genes and among cell type markers?
avgexp_unannotated <- data.frame(AverageExpression(mel.so, features = unannotated_geneIDs))
avgexp_allexpgene <- data.frame(AverageExpression(mel.so,features = rownames(mel.so@assays$RNA@counts)))

avgexp_SC_markers <- data.frame(AverageExpression(mel.so, features =  subset(markers_all_sig, markers_all_sig$cluster == "SC")$gene))
avgexp_MC_markers <- data.frame(AverageExpression(mel.so, features =  subset(markers_all_sig, markers_all_sig$cluster == "MC")$gene))
avgexp_EDC_markers <- data.frame(AverageExpression(mel.so, features =  subset(markers_all_sig, markers_all_sig$cluster == "EDC")$gene))

## examine distributions of unannotated transcripts among markers, and all genes
quantile(avgexp_allexpgene$RNA.MC, seq(0,1,0.05))
quantile(avgexp_allexpgene$RNA.SC, seq(0,1,0.05))
quantile(avgexp_allexpgene$RNA.EDC, seq(0,1,0.05))

quantile(avgexp_MC_markers$RNA.MC, seq(0,1,0.05))
quantile(avgexp_SC_markers$RNA.SC, seq(0,1,0.05))
quantile(avgexp_EDC_markers$RNA.EDC, seq(0,1,0.05))

avgexp_allexpgene$type <- "all genes"
avgexp_MC_markers$type <- "MC markers"
avgexp_SC_markers$type <- "SC markers"
avgexp_EDC_markers$type <- "EDC markers"

# plot SC
avgexp_SC_all <- rbind(avgexp_allexpgene, avgexp_SC_markers)
avgexp_unannotated_SC <- avgexp_unannotated %>% subset(rownames(avgexp_unannotated) %in% c("TRINITY-DN10097-c0-g1-i1",
                                                                                           "TRINITY-DN2695-c0-g1-i1"))
avgexp_unannotated_SC %<>% rownames_to_column("gene")

unannotated_SC_plot <- ggplot(avgexp_SC_all, aes(x = log(RNA.SC), fill = type)) + 
        geom_density(alpha = 0.7, color = "darkgrey") +
        scale_fill_manual(values = c("grey", my_colors[2])) +
        geom_vline(data = avgexp_unannotated_SC,
             aes(xintercept = log(RNA.SC)),
             linetype = 2,
             color = "grey39") +
  labs(fill = "") + 
  annotate("text", x = 3.75,
                          y = c(.35,.325),
                          label = c("DN10097", "DN2695"),
                          angle = 20,
                          hjust = "left",
                          size = 3.4) +
        theme_minimal() +
        xlab("log(SC expression)")
unannotated_SC_plot
# plot MC
avgexp_MC_all <- rbind(avgexp_allexpgene, avgexp_MC_markers)
avgexp_unannotated_MC <- avgexp_unannotated %>% subset(rownames(avgexp_unannotated) %in% c("TRINITY-DN4707-c0-g1-i1",
                                                                                           "TRINITY-DN8354-c0-g1-i1",
                                                                                           "TRINITY-DN35169-c0-g1-i1",
                                                                                           "TRINITY-DN11110-c0-g1-i1",
                                                                                           "TRINITY-DN2736-c0-g1-i4",
                                                                                           "TRINITY-DN5813-c0-g1-i4",
                                                                                           "TRINITY-DN818-c0-g1-i10"))
avgexp_unannotated_MC %<>% rownames_to_column("gene")

unannotated_MC_plot <- ggplot(avgexp_MC_all, aes(x = log(RNA.MC), fill = type)) + 
  geom_density(alpha = 0.7, color = "darkgrey") +
  scale_fill_manual(values = c("grey", my_colors[1])) +
  geom_vline(data = avgexp_unannotated_MC,
             aes(xintercept = log(RNA.MC)),
             linetype = 2,
             color = "grey39") +
  labs(fill = "") + 
  annotate("text", x = 3.2,
                   y = seq(0.4,0.25, length.out = 7),
                   label = c("DN8354",
                             "DN4707",
                             "DN35169",
                             "DN11110",
                             "DN5813",
                             "DN2736",
                             "DN818"),
                  angle = 20,
                  hjust = "left",
                  vjust = "bottom",
                  size = 3.4)+
  theme_minimal() +
  xlab("log(MC expression)")
unannotated_MC_plot

# plot EDC
avgexp_EDC_all <- rbind(avgexp_allexpgene, avgexp_EDC_markers)
avgexp_unannotated_EDC <- avgexp_unannotated %>% subset(rownames(avgexp_unannotated) %in% c("TRINITY-DN16089-c0-g1-i1",
                                                                                            "TRINITY-DN10930-c0-g1-i3"))
avgexp_unannotated_EDC %<>% rownames_to_column("gene")

unannotated_EDC_plot <- ggplot(avgexp_EDC_all, aes(x = log(RNA.EDC), fill = type)) + 
  geom_density(alpha = 0.7, color = "darkgrey") +
  scale_fill_manual(values = c("grey", my_colors[3])) +
  geom_vline(data = avgexp_unannotated_EDC,
             aes(xintercept = log(RNA.EDC)),
             linetype = 2,
             color = "grey39") +
  labs(fill = "") + 
  annotate("text", x = 3.1,
                   y = c(0.325, 0.302),
                   label = c("DN10930", "DN16089"),
                   angle = 20,
                   hjust = "left",
                   size = 3.4) +
  theme_minimal() +
  xlab("log(EDC expression)")
unannotated_EDC_plot

plot_grid(unannotated_MC_plot, unannotated_SC_plot, unannotated_EDC_plot, ncol = 1)

# FeaturePlots of unannotated genes
unannotated_umap_A <- mel.so %>% FeaturePlot(unannotated_geneIDs[8], order = T, cols = c("lightgrey", "darkcyan"), combine = F, pt.size = 0.3, label = T, repel = T) 
unannotated_umap_B <- mel.so %>% FeaturePlot(unannotated_geneIDs[c(11,7,2,4,3,6,9,10,1,5)], order = T, cols = c("lightgrey", "darkcyan"), combine = F, pt.size = 0.3)

unannotated_umap <- c(unannotated_umap_A, unannotated_umap_B)
unannotated_umap %<>% lapply(FUN = function(x) x + theme(axis.title = element_text(size = 11),
                                                          axis.text = element_text(size = 11),
                                                         plot.title = element_text(size = 11, face = "plain"),
                                                         legend.position = "none"))

unannotated_umap[c(2:5,7:11)] %<>% lapply(FUN = function(x) x + theme(axis.title.y = element_blank()))
unannotated_umap[c(1:5)] %<>% lapply(FUN = function(x) x + theme(axis.title.x = element_blank()))

                                                                                                
plot_grid(plotlist = unannotated_umap[1:5], nrow = 1)
plot_grid(plotlist = unannotated_umap[6:11], nrow = 1)
```
