## Supplementary material for "Single-nucleus transcriptomes reveal functional and evolutionary properties of cell types in the Drosophila accessory gland": Dataset S10: SoupX.R

```
library(SoupX)
library(Seurat)
library(dplyr)
library(Matrix)
library(ggplot2)
library(tidyr)
library(plotly)
library(stringr)
library(RColorBrewer)
library(scales)
library(data.table)

###########################################################
### 1A. do preliminary clustering in for use with SoupX ###
###########################################################

mel.raw <- read.table("mel_cell_background.tsv", row.names = 1, header = T) 
mel.so <- CreateSeuratObject(mel.raw, min.cells = 3, min.features = 100)

# loading the gene expression matrix plus the empty droplets that will later be 
# used in SoupX. this is immaterial for seurat however because all the empty drops are excluded by 
# using min.features = 100 above. this way i wont have to load a separate table for soupX, just re-use
# the mel.raw matrix

FeatureScatter(mel.so, feature1 = "nCount_RNA", feature2 = "nFeature_RNA")

# Show 5% qunatiles for number of genes per cell
do.call("cbind", tapply(mel.so@meta.data$nFeature_RNA,mel.so@active.ident,quantile,probs=seq(0,1,0.05)))
# Show 5% qunatiles for number of counts per cell per sample
do.call("cbind", tapply(mel.so@meta.data$nCount_RNA,mel.so@active.ident,quantile,probs=seq(0,1,0.05)))

mel.so <- subset(mel.so, nCount_RNA <= 3000) 

# next normalize the data
mel.so <- NormalizeData(
  object = mel.so,
  normalization.method = "LogNormalize",
  scale.factor = 10000)

# FindVariableFeatures 

mel.so <- FindVariableFeatures(
  object = mel.so,
  selection.method = "vst")

# scale
mel.so <- ScaleData(object = mel.so)
# PCA
mel.so <- RunPCA(object = mel.so)
DimPlot(object = mel.so, dims = c(1,2), reduction = "pca")
DimPlot(object = mel.so, dims = c(1,3), reduction = "pca")
ElbowPlot(mel.so)

# JackStraw
mel.so <- JackStraw(
  object = mel.so, dims = 40)
mel.so <- ScoreJackStraw(mel.so, dims = 1:40)

JackStrawPlot(object = mel.so, dims = 1:40)
# observe plot to decide what PCs to use
use.pcs = c(1:16)

#UMAP

mel.so <- RunUMAP(mel.so, reduction = "pca", dims = use.pcs)
mel.so <- FindNeighbors(mel.so, reduction = "pca", dims = use.pcs)

# look at cluster nums at a series of resolutions
mel.so <- FindClusters(
  object = mel.so,
  resolution = seq(0.2,1.4,0.1),
  verbose = FALSE)
sapply(grep("res",colnames(mel.so@meta.data),value = TRUE),
       function(x) length(unique(mel.so@meta.data[,x])))
# choose resolution
Idents(mel.so) <- "RNA_snn_res.0.9"

# UMAP Visualization
DimPlot(mel.so, reduction = "umap", label = T, pt.size = 2) +
    theme_minimal(base_size = 15)

FeaturePlot(mel.so, features = c('nCount_RNA'), pt.size=2)

# marker genes 
markers_all <- FindAllMarkers(
  object = mel.so, 
  only.pos = TRUE, 
  min.pct = 0.25, 
  thresh.use = 0.25
)

dim(markers_all)
head(markers_all)
 
# heirarchical cluster tree
mel.so <- BuildClusterTree(
  mel.so, dims = use.pcs)

PlotClusterTree(mel.so)

# merging clusters into known cell types, at least for the purpose of cleaning up with SoupX
## based on UMAP and heirarchical tree
## merging main cell sub-states / sub-types into one
mel.so.merged <- RenameIdents(
  object = mel.so,
  '1' = '0', '2' = '0', '3' = '0', '4' = '0'
)

# naming the cell types based on marker genes
mel.so.merged <- RenameIdents(
  object = mel.so.merged,
  '0' = 'main cells', '6' = 'secondary cells', '5' = 'ejaculatory duct cells'
)

table(Idents(mel.so.merged))
# main cells        secondary cells     ejaculatory duct cells 
#       1055                     50                         62 

#####################################
## 1B. ESTIMATE SOUP TRANSCRIPTOME ##
#####################################

# need to filter out the cells that I already treated as doublets from prelim clustering
good_cells <- rownames(mel.so@meta.data)
# now define the empty droplets
empty_drops <- colnames(mel.raw[,1296:2295])
# combine our good cells w/ the drops and filter the raw dataset
good_cells_plus_empties <- c(good_cells, empty_drops)
mel.filtered <- mel.raw[,colnames(mel.raw) %in% good_cells_plus_empties]

# create the soupchannel, estimate the soup transcriptome
mel.sc <- SoupChannel(tod = as.matrix(mel.filtered), toc = as.matrix(mel.filtered[,c(1:1167)]), soupRange = c(0,100))

#################################
## 2. ESTIMATE SOUP PERCENTAGE ##
#################################

# check barcodes between seurat object and soup channel
head(mel.sc$metaData)
head(mel.so.merged@meta.data[,1:3])
tail(mel.sc$metaData)
tail(mel.so.merged@meta.data[,1:3])

# there is some slight decrease in nUMI (like a few UMIs) between soup channel and seurat object. 
# This is expected because in seurat I eliminated genes with < 3 UMI total from analysis

# add cell type to the metadata of seurat object
mel.so.merged$CellType <- Idents(mel.so.merged)

# add cell type data to soup channel
mel.sc <- setClusters(sc = mel.sc, clusters = mel.so.merged$CellType)

# look at markers as sanity check
quickMarkers(toc = as.matrix(mel.sc$toc), clusters = as.vector(mel.sc$metaData$clusters))

# estimate contamination fraction
mel.sc <- autoEstCont(mel.sc, priorRho = 0.15)

#165 genes passed tf-idf cut-off and 69 soup quantile filter.
#Using 74 independent estimates of rho.
#Estimated global rho of 0.17

################################
#### 3. REMOVE CONTAMINATION ###
################################  

mel.cleaned <- adjustCounts(mel.sc)

### inspect the changes in expression

genecounts.raw = rowSums(mel.sc$toc > 0)
genecounts.filtered = rowSums(mel.cleaned > 0)
mostZeroed = tail(sort((genecounts.raw - genecounts.filtered)/genecounts.raw), n = 300)
mostZeroed # which genes are the most zeroed?

# genes zeroed in every cell: 
rowSums(mel.sc$toc[c(names(mostZeroed)[194:300]),])

# genes zeroed in a fravtion of cells:
rowSums(mel.sc$toc[c(names(mostZeroed)[1:193]),])
rowSums(mel.cleaned[c(names(mostZeroed)[1:193]),])
# some are highly expressed, some are very lowly expressed. No in-between. high expressed ones could be 
# important markers that have a lot of background. eg. Dup99B - very high expressed ED cell marker 
# that shows up as background in most main and secondary cells as well.
FeaturePlot(mel.so, features = "FBgn0250832", pt.size=2) +
  theme_minimal(base_size = 15) # Dup99B
  
FeaturePlot(mel.so, features = "FBgn0003034", pt.size=2) +
  theme_minimal(base_size = 15) # Sex Peptide

FeaturePlot(mel.so, features = "FBgn0023415", pt.size=2) +
  theme_minimal(base_size = 15) # Acp32CD

# another type of gene mostly zeroedd are those that are kind of sporadically expressed, but at 
# relatively high levels on a per-cell basis. eg. lncRNA:roX2 and Obp58b
FeaturePlot(mel.so, features = "FBgn0034768", pt.size=2) # obp
FeaturePlot(mel.so, features = "FBgn0019660", pt.size=2) # lncRNA:rox2

### Visualizing SoupX changes in expression 
mel.umap <- mel.so@reductions$umap@cell.embeddings  # making a dataframe with UMAP coordinates

changemap_Dup99B <- plotChangeMap(mel.sc, mel.cleaned, DR = mel.umap, "FBgn0250832") + 
              scale_color_gradient(low = "#99D8C9", high = "#00441B") + 
              theme_minimal(base_size = 15) + 
              labs(x = "UMAP 1", y = "UMAP 2", color = "soup fraction", title = "Dup99B") +
              geom_point(aes(col=relChange),size=2) 
changemap_Dup99B

changemap_SP <- plotChangeMap(mel.sc, mel.cleaned, DR = mel.umap, "FBgn0003034") + 
  scale_color_gradient(low = "#99D8C9", high = "#00441B") + 
  theme_minimal(base_size = 15) + 
  labs(x = "UMAP 1", y = "UMAP 2", color = "soup fraction", title = "Sex Peptide") +
  geom_point(aes(col=relChange),size=2) 
changemap_SP

changemap_Spn77Bc <- plotChangeMap(mel.sc, mel.cleaned, DR = mel.umap, "FBgn0036970") + 
  scale_color_gradient(low = "#99D8C9", high = "#00441B") + 
  theme_minimal(base_size = 15) + 
  labs(x = "UMAP 1", y = "UMAP 2", color = "soup fraction", title = "Spn77Bc") +
  geom_point(aes(col=relChange),size=2) 
changemap_Spn77Bc

changemap_Acp32CD <- plotChangeMap(mel.sc, mel.cleaned, DR = mel.umap, "FBgn0023415") + 
  scale_color_gradient(low = "#99D8C9", high = "#00441B") + 
  theme_minimal(base_size = 15) + 
  labs(x = "UMAP 1", y = "UMAP 2", color = "soup fraction", title = "Acp32CD") +
  geom_point(aes(col=relChange),size=2) 
changemap_Acp32CD

changemap_Acp36DE <- plotChangeMap(mel.sc, mel.cleaned, DR = mel.umap, "FBgn0011559") + 
  scale_color_gradient(low = "#99D8C9", high = "#00441B") + 
  theme_minimal(base_size = 15) + 
  labs(x = "UMAP 1", y = "UMAP 2", color = "soup fraction", title = "Acp36DE") +
  geom_point(aes(col=relChange),size=2) 
changemap_Acp36DE

# export the soup-corrected reads to a matrix
write.table(mel.cleaned, "mel.cleaned.tsv", sep = "\t", quote = F, col.names = NA)

#
##
#####
#########
################
###########################
###########################################
## soupX cleanup for simulans and yakuba ##
###########################################
###########################
################
#########
#####
##
#

################
## D SIMULANS ##
################

###########################################################
### 1A. do preliminary clustering in for use with SoupX ###
###########################################################

sim.raw <- read.table("sim_cells_background.tsv", row.names = 1, header = T)
# using the FBgn since it will be simpler to do the ortholog annotation later. Doesn't really help too much
# to have gene names during single-species restricted clustering becuase the simulans gene name
# annotation is extremely poor  anyways (despite ortholog annotation being good).
sim.so <- CreateSeuratObject(sim.raw, min.cells = 3, min.features = 100)

FeatureScatter(sim.so, feature1 = "nCount_RNA", feature2 = "nFeature_RNA", pt.size = 2)

# Show 5% qunatiles for number of genes per cell
do.call("cbind", tapply(sim.so@meta.data$nFeature_RNA,sim.so@active.ident,quantile,probs=seq(0,1,0.05)))
# Show 5% qunatiles for number of counts per cell per sample
do.call("cbind", tapply(sim.so@meta.data$nCount_RNA,sim.so@active.ident,quantile,probs=seq(0,1,0.05)))

sim.so <- subset(sim.so, nCount_RNA <= 5000)

# next normalize the data
sim.so <- NormalizeData(
  object = sim.so,
  normalization.method = "LogNormalize",
  scale.factor = 10000)

# FindVariableFeatures 

sim.so <- FindVariableFeatures(
  object = sim.so,
  selection.method = "vst")

# scale
sim.so <- ScaleData(object = sim.so)
# PCA
sim.so <- RunPCA(object = sim.so)
DimPlot(object = sim.so, dims = c(1,2), reduction = "pca")
DimPlot(object = sim.so, dims = c(1,3), reduction = "pca")
ElbowPlot(sim.so)

# JackStraw
sim.so <- JackStraw(
  object = sim.so, dims = 40)
sim.so <- ScoreJackStraw(sim.so, dims = 1:40)

JackStrawPlot(object = sim.so, dims = 1:40)
# observe plot to decide what PCs to use
use.pcs = c(1:8)

#UMAP

sim.so <- RunUMAP(sim.so, reduction = "pca", dims = use.pcs)
sim.so <- FindNeighbors(sim.so, reduction = "pca", dims = use.pcs)

# look at cluster nums at a series of resolutions
sim.so <- FindClusters(
  object = sim.so,
  resolution = seq(0.2,1.4,0.1),
  verbose = FALSE)
sapply(grep("res",colnames(sim.so@meta.data),value = TRUE),
       function(x) length(unique(sim.so@meta.data[,x])))
# choose resolution
Idents(sim.so) <- "RNA_snn_res.0.7"

# UMAP Visualization
DimPlot(sim.so, reduction = "umap", label = T, pt.size = 1)

FeaturePlot(sim.so, features = c('nCount_RNA'), pt.size=1)
FeaturePlot(sim.so, features = c('nFeature_RNA'), pt.size=1)

# marker genes 
markers_sim_all <- FindAllMarkers(
  object = sim.so, 
  only.pos = TRUE, 
  min.pct = 0.25, 
  thresh.use = 0.25
)

dim(markers_sim_all)
head(markers_sim_all)

table(Idents(sim.so))

sim.so$CellType <- Idents(sim.so)

ggplot(data = sim.so@meta.data, aes(x = CellType, y = nCount_RNA)) +
        geom_violin(position = position_dodge(width = 0.9))  + 
        geom_jitter()

# heirarchical cluster tree
sim.so <- BuildClusterTree(
 sim.so, dims = use.pcs)

PlotClusterTree(sim.so)

FeaturePlot(sim.so, features = c("FBgn0189236", "FBgn0191744", "FBgn0041931"))
# FBgn0189236 = Dup99B (duct cell marker)
# FBgn0191744 = abd-A (sec cell marker)
# FBgn0041931 = Acp36DE (main cell marker)

# merging clusters into known cell types, at least for the purpose of cleaning up with SoupX
## based on UMAP and heirarchical tree
## merging main cell sub-states / sub-types into one
sim.so.merged <- RenameIdents(
  object = sim.so,
  '1' = '0', '3' = '0', '4' = '0', '5' = '0',
  '2' = '7'
)

# naming the cell types based on marker genes
sim.so.merged <- RenameIdents(
  object = sim.so.merged,
  '0' = 'main cells', '6' = 'secondary cells', '7' = 'ejaculatory duct cells', '8' = 'muscle cells'
)
table(Idents(sim.so.merged))
#ejaculatory duct cells     main cells        secondary cells       muscle cells 
#364                        1690              41                    20 

# interesting to find muscle cells in simulans, but not in melanogaster
sim.so.merged$CellType <- Idents(sim.so.merged)

ggplot(data = sim.so.merged@meta.data, aes(x = CellType, y = nCount_RNA)) +
  geom_violin(position = position_dodge(width = 0.9))  + 
  geom_jitter(size = 1) + 
  theme_minimal(base_size = 14)

FeatureScatter(sim.so.merged, feature1 = "nCount_RNA", feature2 = "nFeature_RNA", pt.size = 2,) + 
  theme_minimal() +
  labs(title = NULL)

# another sort of interesting thing - muscle cells tend to have a lower number of features to number of UMI
# ratio than MC, SC, & EDC. this squares with my idea of the biology - muscle cells do not have this 
# complex set of secreted proteins and related co-expressed genes involved in the secretory pathways - 
# so you could argue that we might expect their transcriptome to be "simpler"

ggplot(data = sim.so.merged@meta.data, aes(x = CellType, y = nFeature_RNA/nCount_RNA)) +
  geom_violin(position = position_dodge(width = 0.9))  + 
  geom_jitter(size = 1) + 
  theme_minimal(base_size = 14)

#####################################
## 1B. ESTIMATE SOUP TRANSCRIPTOME ##
#####################################

# need to filter out the cells that I already treated as doublets from prelim clustering
good_cells <- rownames(sim.so.merged@meta.data)
empty_drops <- colnames(sim.raw[,2295:3294])
good_cells_plus_empties <- c(good_cells, empty_drops)
sim.filtered <- sim.raw[,colnames(sim.raw) %in% good_cells_plus_empties]

# create the soupchannel, estimate the soup transcriptome
sim.sc <- SoupChannel(tod = sim.filtered, toc = as.matrix(sim.filtered[,1:2115]), keepDroplets = T, soupRange = c(0,100))

View(sim.sc$soupProfile)
# seems to check out

#################################
## 2. ESTIMATE SOUP PERCENTAGE ##
#################################

# check barcodes between seurat object and soup channel
head(sim.sc$metaData)
head(sim.so.merged@meta.data[,1:3])
tail(sim.sc$metaData)
tail(sim.so.merged@meta.data[,1:3])

# there is some slight decrease in nUMI (like a few UMIs) between soup channel and seurat object. 
# This is expected because in seurat I eliminated genes with < 3 UMI total from analysis

# add cell type to the metadata of seurat object
sim.so.merged$CellType <- Idents(sim.so.merged)

# add cell type data to soup channel
sim.sc <- setClusters(sc = sim.sc, clusters = sim.so.merged$CellType)

# look at markers as sanity check
quickMarkers(sim.sc$toc, clusters = sim.sc$metaData$clusters, N = 10)

# estimate contamination fraction
sim.sc <- autoEstCont(sim.sc, priorRho = 0.135) # basing my prior on the estimate of mel contamination %
#139 genes passed tf-idf cut-off and 52 soup quantile filter.
#Using 65 independent estimates of rho.
#Estimated global rho of 0.16

################################
#### 3. REMOVE CONTAMINATION ###
################################  

sim.cleaned <- as.data.frame(adjustCounts(sim.sc))

### inspect the changes in expression

genecounts.raw = rowSums(sim.sc$toc > 0)
genecounts.filtered = rowSums(sim.cleaned > 0)
mostZeroed = tail(sort((genecounts.raw - genecounts.filtered)/genecounts.raw), n = 300)
mostZeroed # which genes are the most zeroed?

length(subset(mostZeroed, mostZeroed == 1.0)) # 109 genes zeroed in every cell

# genes zeroed in every cell: 
rowSums(sim.sc$toc[c(names(mostZeroed)[192:300]),])
# appear not to be highly expressed, probably not important markers

# genes zeroed in a fravtion of cells:
rowSums(sim.sc$toc[c(names(mostZeroed)[1:191]),])
rowSums(sim.cleaned[c(names(mostZeroed)[1:191]),])
# as with melanogaster, some are highly expressed, some are very lowly expressed. No in-between. high
# expressed ones could be important markers that have a lot of background. eg. FBgn0045629/CG17575 - 
# very high expressed secondary cell marke that shows up as background in most main and ED cells as well.
FeaturePlot(sim.so, features = "FBgn0045629", pt.size=1) # CG17575, SC marker
FeaturePlot(sim.so, features = "FBgn0269608", pt.size=1) # ED marker, orthologous to Acp54A1 from tblastx
  #(not an official ortholog, not annotated)

# another type of gene mostly zeroedd are those that are kind of sporadically expressed, but at 
# relatively high levels on per cell basis 
FeaturePlot(sim.so, features = "FBgn0183433", pt.size=1)
FeaturePlot(sim.so, features = "FBgn0270050", pt.size=1)

### Visualizing SoupX changes in expression 
sim.umap <- sim.so@reductions$umap@cell.embeddings  # making a dataframe with UMAP coordinates

changemap_Dup99B <- plotChangeMap(sim.sc, sim.cleaned, DR = sim.umap, "FBgn0189236") + 
  scale_color_gradient(low = "#99D8C9", high = "#00441B") + 
  theme_minimal(base_size = 15) + 
  labs(x = "UMAP 1", y = "UMAP 2", color = "soup fraction", title = NULL) +
  geom_point(aes(col=relChange),size=2) 
changemap_Dup99B

changemap_SP <- plotChangeMap(sim.sc, sim.cleaned, DR = sim.umap, "FBgn0021129") + 
  scale_color_gradient(low = "#99D8C9", high = "#00441B") + 
  theme_minimal(base_size = 15) + 
  labs(x = "UMAP 1", y = "UMAP 2", color = "soup fraction", title = NULL) +
  geom_point(aes(col=relChange),size=2) 
changemap_SP

changemap_Spn77Bc <- plotChangeMap(sim.sc, sim.cleaned, DR = sim.umap, "FBgn0045603") + 
  scale_color_gradient(low = "#99D8C9", high = "#00441B") + 
  theme_minimal(base_size = 15) + 
  labs(x = "UMAP 1", y = "UMAP 2", color = "soup fraction", title = NULL) +
  geom_point(aes(col=relChange),size=2) 
changemap_Spn77Bc

changemap_Acp32CD <- plotChangeMap(sim.sc, sim.cleaned, DR = sim.umap, "FBgn0043403") + 
  scale_color_gradient(low = "#99D8C9", high = "#00441B") + 
  theme_minimal(base_size = 15) + 
  labs(x = "UMAP 1", y = "UMAP 2", color = "soup fraction", title = NULL) +
  geom_point(aes(col=relChange),size=2) 
changemap_Acp32CD

changemap_Acp36DE <- plotChangeMap(sim.sc, sim.cleaned, DR = sim.umap, "FBgn0041931") + 
  scale_color_gradient(low = "#99D8C9", high = "#00441B") + 
  theme_minimal(base_size = 15) + 
  labs(x = "UMAP 1", y = "UMAP 2", color = "soup fraction", title = NULL) +
  geom_point(aes(col=relChange),size=2) 
changemap_Acp36DE

# weird that in sim Acp36DE is a contaminant in ED cells but not really mel cells

# export the soup-corrected reads to a matrix
write.table(sim.cleaned, "sim.cleaned.tsv", sep = "\t", quote = F, col.names = NA)

################
##   YAKUBA   ##
################

###########################################################
### 1A. do preliminary clustering in for use with SoupX ###
###########################################################

yak.raw <- read.table("yak_cells_background.tsv", row.names = 1, header = T)
yak.so <- CreateSeuratObject(yak.raw, min.cells = 3, min.features = 100)

FeatureScatter(yak.so, feature1 = "nCount_RNA", feature2 = "nFeature_RNA", pt.size = 2)

# Show 5% qunatiles for number of genes per cell
do.call("cbind", tapply(yak.so@meta.data$nFeature_RNA,yak.so@active.ident,quantile,probs=seq(0,1,0.05)))
# Show 5% qunatiles for number of counts per cell per sample
do.call("cbind", tapply(yak.so@meta.data$nCount_RNA,yak.so@active.ident,quantile,probs=seq(0,1,0.05)))

yak.so <- subset(yak.so, nCount_RNA <= 2500)

# next normalize the data
yak.so <- NormalizeData(
  object = yak.so,
  normalization.method = "LogNormalize",
  scale.factor = 10000)

# FindVariableFeatures 

yak.so <- FindVariableFeatures(
  object = yak.so,
  selection.method = "vst")

# scale
yak.so <- ScaleData(object = yak.so)
# PCA
yak.so <- RunPCA(object = yak.so)
DimPlot(object = yak.so, dims = c(1,2), reduction = "pca")
DimPlot(object = yak.so, dims = c(1,3), reduction = "pca")
ElbowPlot(yak.so)

# JackStraw
yak.so <- JackStraw(
  object = yak.so, dims = 40)
yak.so <- ScoreJackStraw(yak.so, dims = 1:40)

JackStrawPlot(object = yak.so, dims = 1:40)
# observe plot to decide what PCs to use
use.pcs = c(1:9)

#UMAP

yak.so <- RunUMAP(yak.so, reduction = "pca", dims = use.pcs)
yak.so <- FindNeighbors(yak.so, reduction = "pca", dims = use.pcs)

# look at cluster nums at a series of resolutions
yak.so <- FindClusters(
  object = yak.so,
  resolution = seq(0.2,1.4,0.1),
  verbose = FALSE)
sapply(grep("res",colnames(yak.so@meta.data),value = TRUE),
       function(x) length(unique(yak.so@meta.data[,x])))
# choose resolution
Idents(yak.so) <- "RNA_snn_res.0.9"

# UMAP Visualization
DimPlot(yak.so, reduction = "umap", label = T, pt.size = 1)
FeaturePlot(yak.so, features = c('nCount_RNA'), pt.size=1)

# marker genes 
markers_yak_all <- FindAllMarkers(
  object = yak.so, 
  only.pos = TRUE, 
  min.pct = 0.25, 
  thresh.use = 0.25
)

dim(markers_yak_all)
head(markers_yak_all)

table(Idents(yak.so))
yak.so$CellType <- Idents(yak.so)

ggplot(data = yak.so@meta.data, aes(x = CellType, y = nCount_RNA)) +
  geom_violin(position = position_dodge(width = 0.9))  + 
  geom_jitter()

# heirarchical cluster tree
yak.so <- BuildClusterTree(
  yak.so, dims = use.pcs)

PlotClusterTree(yak.so)

FeaturePlot(yak.so, features = c("FBgn0228300", "FBgn0243166", "FBgn0235532"))
# FBgn0228300 = Dup99B
# FBgn0243166 = abd-A
# FBgn0235532 = Acp36DE

# merging clusters into known cell types, at least for the purpose of cleaning up with SoupX
## based on UMAP and heirarchical tree

yak.so.merged <- RenameIdents(
  object = yak.so,
  '1' = '0', '2' = '0', '3' = '0'
)

# naming the cell types based on marker genes
yak.so.merged <- RenameIdents(
  object = yak.so.merged,
  '0' = 'main cells', '5' = 'secondary cells', '4' = 'ejaculatory duct cells'
)
table(Idents(yak.so.merged))
#main cells        secondary cells ejaculatory duct cells 
#844                     48                     97 

yak.so.merged$CellType <- Idents(yak.so.merged)

ggplot(data = yak.so.merged@meta.data, aes(x = CellType, y = nCount_RNA)) +
  geom_violin(position = position_dodge(width = 0.9))  + 
  geom_jitter(size = 1) + 
  theme_minimal(base_size = 14)

FeatureScatter(yak.so, feature1 = "nCount_RNA", feature2 = "nFeature_RNA", pt.size = 2,) + 
  theme_minimal() +
  labs(title = NULL)

ggplot(data = yak.so.merged@meta.data, aes(x = CellType, y = nFeature_RNA/nCount_RNA)) +
  geom_violin(position = position_dodge(width = 0.9))  + 
  geom_jitter(size = 1) + 
  theme_minimal(base_size = 14)

#####################################
## 1B. ESTIMATE SOUP TRANSCRIPTOME ##
#####################################
# need to filter out the cells that I already treated as doublets from prelim clustering
good_cells <- rownames(yak.so.merged@meta.data)
empty_drops <- colnames(yak.raw[,1158:2157])
good_cells_plus_empties <- c(good_cells, empty_drops)
yak.filtered <- yak.raw[,colnames(yak.raw) %in% good_cells_plus_empties]

# create the soupchannel, estimate the soup transcriptome
yak.sc <- SoupChannel(tod = yak.filtered, toc = as.matrix(yak.filtered[,1:989]), keepDroplets = T, soupRange = c(0,100))

View(yak.sc$soupProfile)
# seems to check out. 

#################################
## 2. ESTIMATE SOUP PERCENTAGE ##
#################################

# check barcodes between seurat object and soup channel
head(yak.sc$metaData)
head(yak.so.merged@meta.data[,1:3])
tail(yak.sc$metaData)
tail(yak.so.merged@meta.data[,1:3])

# there is some slight decrease in nUMI (like a few UMIs) between soup channel and seurat object. 
# This is expected because in seurat I eliminated genes with < 3 UMI total from analysis

# add cell type to the metadata of seurat object
yak.so.merged$CellType <- Idents(yak.so.merged)

# add cell type data to soup channel
yak.sc <- setClusters(sc = yak.sc, clusters = yak.so.merged$CellType)

# look at markers as sanity check
quickMarkers(yak.sc$toc, clusters = yak.sc$metaData$clusters, N = 10)
# markers look correct, good.

# estimate contamination fraction
yak.sc <- autoEstCont(yak.sc, priorRho = 0.135) # basing my prior on the estimate of mel contamination %

################################
#### 3. REMOVE CONTAMINATION ###
################################  

yak.cleaned <- as.data.frame(adjustCounts(yak.sc))

### inspect the changes in expression

genecounts.raw = rowSums(yak.sc$toc > 0)
genecounts.filtered = rowSums(yak.cleaned > 0)
mostZeroed = tail(sort((genecounts.raw - genecounts.filtered)/genecounts.raw), n = 300)
mostZeroed # which genes are the most zeroed?

length(subset(mostZeroed, mostZeroed == 1.0)) #  75 genes zeroed in every cell

# genes zeroed in every cell: 
rowSums(yak.sc$toc[c(names(mostZeroed)[225:300]),])
# appear not to be highly expressed, probably not important markers

# genes zeroed in a fravtion of cells:
rowSums(yak.sc$toc[c(names(mostZeroed)[1:224]),])
rowSums(yak.cleaned[c(names(mostZeroed)[1:224]),])
# as with melanogaster, some are highly expressed, some are very lowly expressed. No in-between. high
# expressed ones could be important markers that have a lot of background. eg. FBgn0045629/CG17575 - 
# very high expressed secondary cell marke that shows up as background in most main and ED cells as well.
FeaturePlot(yak.so, features = "FBgn0276875", pt.size=1) # Dup99B
FeaturePlot(yak.so, features = "FBgn0236653", pt.size=1) # lectin 46A

# another type of gene mostly zeroedd are those that are kind of sporadically expressed, but at 
# relatively high levels on per cell basis 
FeaturePlot(yak.so, features = "FBgn0277014", pt.size=1) # ED marker with no orthologs
FeaturePlot(yak.so, features = "FBgn0275440", pt.size=1) # Met75Cb

### Visualizing SoupX changes in expression 
yak.umap <- yak.so@reductions$umap@cell.embeddings  # making a dataframe with UMAP coordinates

changemap_Dup99B <- plotChangeMap(yak.sc, yak.cleaned, DR = yak.umap, "FBgn0276875") + 
  scale_color_gradient(low = "#99D8C9", high = "#00441B") + 
  theme_minimal(base_size = 15) + 
  labs(x = "UMAP 1", y = "UMAP 2", color = "soup fraction", title = NULL) +
  geom_point(aes(col=relChange),size=2) 
changemap_Dup99B

changemap_SP <- plotChangeMap(yak.sc, yak.cleaned, DR = yak.umap, "FBgn0239194") + 
  scale_color_gradient(low = "#99D8C9", high = "#00441B") + 
  theme_minimal(base_size = 15) + 
  labs(x = "UMAP 1", y = "UMAP 2", color = "soup fraction", title = NULL) +
  geom_point(aes(col=relChange),size=2) 
changemap_SP

changemap_Spn77Bc <- plotChangeMap(yak.sc, yak.cleaned, DR = yak.umap, "FBgn0236999") + 
  scale_color_gradient(low = "#99D8C9", high = "#00441B") + 
  theme_minimal(base_size = 15) + 
  labs(x = "UMAP 1", y = "UMAP 2", color = "soup fraction", title = NULL) +
  geom_point(aes(col=relChange),size=2) 
changemap_Spn77Bc

changemap_Acp32CD <- plotChangeMap(yak.sc, yak.cleaned, DR = yak.umap, "FBgn0235918") + 
  scale_color_gradient(low = "#99D8C9", high = "#00441B") + 
  theme_minimal(base_size = 15) + 
  labs(x = "UMAP 1", y = "UMAP 2", color = "soup fraction", title = NULL) +
  geom_point(aes(col=relChange),size=2) 
changemap_Acp32CD

changemap_Acp36DE <- plotChangeMap(yak.sc, yak.cleaned, DR = yak.umap, "FBgn0235532") + 
  scale_color_gradient(low = "#99D8C9", high = "#00441B") + 
  theme_minimal(base_size = 15) + 
  labs(x = "UMAP 1", y = "UMAP 2", color = "soup fraction", title = NULL) +
  geom_point(aes(col=relChange),size=2) 
changemap_Acp36DE

# export the soup-corrected reads to a matrix
write.table(yak.cleaned, "yak.cleaned.tsv", sep = "\t", quote = F, col.names = NA)
```
